# A circular engineered sortase for interrogating histone H3 in chromatin

**DOI:** 10.1101/2024.09.10.612318

**Authors:** Samuel D. Whedon, Kwangwoon Lee, Zhipeng A. Wang, Emily Zahn, Congcong Lu, Maheeshi Yapa-Abeywardana, Louise Fairall, Eunju Nam, Sarah Dubois-Coyne, Pablo De Ioannes, Xinlei Sheng, Adelina Andrei, Emily Lundberg, Jennifer Jiang, Karim-Jean Armache, Yingming Zhao, John W. R. Schwabe, Mingxuan Wu, Benjamin A. Garcia, Philip A. Cole

## Abstract

Reversible modification of the histone H3 N-terminal tail is critical in regulating chromatin structure, gene expression, and cell states, while its dysregulation contributes to disease pathogenesis. Understanding the crosstalk between H3 tail modifications in nucleosomes constitutes a central challenge in epigenetics. Here we describe an engineered sortase transpeptidase, cW11, that displays highly favorable properties for introducing scarless H3 tails onto nucleosomes. This approach significantly accelerates the production of both symmetrically and asymmetrically modified nucleosomes. We demonstrate the utility of asymmetrically modified nucleosomes produced in this way in dissecting the impact of multiple modifications on eraser enzyme processing and molecular recognition by a reader protein. Moreover, we show that cW11 sortase is very effective at cutting and tagging histone H3 tails from endogenous histones, facilitating multiplex “cut-and-paste” middle down proteomics with tandem mass tags. This cut-and-paste proteomics approach permits the quantitative analysis of histone H3 modification crosstalk after treatment with different histone deacetylase inhibitors. We propose that these chemoenzymatic tail isolation and modification strategies made possible with cW11 sortase will broadly power epigenetics discovery and therapeutic development.

## Main

Understanding the patterns and functional interactions of histone tail posttranslational modifications (PTMs) has emerged as a central challenge in epigenetics.^1^ The building blocks of cellular chromatin are nucleosomes, which are comprised of a histone octamer (four pairs of histones H2A, H2B, H3, and H4) wrapped by 147 bp DNA. Conformational changes in chromatin contribute to the regulation of cell growth, differentiation, and gene expression.^2,3^ Such chromatin structural changes are influenced by reversible histone modifications which are inscribed by “writer” enzymes, removed by “eraser” enzymes, and functionally interpreted by “reader” domain proteins. Histone H3 N-terminal modifications are subjects of particular attention due to their central importance in gene regulation. Histone H3 N-terminal tail modifications include well-established Lys acetylation, methylation, and ubiquitination and lesser studied acylation modifications like propionylation, butyrylation, crotonylation, and succinylation.^4,5^ These and other PTMs are found in various combinations on histone H3 tails. This complex pattern of modifications affects chromatin structure and the writers, erasers, and readers that act on histones, but detailed molecular insights into how histone PTM crosstalk modulates these processes are generally lacking.^6,7^

In this study, we address two significant challenges in the field: limitations on the ready availability of designer nucleosomes and middle-down mass spectrometric analysis of histone H3 modifications. Although progress has been made over the past 15 years in the ability to prepare nucleosomes containing site-specific modifications on the H3 tail, this is still an onerous multistep task. This workflow requires individual expression and purification of all four histone proteins, including a truncated form of the modified histone, semisynthesis of the modified histone, octamer refolding, DNA isolation and finally nucleosome reconstitution.^8,9^ Starting from scratch this process takes about a month even in labs with experience. Moreover, substantial additional labor is required to produce nucleosomes containing distinct histone H3 tails, “asymmetric nucleosomes,” that can be employed to distinguish between biochemical effects mediated by modifications of H3 that occur on the same tails versus different tails.^10–13^

Middle-down mass spectrometric analysis of histone H3 can provide precise information about the interplay between modifications within individual histone H3 tails by evaluating intact H3 protein tails isolated from cellular histones.^14^ Current middle-down methods require purification of cellular histone H3 prior to treatment with the protease GluC.^15–17^ The resultant aa1-51 H3 peptide tails typically require complex and specialized chromatography for separation and characterization by tandem mass spectrometry, which is particularly challenging on such large peptide segments. Furthermore, this, like other middle-down proteomics approaches, lacks the quantitative strength of current tandem mass tag (TMT) isotopic labels, which are widely used in bottom-up mass spectrometry.^18^

Here, we address these limitations in histone H3 analysis through the application of a novel engineered sortase transpeptidase, cW11. The development of cW11 builds on the earlier work of Schwarzer and colleagues who reported F40 sortase as a tool for histone H3 semisynthesis.^19^ F40 sortase was developed as a chemoenzymatic tool to catalyze H3 tail attachment to tailless histone H3 tail recombinant protein which contains the sequence APXTG (aa29-33) rather than the natural sortase recognition epitope LPXTG.^19,20^ While useful, F40 sortase shows relatively slow rates of H3 tail attachment with standard amide bond sequence and relatively low yields of semisynthetic histone H3s. Sortase cW11 has been found to be a more effective catalyst of histone H3 transpeptidation reactions, facilitating H3 tail ligation to prefabricated tailless nucleosomes. Moreover, cW11 sortase can be employed to create asymmetric nucleosomes bearing distinct patterns of modifications on the H3 tails. The availability of such asymmetric nucleosomes has enabled new insights into molecular recognition of nucleosomes by eraser and reader proteins.^7,17^ In addition, efficient isolation of H3 peptide tails from crude extracts of endogenous histones with cW11 permits concurrent labeling with tandem mass tags. This ‘cut-and-paste’ approach enhances the middle-down proteomics analysis of histone H3 tails.

## Results

### Development of cW11 sortase

To improve on F40 sortase, mutant sortases containing amino acid replacements were designed based on structural considerations and previous studies on the development of enhanced sortase A (eSrtA).^20^ These mutations were introduced individually, then in combination, and mutant sortases were screened for transpeptidation-based cleavage of histone H3 in the presence of excess oligoglycine peptide. First pass screening employed purified histone H3 with no modifications (**Figure S1**). D165A, the single most activating mutation identified in eSrtA, falls within the helix mutagenized in F40, and generally hinders the cleavage reaction. Combined mutation of D160, K190 and K196 synergistically enhanced activity without disrupting the selectivity imparted by F40 mutations (T164 and V168-Q172). The best performing enzyme was tested against synthetic histone substrates with single modifications, and no significant bias was observed in the cleavage reaction (**Figure S2**).

With a sequence-selective sortase, sample handling steps in the typical middle-down proteomic workflow can be reduced (**Figure 1a**). Conducting the sortase reaction in a crude nuclear extract eliminates histone purification as a prerequisite for proteolytic isolation of histone tail (**Figure 1b**, **Figure S3**). Sortase performs optimally at neutral pH and low salt concentrations, however, this compromises the solubility of endogenous histones. Sortase exhibits minimal tolerance for detergents, co-solvents and chaotropes, which we sought to mitigate (**Figure S4**).^21^ Individual thermostabilizing mutations were identified using the FireProt web server and screened in combination (**Figure S4**).^22^ Sortase backbone cyclization was previously reported to enhance stability, and was also incorporated, leading to enzyme cW11.^21,22^ Sortase cW11 exhibits superior activity in nuclear acid extracts at physiological temperatures. The cleaved histone tail peptides are readily separated from sortase and other components of the nuclear acid extract reaction by trichloroacetic acid precipitation (**Figure 1c, Figure S5**). Following supernatant buffer exchange and cleanup with C18 stage tips, the peptides are suitable for LC-MS/MS analysis.

**Figure 1.**
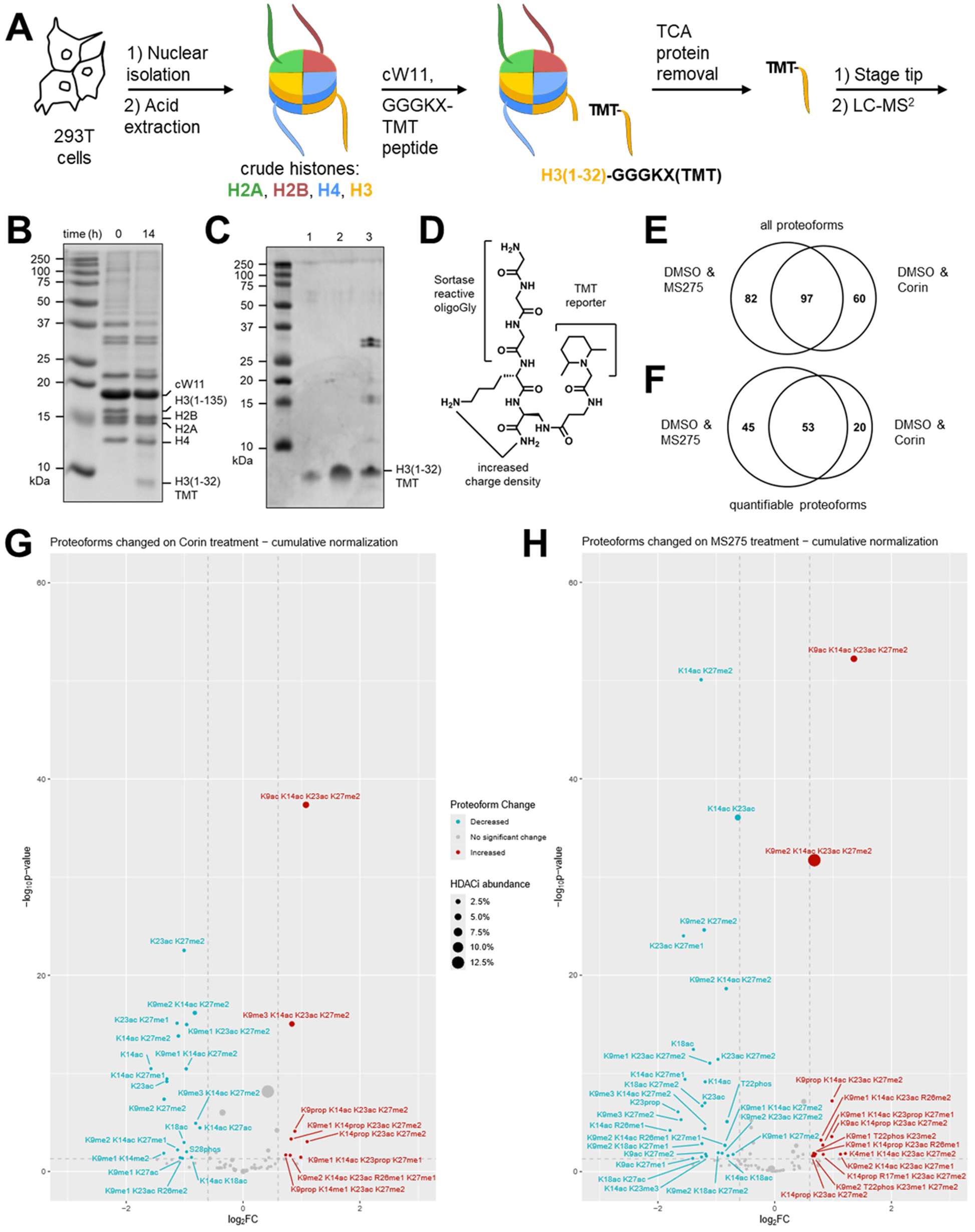
‘Cut-and-paste’ isolation of histone H3 tail peptides with cW11 sortase enables quantitative middle-down proteomics. (**A**) Workflow for isolating tandem mass tagged histone H3 tails with cW11 sortase. (**B**) SDS-PAGE of 14-hour sortase reaction in a nuclear acid extract. (**C**) Tris-Tricine PAGE of histone H3(1-34) synthetic standards (1: 140 ng; 2: 420 ng) and TCA precipitated H3(1-32)-TMT from the acid extract reaction; bands between 25 and 37 kDa are histone H1. (**D**) Structure of the TMT-labelled oligoglycine peptide illustrating sortase reactive GGG, charge carrying K and carboxamide, and TMT labelled aminoalanine. (**E**) Unique proteoforms detected in each 6plex sample and shared between the two samples. (**F**) Quantifiable proteoforms detected in each 6plex sample and shared between the two samples. Volcano plot of significant (p<0.05) log2-fold change and overall abundance of H3 proteoforms in HEK293T cells following treatment with (**G**) LSD1/HDAC1/CoREST complex-specific inhibitor Corin, or (**H**) pan class one HDAC inhibitor MS275.

### ‘Cut-and-paste’ middle-down proteomics of histone H3 modifications

In the sortase transpeptidation an N-terminal Gly is required for reaction with the enzyme-thioester intermediate, while C-terminal diversity is tolerated. By solid-phase peptide synthesis a GGGKX peptide was prepared with β-aminoalanine at the C-terminal position. After orthogonal deprotection of the side chain β-amino group the peptide resin was split into 6 portions, each of which was reacted with a unique TMT 6-plex NHS-ester reagent (**Figure 1d**). This peptide tag 6-plex system allowed concurrent analysis of histone tails from two cell treatment conditions, each in biological triplicate.

To explore cW11 sortase-mediated mass spectrometry analysis of H3 tails in a pharmacological application, we utilized this approach to evaluate the changes in the patterns of tail modifications in response to two different histone deacetylase (HDAC) inhibitors: MS275 and Corin.^23^ MS275 is a class I selective HDAC inhibitor, targeting HDACs 1-3 whereas Corin is a bivalent inhibitor that contains HDAC as well as an LSD1 demethylase targeting warhead. In this way, Corin appears to show selectivity toward CoREST-containing HDAC complexes. Treatment of HEK293T cells with these HDAC inhibitors or DMSO was performed for 6 hours, after which nuclear acid extracts were isolated. Samples from these experiments were divided for separate bottom-up and middle-down proteomic analyses. Histone H3 tails isolated for middle down were tandem mass tagged by cW11 sortase, with each replicate assigned a unique tandem mass tag. Quantification of individual H3 PTMs by GluC-based middle-down and bottom-up proteomics typically results in a Pearson correlation between 0.3 and 0.6.^24^ In this case, middle-down analysis of the sortase-derived H3 peptides resulted in a stronger correlation with bottom-up analysis of the same acid extracted histone samples (Pearson correlations of ∼0.8) (**Table S1**).

By LC-MS/MS we identified ∼150 uniquely modified peptides (proteoforms) in a combined DMSO vehicle / Corin sample, and ∼180 in a combined DMSO vehicle / MS275 sample (**Figure 1e, Table S2, S3**). Nearly half of the proteoforms identified had sufficient TMT ion signals for quantitation (≥2 per sample, **Figure 1f, Table S4, S5**). This quantifiable fraction accounted for ∼45-50% of all signal intensity attributable to H3 tail peptides. Relative to the DMSO vehicle, both HDAC inhibitors significantly decreased H3 tail proteoforms with one or two modifications, while increasing those with three or four modifications (**Figure 1g, h**). The most significant increase was seen in H3K9acK14acK23acK27me2, however, a greater increase was seen with MS275 than with Corin (155% v. 110% increase). It was hypothesized that H3K4me1/2 proteoforms would increase following Corin treatment due to LSD1 inhibition by Corin, however H3K4 modifications were detected infrequently in this analysis, consistent with prior middle-down analyses.^25^ H3K4me1K14acK23acK27me2 was the sole H3K4me proteoform to exhibit a significant change, an increase following MS275 treatment. This increase is in line with recent work pointing toward a general stimulatory effect of H3 acetylation on the MLL family K4 methyltransferases.^17^

### Streamlined production of modified nucleosomes

Histone octamers are typically prepared by the individual purification of each core histone protein followed by in vitro octamer assembly. Co-expression of the four core Xenopus histones in *E. coli* has been used to directly furnish octamer.^26^ In the course of this work, we attempted co-expression of the core histones with tailless histone H3 (aa33-135) and found that, in our hands, deletion of the H3 tail facilitated the isolation of intact histone octamer about 5-fold over comparable full-length H3 octamer (5 ± 2.5 mg/L culture) (**Figure S6**). This octamer could be readily wrapped by DNA (147 or 185 bp) to produce H3 tailless nucleosomes. These findings led us to pursue late-stage tail attachment to tailless nucleosomes by cW11 sortase. This would enable rapid designer nucleosome production from a common late-stage precursor and minimize the consumption of synthetic H3 tail peptide. Enhanced sortase variants eSrtA and 5M have been used to ligate histone H3 tail to nucleosomes and endogenous chromatin respectively but require an A29L mutation that complicates recognition of the heavily modified H3K27 site.^27,28^ F40 sortase is ineffective in this setting.

We investigated the activity of cW11 sortase in ligating H3 tails to tailless nucleosomes (aa33-135), and western blot revealed reasonable efficiency (∼90% ligation of H3) (**Figure S7**). Analytical anion exchange chromatography was used to precisely delineate the distribution of products, resolving nucleosomes with zero, one or two H3 tails ligated (**Figure S8**). This revealed a similar ligation efficiency with respect to histone H3 (79 ± 13%), with optimized conditions converting all tailless nucleosomes to products with either one or two H3 tails ligated. Nucleosomes with two ligated H3 tails typically accounted for 58 ± 6% of products (25 ± 3% isolated yield) (**Figure 2**). At our standard nanomole scale, 100 µg of nucleosome is obtained for every 100 µg of peptide, which is ∼100-fold less peptide than required to make the same amount of nucleosome by sortase semisynthesis of histone protein.^29^

**Figure 2.**
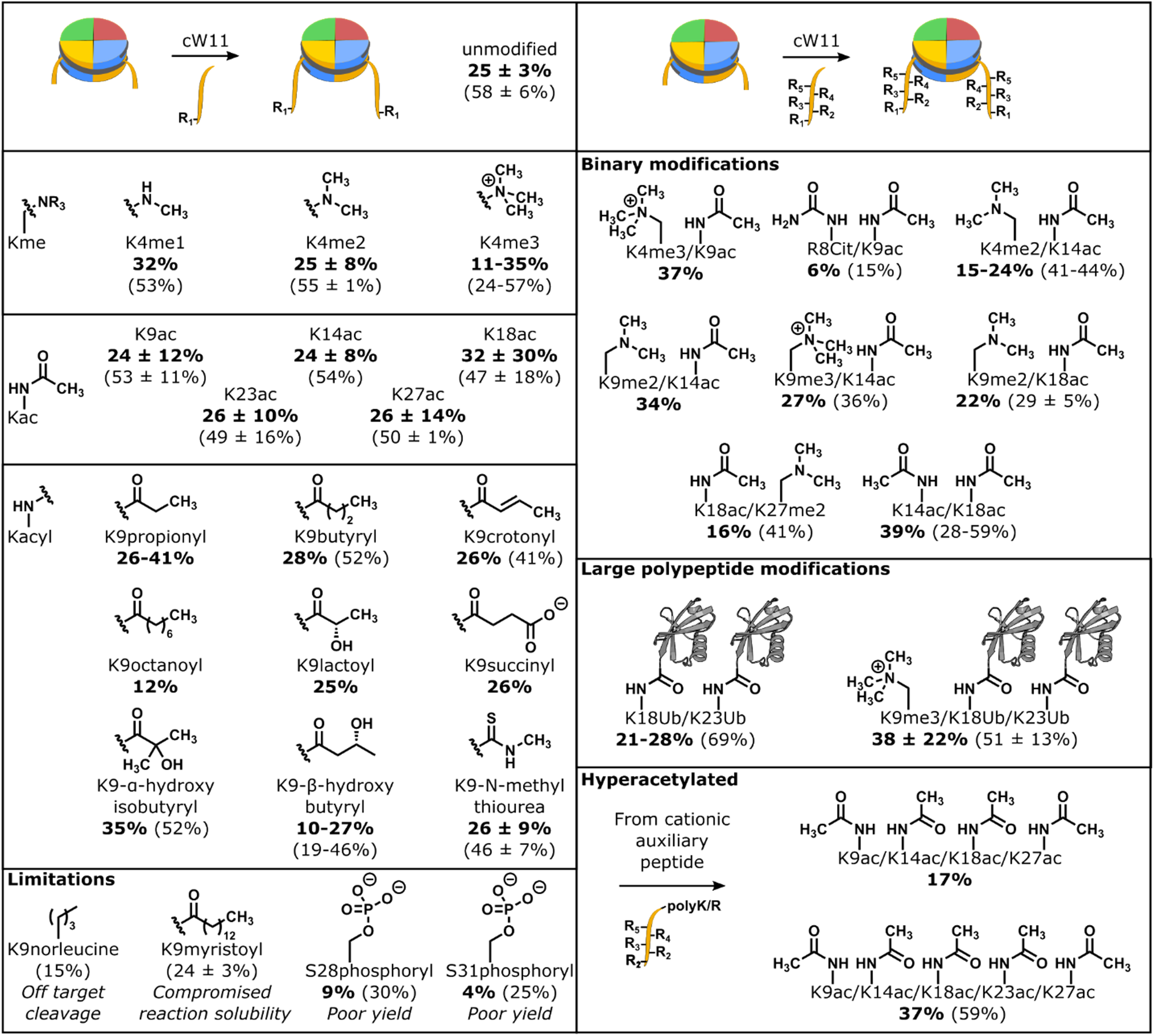
Designer nucleosome synthesis by sortase ligation. Single, combinatorial and asymmetric modifications prepared by sortase nucleosome ligation (ubiquitin structure PDBID: 1UBQ).^62^ Isolated yields (bold) and yields estimated by area under the curve (parenthetical) are reported as a value (n = 1), range (n = 2) or mean with standard deviation (n ≥ 3).

Late-stage H3 functionalization by cW11 ligation was evaluated for compatibility with modifications of lysine (acylations, alkylations, ubiquitination), arginine (citrullination) and serine/threonine (phosphorylation) (**Figure 2, Figure S9, S10**). Ligation of alkylated peptides resulted in yields similar to those of an unmodified peptide (LC: 53 ± 11%; isolated: 26 ± 9%), however, modifications decreasing the peptide positive charge were observed to decrease yield. For single acylations this could be overcome by substituting the Thr32 Gly33 amide linkage for a depsipeptide ester linkage.^19^ The first step in sortase transpeptidation, cleavage of the T-G amide bond, liberates a glycine dipeptide that competes with H3 in the transpeptidation reaction, however, the alcohol terminated byproduct of ester cleavage is incompatible with the reverse reaction. Irreversible cleavage of the synthetic peptide C-terminus proved invaluable in the synthesis of multiply acetylated nucleosomes, where the charge masking effect of the acetylations was offset by addition of multiple cationic residues after G34. Introduction of this cleavable C-terminal cationic auxiliary enabled ligation of peptides with five concurrent acetylations spanning all the major acetylation sites (K9, K14, K18, K23 and K27) (**Figure 2, Figure S11**).

Though broadly compatible with histone PTMs, there are limitations to the cW11 ligation. Introduction of a negative charge near the sorting motif (S28, S31) hinders ligation progress, resulting in isolable yields of 5-10%. Phosphorylation of S28 was found to have no effect on H3 tail cleavage during cW11 development (**Figure S2**). Thus, we hypothesize that the proximity of Gly33 to the DNA backbone (∼10-12Å, PDBID: A1OI) leads to repulsion between the DNA backbone and proximal histone tail peptide phosphorylation.^27,30^

Anion exchange chromatography resolves nucleosome ligation products from free DNA, and also separates nucleosomes with zero, one or two copies of H3(33-135). Isolation of nucleosomes with one copy of full length H3 and one copy of H3(33-135), followed by a second round of sortase ligation allows production of nucleosomes with asymmetric modifications of the H3 tail (**Figure 3**, **Figure S12**). Asymmetric, or heterotypic, nucleosomes are sought after tools for unraveling crosstalk between modifications of the histone tails, but current strategies for their production are cumbersome.^11^ The cW11 ligation simplifies their production, requiring only minor changes in peptide stoichiometry to favor production of the intermediate single tail nucleosome and subsequent asymmetric nucleosome.

**Figure 3.**
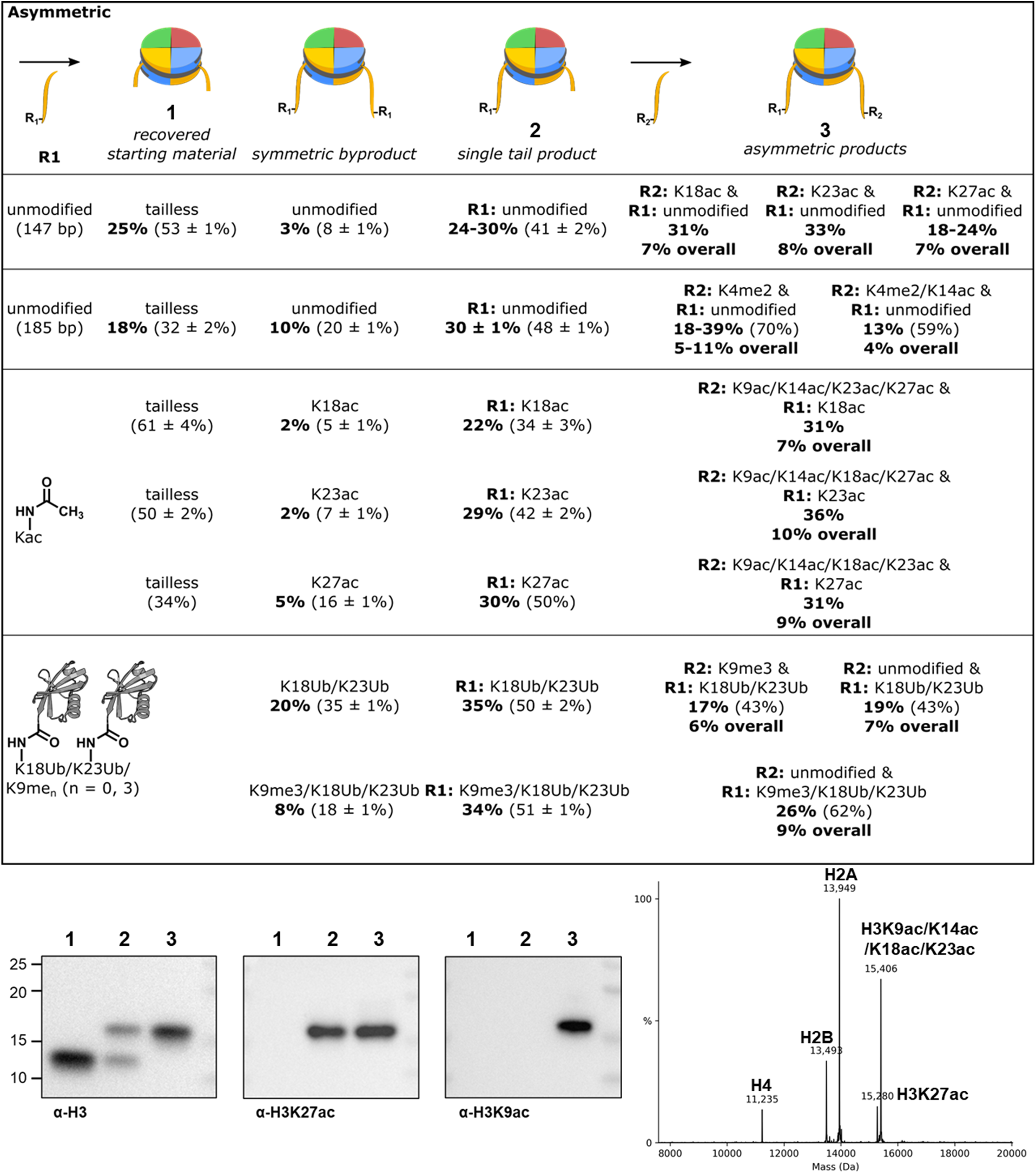
Asymmetric nucleosome synthesis by sortase ligation. (Top) Asymmetric modifications prepared by sortase nucleosome ligation. Isolated yields (bold) and yields estimated by area under the curve (parenthetical) are reported as a value (n = 1), range (n = 2) or mean with standard deviation (n ≥ 3). (Bottom left) Western blot characterization of 147 bp nucleosomes: (**1**) H3 (aa33-135) starting material; (**2**) asymmetric H3K27ac & H3 (aa33-135) intermediate; (**3**) asymmetric H3K27ac & H3K9ac/K14ac/K18ac/K23ac product. (Bottom right) Deconvoluted mass spectrum of 147 bp asymmetric H3K27ac & H3K9ac/K14ac/K18ac/K23ac nucleosome.

### Confirmation of nucleosome biochemical integrity

We have previously reported site-specific nucleosome deacetylation rates for multiple enzymes and complexes at most of the major acetylation sites on histone H3.^29,31,32^ In theory, our new approach to preparing nucleosomes by late-stage tail addition could lead to unintended consequences in biochemical analysis. For example, residual peptide could be present, which might compete with nucleosome as an enzyme substrate and alter nucleosome recognition by masking DNA, either of which was predicted to affect deacetylase rate. As a quality control for the cW11 ligation, we conducted parallel deacetylation rate studies using nucleosome substrates prepared by our new approach and using previously established methods. Using those HDACs most extensively characterized in our lab we found no significant difference in deacetylation rates (**Figure S13, S14**).^33,34^ Finally, cryo-EM was used to validate nucleosome structure and integrity, and resulted in a structure consistent with numerous other 147 bp nucleosome structures (**Figure S15**).

### Revealing histone deacylase activity with acylated nucleosomes

Reports of histone lysine succinylation, propionylation and butyrylation, among others have steadily accumulated in recent years, while characterization of their erasers has lagged. Broad functional group compatibility in the sortase ligation initiated characterization of histone deacylase activity toward these variant acylations. Drawing upon previously reported histone acylation selectivity and proposed modes of nucleosome-recognition, we selected the nuclear or nucleocytoplasmic HDACs mitosis-associated HDAC complex (MiDAC) and LSD1/HDAC/CoREST (LHC) complex as representatives of class I HDAC complexes, and Sirtuins 1, 2, and 6 as representatives of class III HDACs with which to explore the class- or complex-specific modes of deacylation.^29,33,35,36^

Across all sites on the H3 tail K9 is generally the most rapidly deacetylated, and was therefore selected to probe reactivity trends. Removal of acetyl, propionyl, butyryl and octanoyl modifications were compared (**Figure 4a**, **Figure S16-S19**), while myristoylation proved intractable in the sortase nucleosome ligation (**Figure 3**). Sirtuins 2 and 6 were superior long chain deacylases processing K9octanoyl substrates 18-fold faster than K9ac and ∼5-fold faster than K9ac respectively. Sirtuin 1 exhibited no activity toward any acylation, consistent with prior observations. Long chain deacylation by the class I HDACs is not much evidenced, so MiDAC was selected for comparison as its baseline deacetylation of K9 is the fastest measured by an order of magnitude. Deacylation by MiDAC slowed with chain length, decreasing ∼180-fold (K9butyryl) and ∼16-fold (K9octanoyl) relative to K9ac. That octanoylation was processed at all is chemically interesting, and particularly that MiDAC favors it ∼10-fold over butyrylation (**Figure 4a**, **Table S6**).

**Figure 4.**
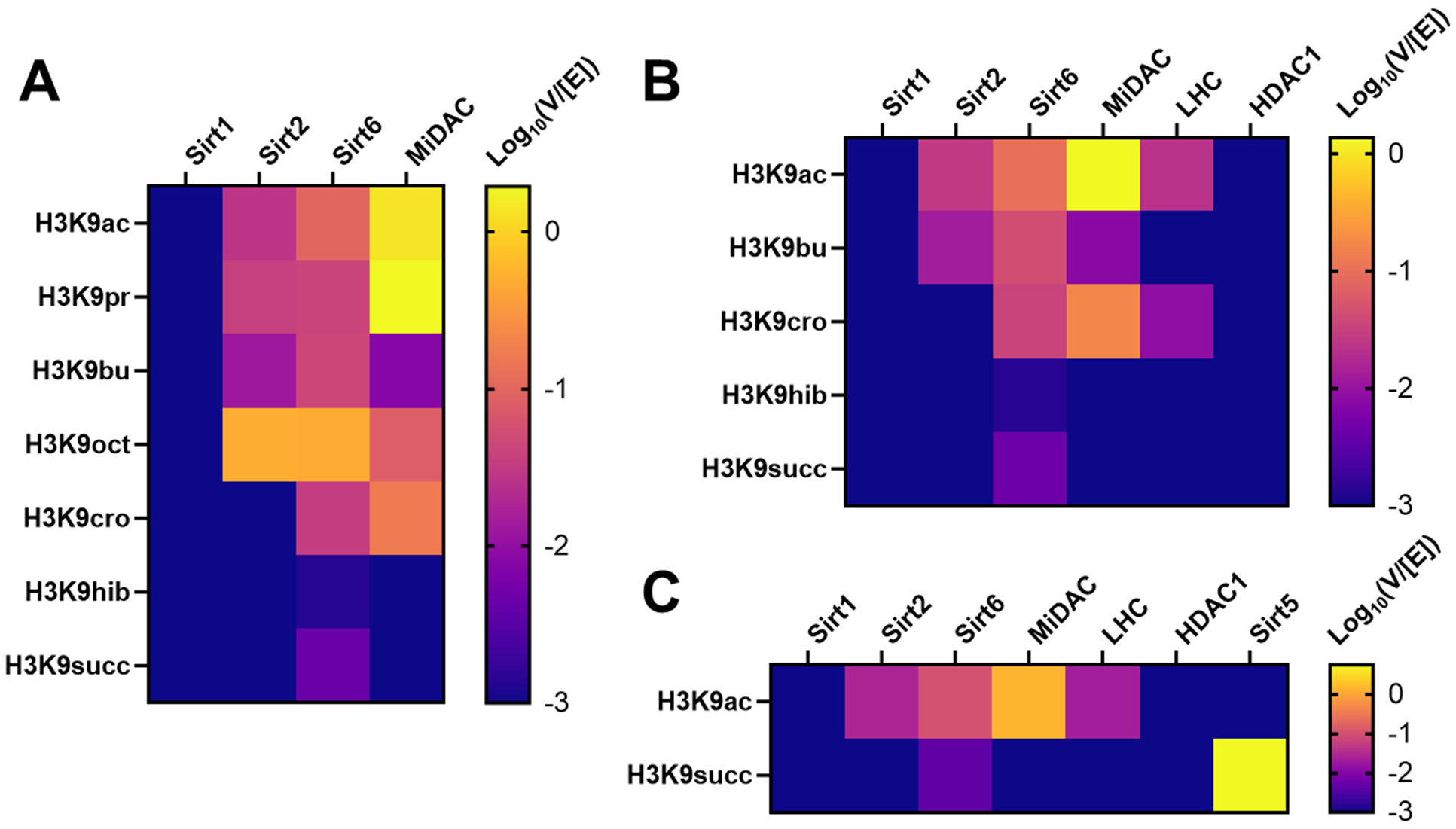
Trends in histone deacylase activity toward metabolic acylations. (**A**) Log10 transformed V/[E] (min^-1^) of Sirtuins 1, 2 and 6 and MiDAC toward 147 bp nucleosomes with one to eight carbon acylations of H3K9. (**B**) Log10 transformed V/[E] (min^-1^) of class one and class three HDACs toward 147 bp nucleosomes with four carbon metabolism-linked acylations of H3K9. (**C**) Log10 transformed V/[E] (min^-1^) of class one and class three HDACs toward 147 bp nucleosomes with H3K9 succinylation.

For each deacylase enzyme with detectable activity, butyrylated nucleosomes were consistently the slow substrate, perhaps accounting for its observed metabolic accumulation in specific tissue types.^37^ However, sirtuin catalysis is reported to be enhanced by long chain acylations that pack within the active site. Exploring the conformational contribution to substrate selectivity for deacylases, four carbon acylations with links to metabolic state were tested, including crotonylation, hydroxyisobutyrylation and succinylation (**Figure 4b, Figure S20-S25**). Of these decrotonylation exhibited a rate similar to debutyrylation for Sirt6, and faster than debutyrylation for MiDAC and LHC (**Table S7**). Removal of the branched and charged modifications was unmeasurable for all but Sirt6, which was exceptionally slow. Succinylation is the preferred substrate of Sirt5, which desuccinylated nucleosomes at the fastest rate measured among all deacylases and acylations surveyed here (V/[E] = 5.7 min^-1^) (**Figure 4c, Figure S26, Table S8**).

### Symmetric and asymmetric nucleosome tools for testing consequences of histone hyperacetylation

Among the synthetic nucleosomes prepared were those with five acetylations of H3 at K9, K14, K18, K23, and K27, which are the predominant sites of H3 acetylation detected in the sortase middle-down data, and reported by others. The sharp rise in multiply acetylated peptides following HDAC inhibition, and prior reports of histone acetyltransferase ‘acetyl spray’ activity, prompted investigation of these hyperacetylated substrates.^38^ Extensive acetylation increases histone tail dynamic motion and alters the local electrostatic environment encountered by regulators like HDACs.^39–41^ Whether this influences the rate of site-specific deacylation was tested using a K9ac/K14ac/K18ac/K23ac/K27ac penta-acetylated substrate, and site-specific rates were compared to previously reported rates for each of K9ac, K14ac, K18ac, K23ac, and K27ac.^31,32^

The rate of deacetylation of penta-acetylated nucleosomes by different enzymes was monitored by western blot with site-selective anti-acetyl-Lys antibodies, the site-specificity of which was validated (**Figure S27**) with our designer nucleosomes (**Figure 5a-c, Figure S28-S31**). Penta-acetylation influenced removal for a select set of acetyl-Lys locations with distinct impacts among the different enzymes. Deacetylation of H3K27 by both Sirt2 and Sirt6 slowed ∼2-fold, while deacetylation by MiDAC increased nearly 4-fold. Sirt2 exhibited a striking change in site-selectivity from K27 to K18, due to ∼2-fold deacetylation rate increases at K18 and K23. To try to better understand the mechanistic basis for these effects, we used the sequential ligations enabled by cW11 sortase to prepare asymmetrically modified nucleosome substrates.

**Figure 5.**
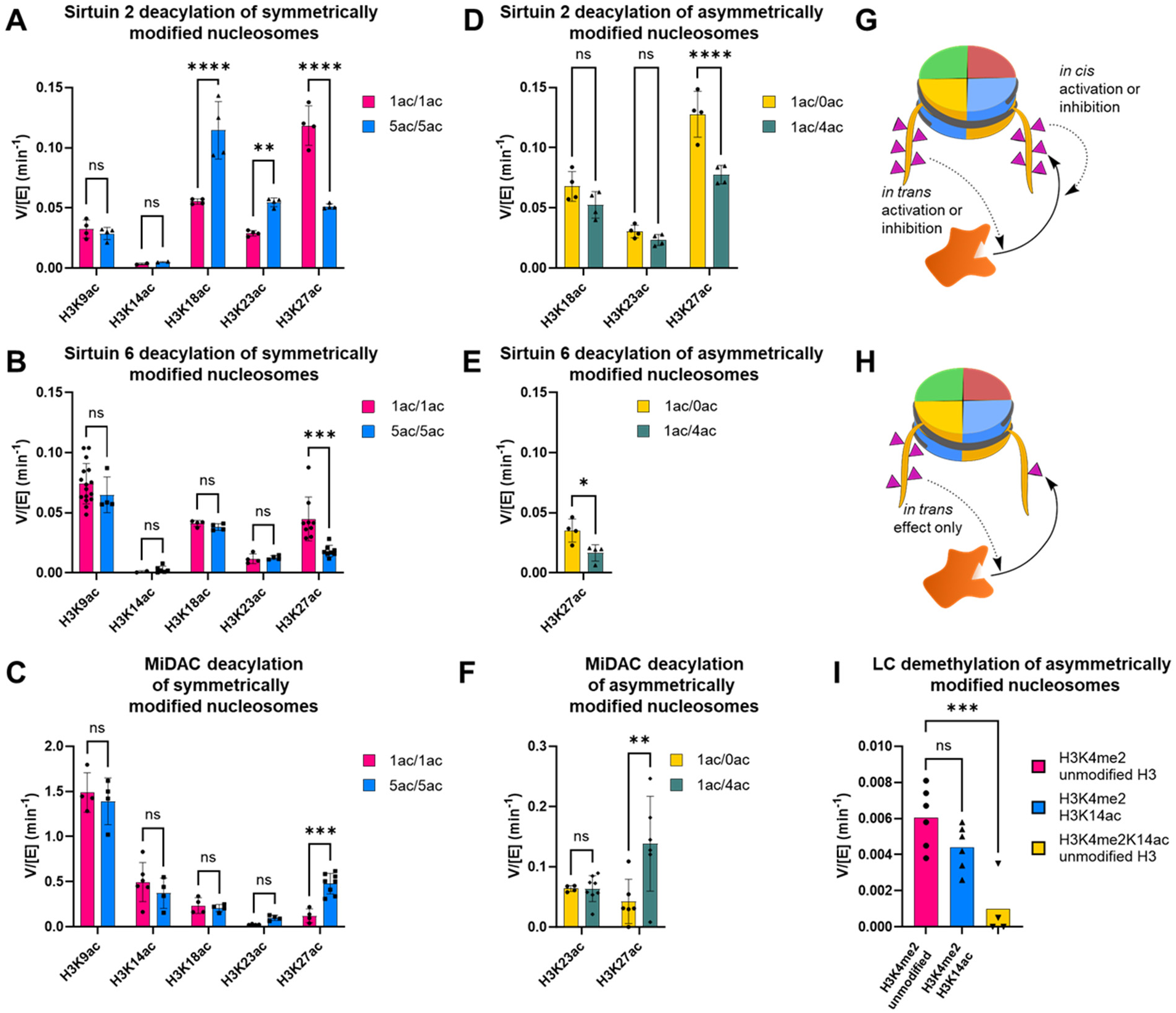
Symmetric and Asymmetric effects of hyperacetylation on deacylase and demethylase activity. (**A**) Sirt2, (**B**) Sirt6, and (**C**) MiDAC site-specific deacetylation rates with symmetrical mono-acetylated (1ac/1ac) and penta-acetylated (5ac/5ac) 147 bp nucleosomes. (**D**) Sirt2, (**E**) Sirt6, and (**F**) MiDAC site-specific deacetylation rates with asymmetric 147 bp nucleosomes isolating the specified acetylation site on a single tail; the second H3 tail was modified with either four acetylations (1ac/4ac) at the other predominant acetylation sites, or zero acetylations (1ac/0ac). (**G**) Illustration of *in cis* and *in trans* PTM interactions with a regulatory enzyme acting on a nucleosome substrate. (**H**) Illustration of an asymmetric nucleosome used to test for an *in trans* effect on a PTM regulatory enzyme. (**I**) LSD1-CoREST1 (LC) demethylation rates with asymmetric nucleosomes containing H3K14ac concurrently on the same histone H3 with H3K4me2 (yellow), H3K4me2 and H3K14ac separately on different histone H3 (blue), and without H3K14ac (pink) (* indicates p < 0.05, ** indicates p < 0.01,*** indicates p < 0.001, **** indicates p < 0.0001).

For each site exhibiting a rate changed by penta-acetylation, an asymmetrically modified nucleosome was prepared to isolate that site from the other four acetylations. For example, all three enzymes surveyed exhibited altered rates of deacetylation at K27, and the nature of those rate changes could be assayed using nucleosomes with one copy of H3K27ac and one copy of H3K9ac/K14ac/K18ac/K23ac. Compared to the symmetrically acetylated substrates assayed previously, this substrate has half the effective concentration of K27ac, one copy per nucleosome (**Figure S32, S33**). To control for this, rate comparisons were made using asymmetric nucleosomes with one unmodified copy of H3 and one copy of H3K27ac (**Figure S33**). Deacetylation of asymmetric K9ac/K14ac/K18ac/K23ac and K27ac nucleosomes by Sirt2 and Sirt6 was ∼2-fold slower than the matched control (**Figure 5d, e, Figure S34, S35, Table S9, S10**). This is consistent with competition between acetylation sites driving the rate decrease, rather than a local acetylation slowing the rate through altered enzyme recognition. With the same substrates MiDAC showed ∼3-fold faster deacetylation of the more acetylated asymmetric substrate than the singly acetylated asymmetric substrate (**Figure 5f, Figure S36, Table S11**). This *in trans* stimulation (**Figure 5g, h**) suggests a processive mechanism at the level of the nucleosome rather than a single histone tail and suggests that histone tail mobility alone does not explain accelerated deacetylation of K27 by MiDAC in a hyperacetylated context. Notably, MiDAC exists as a predominantly tetrameric complex with four catalytic HDAC1/2 modules and the dimeric MiDAC subunit DNTTIP1 is a nucleosome acidic patch binder.^35,42^ We speculate that multivalent binding through these domains could drive crosstalk between H3 tails.

Deacetylation of K18 and K23 by Sirt2 was accelerated in the penta-acetylated context, prompting assays in which each site was isolated individually (**Figure 5d**). In each case, whether the opposite H3 tail had zero or four acetylations, the deacetylation rate was insignificantly different. Here, accelerated deacetylation depends on *in cis* modification of one H3 tail with multiple acetylations.

### Symmetric and asymmetric nucleosomes as tools for unraveling methylation-acetylation crosstalk

We have previously reported that demethylation of H3K4me1/2 by LC or LHC slows >5-fold in the presence of H3K14ac specifically.^43^ Recent crystallographic evidence points at a candidate *in cis* mechanism but cannot rule out *in trans* inhibition.^34^ Final resolution of the inhibitory effect was attainable using a series of asymmetrically modified nucleosome substrates in which K4me2 and K14ac modifications could be sequentially isolated. Demethylation rates for nucleosomes with H3K4me2 and one of either unmodified H3 or H3K14ac were insignificantly different (**Figure 5i, Figure S37**). To control for absolute PTM concentration these were both compared to asymmetric nucleosomes with H3K4me2/K14ac and unmodified H3, which again exhibited >5-fold reduction in demethylase activity by LC.

### DNMT1 recognition of symmetric versus asymmetric nucleosomes

Multi-monoubiquitination of histone H3 has been reported to direct binding by DNMT1, as has H3K9me3. The contribution of each modification has been partially evaluated using peptide, protein, and recently nucleosome substrates.^44–46^ The replication foci targeting sequence domain (RFTS) of DNMT1 has an atypical H3K9me3 binding site, but whether this exclusively reads methylation *in cis* was untestable in prior investigations. To address this, peptides, and subsequently asymmetric nucleosomes were prepared with unmodified H3 and either K18Ub/K23Ub or K9me3/K18Ub/K23Ub, as well as H3K9me3 and K18Ub/K23Ub (**Figure S38, S39**). An electrophoretic mobility shift assay (EMSA) was used to assess differences in binding of each substrate with a fluorescent RFTS domain fusion construct. The combination of K9me3/K18Ub/K23Ub nucleosome and RFTS domain resulted in a clearly resolved band shift at single digit nanomolar concentrations (**Figure 6**, left). The other two nucleosomes produced diffuse and smeared shifts of the RFTS band, and consistently showed a weak signal for the complex band (**Figure 6**, center, right). This was observed with either component (RFTS or nucleosome) used as titrant, and appeared constant over time (**Figure S40, S41**). Based on this EMSA behavior we deduce that the *in cis* K9me3/K18Ub/K23Ub modifications confer substantially enhanced stability of the RFTS-nucleosome interaction, which are not maintained when K9me3 is present *in trans*.

**Figure 6.**
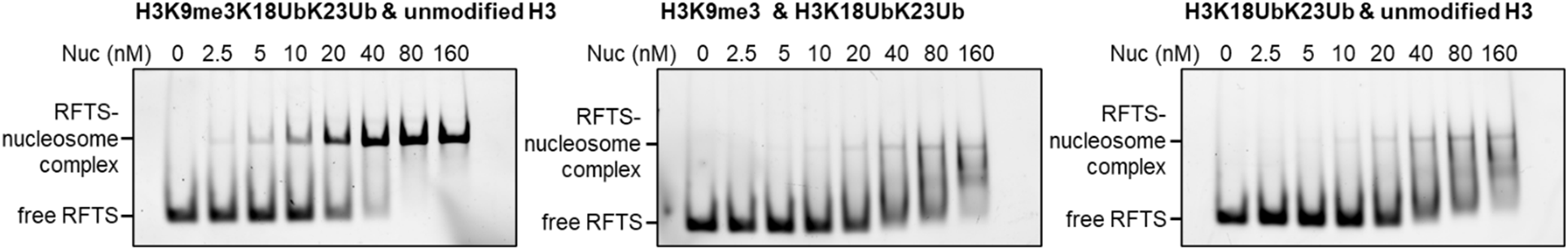
Combinatorial effect of H3K9me3 and K18Ub/K23Ub on DNMT1 RFTS domain binding. RFTS-sfGFP EMSA following titration with asymmetric H3K9me3/K18Ub/K23Ub & unmodified H3 nucleosome (left), asymmetric H3K18Ub/K23Ub & H3K9me3 nucleosome (middle), asymmetric H3K18Ub/K23Ub & unmodified H3 nucleosome (right).

## Discussion

Here we identify a powerful transpeptidase, cW11 sortase, that can efficiently ligate and cleave N-terminal H3 tails from histone H3. This has facilitated three important methods in chromatin analysis. First, cW11 sortase can be used to simply and rapidly produce site-specifically modified nucleosomes by permitting late-stage attachment of H3 tails to H3 tailless nucleosomes. An unexpected dividend that accelerates this approach is that the H3 tailless histone octamer can be efficiently generated by co-expressing tailless H3 with the other three core histones. This not only reduces the time needed to generate designer nucleosomes from about one month to one week, this approach dramatically reduces synthetic tail peptide amounts needed for ligation. We have shown that this cW11 sortase-driven strategy can easily incorporate a wide range of single and multiple PTMs (acetyl, acyl, methyl) into nucleosomes. This is especially useful for ubiquitin-like modifications, which present a formidable synthetic challenge, particularly when introduced along with other PTMs.^47–49^ Streamlining such syntheses facilitates analysis of these nucleosomes in enzymatic and binding experiments.^46,50^ These studies have uncovered interesting new selectivities of deacylases that interconnect metabolism and chromatin structure. Of particular note, both class I and class IV HDACs remove acylations with linear chains up to eight carbons, however, branched or polar acylations are exclusively processed by specific sirtuins.

Second, cW11 sortase provides an attractive and reliable route to asymmetric nucleosomes in which the two different H3 tails contain distinct patterns of site-specific PTMs. Prior work on the construction of asymmetric nucleosomes with PTMs on histones H2A, H2B, and H3 have been reported but are technically demanding and not widely used.^11,12^ Our simple stepwise addition of H3 tails with a single chromatographic isolation of the intermediate single tail form represents a relatively convenient alternative. Using this approach, we examined in asymmetric nucleosomes the crosstalk between H3K4me2 and H3K14ac by LSD1 demethylase, multi-acetylated H3 tails by various histone deacetylases, and Lys ubiquitination and methylation by the RFTS reader domain of DNMT1. These experiments have revealed cases where specific patterns can impact eraser or reader interactions specifically in *cis* or *trans* depending on the site of modification and eraser/reader involved.^7,17^ While the precise mechanisms for these PTM crosstalk influences remain to be elucidated, these multifaceted regulatory features provide interesting and novel insights into histone mark crosstalk and highlight the necessity of studying asymmetric nucleosomes to understand molecular recognition of individual chromatin interactors.

Third, we have adopted cW11 sortase to perform a “cut-and-paste” method to isolate and analyze purified intact histone H3 tails from a human cell line. Though the position of the sortase recognition motif precludes quantification of H3K36 modifications, this procedure imparts quantitative power to LC-MS/MS analysis of these tails through multiplex tandem mass tagging. With this approach, we were able to discern some overlapping but also distinct H3 tail PTM patterns after treatment with two types of HDAC inhibitors. We believe that this method can be generally useful for analyzing other epigenetic agents and to analyze specific cell states. The use of the chemoenzymatic labeling step with isotopic barcodes avoids the more complicated electrophilic chemical tandem mass tagging which is avoided on large peptides because of complex reactivities. We propose that the use of engineered sortase tagging may be broadly useful in middle-down proteomics.

## Methods

### Protein expression and purification

Plasmids for the relevant proteins and nucleic acids were transformed into *E. coli*, or co-transformed in the case of octamer, then selected with the relevant antibiotics, and grown from cell stocks. Sortase cW11 (from LOBSTR Rosetta *E. coli*) was grown at 37°C from an overnight starter culture of Luria-Bertani (LB) medium (Sigma) with ampicillin (100 mg/L), of which 10 mL was used to inoculate each 1 L of the same medium used for overexpression. Cells were grown to an OD600 of 0.6-0.8, then induced with 0.5 mM IPTG for 3 hours at 37 °C. Cells were harvested by centrifugation (4 k x g, 30 min, 4°C), and the cell pellet was resuspended in 5 volumes of chilled (4 °C) lysis buffer (10 mM Tris, pH 7.5 at 25 C, 0.1 % Tween-20). Resuspended pellets were mixed on a dounce homogenizer to uniformity and lysed by three passages through a microfluidizer. The supernatant was cleared by centrifugation (12 k x g, 30 min, 4 °C) and the supernatant was passed over 5 mL Ni NTA agarose twice by gravity. Resin was washed with 10 column volumes (CV) lysis buffer, followed by 10 CV wash buffer (10 mM Tris, pH 7.5 at 25 °C, 500 mM NaCl), then 5 CV imidazole wash buffer (10 mM Tris, pH 7.5 at 25 °C, 0.1 % Tween-20, 10 mM imidazole), and eluted with 5 CV elution buffer (10 mM Tris, pH 7.5 at 25 °C, 400 mM imidazole). The elution was dialyzed three times (Spectra/Por 12-14 kDa MWCO membrane, Spectrum Labs, 50 mM Tris, pH 7.5 at 25 °C, 150 mM NaCl, 5 mM CaCl_2_) at 4 °C, then concentrated (Amicon, 10 kDa MWCO, 4 °C) to 2-6 mM with frequent sample mixing to prevent aggregation.

Histone octamer (from LOBSTR Rosetta *E. coli*) was grown from an overnight starter culture (30 °C) of LB medium (Sigma) with ampicillin (100 mg/L), kanamycin (50 mg/L), streptomycin (100 mg/L) and chloramphenicol (34 mg/L), which was not allowed to reach saturation. Growth for overexpression was initiated by adding 10 mL of starter culture to inoculate each 1 L of the same medium. Octamer growth is typically slow, taking up to 12 hours to reach the target induction OD600 of 0.8. Expression was induced by adding IPTG to 0.5 mM, and cultures were grown overnight (12-16 h) at 25°C. Cells were harvested by centrifugation (4 k x g, 30 min, 4 °C), and the cell pellet was resuspended in 5 volumes of chilled (4 °C) lysis buffer (20 mM Tris, pH 7.5 at 25 °C, 2 M NaCl, 0.5 mM TCEP). Resuspended pellets were dounced to uniformity and lysed by three passages through a microfluidizer. Supernatant was cleared by centrifugation (12 k x g, 30 min, 4 °C) and the supernatant was batch bound to 10 mL Ni NTA agarose with gentle agitation at 4 °C. Resin was pelleted by centrifugation (500 x g, 5 min, 4 °C), and the supernatant was removed. Resin was washed with 5 CV lysis buffer, followed by 5 CV wash buffer (20 mM Tris, pH 7.5 at 25 °C, 2 M NaCl, 0.5 mM TCEP, 20 mM imidazole), then eluted with 5 CV elution buffer (20 mM Tris, pH 7.5 at 25 C, 2 M NaCl, 0.5 mM TCEP, 200 mM imidazole). The elution was concentrated (Amicon, 10 kDa MWCO, 4 °C) to 5-10 mg/mL by 280 nm absorbance (nanodrop), and incubated overnight with 0.01 mass equivalents of TEV protease at 4 °C. Octamer was further purified on an FPLC using a superdex 200 column (20 mM Tris, pH 7.5 at 25 °C, 2 M NaCl, 0.5 mM TCEP), then purified fractions were pooled and concentrated to 5-10 mg/mL. Glycerol and NaCl (5 M) were added to final concentrations of 15% and 2 M respectively, the samples mixed thoroughly, and aliquoted before flash freezing for storage at -80 °C.

Widom 601 DNAs (147 and 185 bp) were produced from multi-copy plasmids (16 x 147, 12 x 185) isolated from transformed DH5α *E. coli* grown for 36-48 h in CircleGrow media supplemented with antibiotic (147 bp: Ampicillin; 185 bp: Kanamycin). Cells were harvested by centrifugation (4 k x g, 30 min, 4 °C), and the cell pellet was resuspended in 5 volumes of lysis buffer (10 mM Tris, pH 8 at 25 °C, 10 mM EDTA). Lysis was initiated with lysozyme, after which 0.8 volumes of alkaline lysis buffer (2% SDS, 0.2 M NaOH) was added and the sample was mixed by inverting. One volume of neutralization buffer (1.5 M potassium acetate / acetic acid (pH 4.9)) was added and the sample was mixed by inverting, then insoluble material was removed by centrifugation. Clarified supernatant was filtered, then crude product was precipitated with isopropanol (0.6 volumes). Crude product was resuspended in 10 mM Tris (pH 8), 10 mM EDTA, and one volume of 5 M LiCl was added to precipitate RNA and proteins. The solution was clarified by centrifugation and the supernatant was combined with isopropanol (0.6 volumes) to precipitate crude product. The crude product was again resuspended in 10 mM Tris (pH 8), 1 mM EDTA and treated with RNase A, before precipitating again with isopropanol (3 volumes). The pelleted plasmid was resuspended in 10 mM Tris (pH 8), 1 mM EDTA, and digested with EcoRV in CutSmart buffer (New England Biolabs) at 37 °C. Insert DNA was purified by diluting the reaction 10-fold with dilution buffer (10 mM Tris, pH 8 at 25 °C, 1 mM EDTA) and passing twice over Sepharose fast-flow Q resin. The resin was washed with 20 volumes of wash buffer (10 mM Tris, pH 8 at 25 °C, 300 mM NaCl, 1 mM EDTA) and eluted with 5 volumes of elution buffer (10 mM Tris, pH 8 at 25 °C, 700 mM NaCl, 1 mM EDTA) then concentrated and stored at -20 °C.

C-terminally truncated ubiquitin (aa1-75) was obtained by expression of a C-terminal Ava intein fusion construct (A kind gift from Dr. Champak Chatterjee, University of Washington) from BL21(DE3) *E. coli*. An overnight starter culture of Luria-Bertani medium (Sigma) with ampicillin (100 mg/L) was grown overnight at 37 °C, of which 10 mL was used to inoculate each 1 L of the same medium used for overexpression. Cells were grown to an OD600 of 0.6-0.8, then chilled on ice and induced with 0.3 mM IPTG for 16 hours at 16 °C. Cells were harvested by centrifugation (4 k x g, 30 min, 4 °C), and the cell pellet was resuspended in 5 volumes of chilled (4 °C) lysis buffer (50 mM sodium phosphate, pH 8.0, 300 mM NaCl, 5 mM imidazole, 1 mM beta-mercaptoethanol). Resuspended pellets were dounced to uniformity and lysed by three passages through a microfluidizer. The supernatant was cleared by centrifugation (12 k x g, 30 min, 4 °C) and the supernatant was passed over 5 mL Ni NTA agarose twice by gravity. Resin was washed with 20 CV lysis buffer, then washed with 20 CV wash buffer (50 mM sodium phosphate, pH 8.0, 300 mM NaCl, 20 mM imidazole, 1 mM beta-mercaptoethanol), and eluted with 6 CV of elution buffer (50 mM sodium phosphate, pH 8.0, 300 mM NaCl, 250 mM imidazole, 1 mM beta-mercaptoethanol). Elution fractions were pooled and dialyzed three times against dialysis buffer (100 mM sodium phosphate, pH 7.2, 150 mM NaCl, 1 mM EDTA, 1 mM sodium 2-mercaptoethanesuflonate) at 4 °C. Thiolysis was initiated by recovering the dialyzed solution, bringing it to 100 mM in sodium 2-mercaptoethanesulfonate while maintaining the pH at 7.2, then moving to a 30 °C incubator. Thiolysis was monitored by RP-HPLC, and ESI-MS, and was complete within 12 hours. Thiolyzed mutant ubiquitin (aa1-75)-thioester was dialyzed into 0.05% trifluoroacetic acid at 4 °C, then freeze dried, resuspended in a minimal volume and separated from Ava intein by RP-HPLC.^51^

The fusion construct of superfolder GFP and human DNMT1 RFTS domain (aa246-495; sfGFP-RFTS) (from LOBSTR Rosetta *E. coli*) was grown from an overnight starter culture (37 °C) of LB medium (Sigma) with kanamycin (50 mg/L) and chloramphenicol (34 mg/L). Growth for overexpression was initiated by adding 10 mL of starter to inoculate each 1 L of the same medium. Cells were grown to an OD600 of 0.6-0.8, then chilled on ice and induced with 0.3 mM IPTG for 16 hours at 16 °C. Cells were harvested by centrifugation (4 k x g, 30 min, 4 °C), and the cell pellet was resuspended in 5 volumes of chilled (4 °C) lysis buffer (50 mM Tris, pH 7.5 at 25 °C, 300 mM NaCl, 0.5 mM TCEP, 10% glycerol). Resuspended pellets were dounced to uniformity and lysed by three passages through a microfluidizer. Supernatant was cleared by centrifugation (12 k x g, 30 min, 4 °C) and the supernatant was passed over 1 mL Ni NTA agarose twice by gravity. Resin was washed with 20 CV lysis buffer, 10 CV wash buffer (50 mM Tris, pH 7.5 at 25 °C, 300 mM NaCl, 0.5 mM TCEP, 10% glycerol, 25 mM imidazole), and eluted with 10 CV elution buffer (50 mM Tris, pH 7.5 at 25 °C, 300 mM NaCl, 0.5 mM TCEP, 10% glycerol, 200 mM imidazole). Elution fractions were checked by Coomassie-stained SDS-PAGE and purified fractions were pooled, and concentrated (Amicon, 30 kDa MWCO). Concentrated sfGFP-RFTS was further purified by size exclusion chromatography by FPLC (superdex200) run in lysis buffer, after which fractions were again checked by SDS-PAGE, pooled, concentrated as before, aliquoted and flash frozen.

His-SUMO-Sirt2 (aa56-356) was prepared as previously described.^52^ Briefly His-SUMO-Sirt2 (from BL21 (DE3) *E. coli*) was grown at 37 °C from an overnight starter culture of Luria-Bertani (LB) medium (Sigma) with kanamycin (50 mg/L) and chloramphenicol (35 mg/L), of which 10 mL was used to inoculate each 1 L of the same medium used for overexpression. Cells were grown with constant shaking (200 rpm) at 37 °C to an OD600 of 0.6-0.8. Culture temperature was dropped to 16 °C, then induced with 0.5 mM IPTG for 16 hours at 16 °C. Cells were harvested by centrifugation (4 k x g, 15 min, 4°C), and the cell pellet was resuspended in 5 volumes of chilled (4 °C) lysis buffer (20 mM Tris, pH 7.8 at 25 °C, 500 mM NaCl, 20 mM imidazole, and 0.5 mM TCEP). Resuspended cell pellets were lysed by three passages through a French press, and the cell lysate was clarified by centrifugation (12,000 x g, 4°C, 30 min). The lysate supernatant was incubated with pre-equilibrated Ni-NTA resin (2 mL resin bed volume/1 L culture) for 1 hour at 4°C. Resin was drained and washed with 10 column volumes (CV) of wash buffer (20 mM Tris, pH 7.8 at 25 °C, 500 mM NaCl, 50 mM imidazole, and 0.5 mM TCEP), followed by 25 CV of high-salt wash buffer (20 mM Tris, pH 7.8 at 25 °C, 2 M NaCl, 20 mM imidazole, and 0.5 mM TCEP), 25 CV of lysis buffer, and eluted with 10 CV of elution buffer (20 mM Tris, pH 7.8 at 25 °C, 500 mM NaCl, 250 mM imidazole, and 0.5 mM TCEP). The elution was concentrated (Amicon, 10 kDa MWCO, 4000 rpm, 4°C), mixed with UlpI protease (1:100 mass:mass, UlpI:Sirt2) to remove the His-SUMO tag, and dialyzed into dialysis buffer (20 mM Tris, pH 7.5 at 25 °C, 150 mM NaCl, 0.5 mM TCEP) overnight at 4°C. UlpI was depleted by re-incubation with pre-equilibrated Ni-NTA resin for 1 hour at 4°C, after which the the flow-through was collected and concentrated (Amicon, 30 kDa MWCO, 4000 rpm, 4°C). The concentrated flow through was purified by Superdex200 in SEC buffer (50 mM Tris, pH 7.5 at 25 °C, 150 mM NaCl, 0.5 mM TCEP). Fractions were evaluated by SDS-PAGE, pooled, and concentrated using an (Amicon, 10 kDa, 4000 rpm, 4°C). Sample concentration was determined by densitometry, after which protein was aliquoted, flash-frozen in liquid nitrogen, and stored at -80°C until use.

His-TEV-Sirt6 (aa1-355) was prepared as previously described.^32^ Briefly, His-TEV-Sirt6 (from LOBSTR Rosetta *E. coli*) was grown at 37 °C from an overnight starter culture of Luria-Bertani (LB) medium (Sigma) with kanamycin (50 mg/L), of which 10 mL was used to inoculate each 1 L of the same medium used for overexpression. Cells were grown with constant shaking (200 rpm) at 37 °C to an OD600 of 0.6-0.8. Culture temperature was dropped to 25 °C, then induced with 0.5 mM IPTG for 18 hours at 25 °C. Cells were harvested by centrifugation (4 k x g, 15 min, 4°C), and the cell pellet was resuspended in chilled (4 °C) lysis buffer (20 mM Tris, pH 7.5 at 25 °C, 500 mM NaCl, 20 mM imidazole, 30 mL buffer / 1 L culture). Resuspended cell pellets were lysed by three passages through a French pressure cell, and the cell lysate was clarified by centrifugation (12,000 x g, 4°C, 30 min). The lysate supernatant was incubated with pre-equilibrated Ni-NTA resin (2 mL resin bed volume/1 L culture) for 1 hour at 4°C. Resin was drained and washed with 10 CV of wash buffer (20 mM Tris, pH 7.5 at 25 °C, 500 mM NaCl, 50 mM imidazole 0.5 mM TCEP), followed by 25 CV of high-salt wash buffer (20 mM Tris, pH 7.5 at 25 °C, 2 M NaCl, 20 mM imidazole, 0.5 mM TCEP), 25 CV of lysis buffer, and eluted with 10 CV of elution buffer (20 mM Tris, pH 7.5 at 25 °C, 500 mM NaCl, 250 mM imidazole, and 0.5 mM TCEP). The elution was concentrated (Amicon, 10 kDa MWCO, 4,000 rpm, 4 °C), mixed with TEV protease (1:100 mass:mass, TEV:Sirt6) to remove the His-TEV tag, and dialyzed overnight into dialysis buffer (20 mM Tris, pH 7.5 at 25 °C, 150 mM NaCl, 20 mM imidazole, 0.5 mM TCEP). His-TEV and uncleaved His-TEV-Sirt6 were depleted by re-incubation with pre-equilibrated Ni-NTA resin for 1 hour at 4°C, after which the flow-through was collected, diluted with excess Heparin buffer A (50 mM Tris, pH 7.5 at 25 °C, 150 mM NaCl, 5 % glycerol, 0.5 mM TCEP), and concentrated (Amicon, 30 kDa MWCO, 4000 rpm, 4°C). The concentrated sample was purified by heparin column (1 mL, Cytiva, #17040601), and eluted with a linear gradient from 0-50 % heparin buffer B (50 mM Tris, 2000 mM NaCl, 5 % glycerol, 0.5 mM TCEP, pH 7.5) over 20 column volumes (CV) at a flow rate of 0.6 mL/min. Fractions were evaluated by SDS-PAGE, pooled, and concentrated using an (Amicon, 10 kDa, 4000 rpm, 4°C). Concentrated heparin elution further purified Superdex200 (GE, #28-9909-44) with Superdex running buffer (50 mM Tris, pH 7.5, 150 mM NaCl, 0.5 mM TCEP). Fractions were again evaluated by SDS-PAGE, pooled, and concentrated using an (Amicon, 10 kDa, 4000 rpm, 4°C). Sample concentration was determined by densitometry, after which protein was aliquoted, flash-frozen in liquid nitrogen, and stored at -80°C until use.

For the demethylase assays, LSD1-CoREST1 (LC) complex was prepared. His6-tagged LSD1 (aa171-852) and His6-tagged CoREST1 (aa286-482), subcloned in pET15b and pET28a vectors respectively, were co-transformed into BL21-CodonPlus (DE3)-RIPL competent cells (Agilent). Transformed cells were inoculated in LB media containing 100 mg/L ampicillin, 50 mg/L kanamycin, and 35 mg/L chloramphenicol at 37°C. The overnight starter culture was subsequently inoculated into a larger volume of LB media. When the culture reached an A600 of 0.6, 0.5 mM isopropyl β-D-1-thiogalactopyranoside (IPTG) was added to induce protein expression at 18°C for 20 hours. After induction, cells were harvested by centrifugation at 2,702g for 10 minutes at 4°C. The cell pellets were resuspended in lysis buffer containing 20 mM Tris (pH 7.8), 200 mM NaCl, 20 mM imidazole, and 0.5 mM tris(2-carboxyethyl)phosphine (TCEP), then lysed using a french press. The lysate was centrifuged at 20,853g for 30 minutes, and the supernatant was incubated with nickel-nitrilotriacetic acid (NiNTA) resin (MCLAB) for 1 hour at 4°C. The resin was washed with lysis buffer and the bound proteins were eluted with lysis buffer containing 200 mM imidazole. The eluates were concentrated and buffer-exchanged against size exclusion chromatography (SEC) buffer (20 mM Tris, pH 7.8, 200 mM NaCl, and 0.5 mM TCEP) using a 10 kDa MWCO concentrator (Amicon, Millipore Sigma). The buffer-exchanged eluates were further purified by FPLC, using a Superdex 200 Increase 10/300 GL size exclusion column (Cytiva) pre-equilibrated with SEC buffer to obtain stoichiometric complexes. The final LC complex, verified to be >90% pure by SDS-PAGE and Coomassie staining, was collected from the major peak (around 12.5 mL elution volume), pooled, and quantified by A280 absorbance for enzymological studies without further concentration. LC protein solutions were flash-frozen and stored at -80°C.

For the deacylase assays, LSD1-CoREST1-HDAC1 (LHC) was expressed and purified following previously reported protocols.^23,36,43^ Full-length human LSD1, N-terminally 3X FLAG-tagged CoREST1 (aa84-482), and full-length human HDAC1 in pcDNA3.1 vectors were co-transfected into HEK293F cells using polyethylenimine (PEI) (Millipore Sigma, cat 408727-100ML). After 48 hours of transfection, the transfected HEK293F cells were harvested by centrifugation at 2,702g for 10 minutes, resuspended in lysis buffer (50 mM HEPES, pH 7.5, 100 mM KCl, 0.3% Triton X-100, 5% glycerol, and a protease inhibitor cocktail (Thermo Fisher Scientific)), lysed by sonication, and centrifuged at 20,853g for 30 minutes at 4°C. The supernatant was purified using anti-FLAG M2 affinity resin (Millipore Sigma). The FLAG tag on CoREST1 was removed by treatment with TEV protease (1:100 M/M TEV to CoREST1 ratio). Further purification was carried out using gel filtration on a Superose 6 10/300 GL column (Cytiva) pre-equilibrated with buffer containing 50 mM HEPES (pH 7.5), 50 mM KCl, and 0.5 mM TCEP. The pooled fractions from the main peak around 13 mL elution volume were concentrated down to approximately 5 μM using a concentrator with a 30 kDa MWCO (Pall). The LHC complex showed a 1:1:1 stoichiometry and was >90% pure, as confirmed by SDS-PAGE with Coomassie staining. The LHC protein samples were aliquoted, flash-frozen, and stored at -80°C.

The expression and purification of MiDAC (DNTTIP1, MIDEAS and HDAC1) was performed as described previously.^35^ Plasmids (pcDNA3) encoding DNTTIP1, HDAC1, and a N-terminally 10xHis-3xFLAG-TEV tagged MIDEAS were co-transfected into HEK293F cells with PEI (Sigma). After 48 hours of growth, cells were resuspended in resuspension buffer (50 mM Tris, pH 7.5, 100 mM potassium acetate, 10% (v/v) glycerol, 0.5% (v/v) Triton X-100, Complete EDTA-free protease inhibitor (Roche)) and lysed by sonication. Insoluble material was removed by centrifugation, and the supernatant was passed over anti-FLAG immunoaffinity resin. The bound complex was washed twice with resuspension buffer, three times with wash buffer (50 mM Tris/Cl pH 7.5, 50 mM potassium acetate, 5% (v/v) glycerol, 0.5 mM TCEP), and then incubated with RNaseA for 1 h at 4 °C. After RNAse incubation the resin was washed a further five times with wash buffer. MiDAC complex was liberated from the resin by overnight incubation with TEV protease on a roller at 4 °C. The mixture was purified further using gel filtration on a Superdex 200 Increase 10/ 300 GL column (run in 25 mM HEPES, pH 7.5, 50 mM potassium acetate, 0.5 mM TCEP). The purified complex was concentrated to 1–5 mM and analyzed by SDS-PAGE stained with Coomassie to confirm purity (>80%). Glycerol was added to 10% (final concentration), then samples were aliquoted, flash frozen, and stored at -80 °C until use.

HDAC1 was purchased from BPS Bioscience (cat 50051). Sirt1 was purchased from BPS Bioscience (cat 50081). Sirt5 was purchased from BPS Bioscience (cat 50085).

### Mammalian cell growth, nuclear acid extraction and histone tail isolation

HEK293T cells (ATCC) were grown in DMEM supplemented with 5% FBS and 1% penn/strep (37 °C, 5% CO_2_), and regularly tested by PCR for mycoplasma contamination. All experiments were conducted within 10 passages of stock thawing. For treatment with DMSO vehicle, MS275 (LC Laboratories, Cat# E-3866, Lot: ENT-102) or Corin (Med Chem Express, Cat# HY-111048/CS0034060, Lot: 41808), a single 15 cm plate of HEK293T cells was grown to near confluence and split. Cells were washed with Dulbecco’s phosphate buffered saline (DPBS, D8537, Sigma-Aldrich), liberated with TryplE, and quenched with DMEM, then cell count and viability (99%) were determined by Trypan blue staining (Countess III). Plates (10 cm) were seeded in triplicate (4.4e6 cells / plate) and recovered for two days in DMEM to ∼50% confluence. Following media change, stocks of drug were prepared in sterile filtered DMSO, then diluted in DMEM and added to plates to a final concentration of 10 µM MS275, or 2 µM Corin, and 1% DMSO. Cells were incubated for 6 hours in the presence of vehicle or drug, after which the media was aspirated, cells were washed with DPBS, liberated with TryplE, and quenched with DMEM. Cells were pelleted and the supernatant removed by vacuum, then washed once with cold (4 °C) DPBS, pelleted again, and the supernatant was again removed by vacuum before flash freezing the pellets.

Pellets were thawed on ice and gently resuspended in 10 pellet volumes of nuclear isolation buffer (15 mM Tris, pH 7.5 at 25 °C, 60 mM KCl, 15 mM NaCl, 5 mM MgCl_2_, 1 mM CaCl_2_, 250 mM sucrose, 5 mM sodium butyrate, 0.3% NP-40, 1 x Pierce protease inhibitor). After incubating for 5 minutes on ice nuclei were pelleted (600 x g, 5 min, 4 °C) and the supernatant was removed. Nuclear pellets were washed twice by gentle pipetting with the same volume of detergent free (no NP-40) nuclear isolation buffer, pelleting and removing the supernatant as before. To the pellet was added 5 initial pellet volumes of ice cold 0.4 N H2SO4, followed by vigorous mixing, then 2 hours of constant agitation at 4 °C. Insoluble material was separated by centrifugation (3400 x g, 10 min, 4 °C), and the supernatant was collected. The supernatant was dialyzed against chilled 2.5% acetic acid, then twice against chilled 0.05% trifluoroacetic acid. The concentration of histone protein in the dialyzed supernatant was estimated by A280 (nanodrop), and the solution was then aliquoted, flash frozen and freeze dried.

Acid extracted protein was resuspended in reagent grade water to 100 μM in histone H3. This was diluted into reaction buffer (final concentration 40 mM PIPES, pH 7.5, 1 mM DTT, 5 mM CaCl_2_) to a final H3 concentration of 20 μM. To this was added cW11 sortase to 100 μM and a single oligoglycine TMT peptide to 1 mM. The reactions were placed in a 37 °C incubator overnight (12-16 h), then moved onto ice and chilled for 10 minutes. Saturated trichloroacetic acid was slowly added to 5% of the reaction volume, and placed back on ice for 5 minute, then vigorously mixed by vortexing, and incubated for a further 10 minutes on ice. Insoluble material was pelleted by centrifugation (16 k x g, 10 min, 4 °C) and the supernatant was recovered. The supernatant was buffer exchanged against ice-cold 0.05% trifluoroacetic acid in a pre-equilibrated spin concentrator (Amicon, 3 kDa MWCO, 14 k x g, 10 min, 4 °C) for a total of 5 rounds. The buffer exchanged solution was recovered and freeze dried. The sample was resuspended in reagent grade water and one third was taken for duplicate quantification against an amino acid analyzed standard by 16% Tris-Tricine gel. Samples were again freeze dried, then resuspended in 0.05% trifluoroacetic acid, and cleaned up using stage tips packed in house, and concentrated by speed vac.

### Middle-down proteomics

Histone tails were resuspended in 0.1% formic acid and mixed to a 1:1:1:1:1 mass ratio based on gel quantification. Mixtures of DMSO treated samples (TMT-126, 127, 128) and samples treated with either MS275 or Corin (TMT-129, 130, 131) were prepared to a concentration of 10 ng/uL and mixed thoroughly. 20 ng was injected for each analysis. Samples were analyzed on a Vanquish Neo LC (Thermo Fisher Scientific) coupled to an Orbitrap Ascend Tribrid mass spectrometer equipped with ETD, PTCR, and UVPD (Thermo Fisher Scientific). The LC was operated in trap-and-elute mode using a PepMap™ Neo C18, 100 Å, 5 μm, 0.3 x 5 mm Trap Cartridge (Thermo Fisher Scientific) and an Easy-Spray PepMap Neo C18, 100 Å, 2 μm, 75 μm x 150 mm column (Thermo Fisher Scientific) heated to 40°C. The flow rate was 0.3 μL/min. Solvent A was 0.1% formic acid and Solvent B was 0.1% formic acid in acetonitrile. The LC gradient consisted of 2 to 15% Solvent B in 40 min, followed by 15 to 25% Solvent B in 6 min. The mass spectrometer was operated in data-dependent acquisition mode with a 2 second cycle time. MS1 spectra were collected in the Orbitrap with a resolution of 60k and scan range of 375-600 m/z. Charge state and precursor selection range filters were applied to select peptides with a charge state of 8 from 480-540 m/z for MS2. Dynamic exclusion was enabled, and precursors were excluded for 9 seconds after being selected 1 time. MS2 spectra were collected with an isolation window of 1.6 m/z, a resolution of 30k, and 2 microscans. EThcD was performed with an ETD reaction time of 20 ms, ETD reagent target of 1E5, max ETD reagent injection time of 20 ms, supplemental activation collision energy of 15%, and normalized AGC target of 200%.

Raw data (available at https://doi.org/10.7910/DVN/3IOGQ8) was analyzed using Byonic MS/MS search engine (Protein Metrics) using the following settings: No protease cleavage; 500 ppm precursor tolerance; 15 ppm fragment tolerance; maximum 5 precursors; fragmentation type EThcD; maximum 6 common PTMs; Common PTMs Methyl / +14.015650 @ K, R, max 3 | Dimethyl / +28.031300 @ K, R, max 3 | Trimethyl / +42.046950 @ K, max 3 | Phospho / +79.966331 @ S, T, max 2 | Acetyl / +42.010565 @ K, max 5; maximum 1 rare PTM; Propionyl / +56.026215 @ K | rare1; fixed modification of H with TMT 6-plex; TMT6plex / +229.162932 @ H; fixed modification of protein C-terminus with 76.1001 Da mass (accounting for the mass difference between H and [K + 2,3-diaminopropionamide]). Data was searched against a fasta database of histone protein sequences, common contaminants, and the following sequences for H3.1 and H3.3 peptides: H3.1 ARTKQTARKSTGGKAPRKQLATKAARKSAPATGGGH H3.3 ARTKQTARKSTGGKAPRKQLATKAARKSAPSTGGGH

Peptide spectrum matches (PSMs) from Byonic were exported and subsequently processed in R (4.3.2) using RStudio, custom scripts (available at https://doi.org/10.7910/DVN/3IOGQ8) and packages parallel (4.3.2), stats (4.3.2), purrr (1.0.2), mzR (2.36.0), tidyr (1.3.1), stringr (1.5.1), dplyr (1.1.4), xml2 (1.3.6), readr (2.1.5), readxl (1.4.3), writexl (1.5.0), tidyverse (2.0.0), RColorBrewer (1.1-3) and ggrepel (0.9.5). Briefly, PSMs were filtered for high confidence matches with a posterior error probability ≤ 0.05, and a Byonic delta mod score ≥ 10 (a measure of confidence in PTM sequence location). Spectra were further subdivided by the number of PSMs reported in Byonic, and then EThcD ions (a, b, c, y, z, and neutral loss; 15 ppm error tolerance) consistent with the PSM(s) were identified within the spectrum. Fragment ions with at least one heavy isotope peak (e.g. ^13^C) were treated as high confidence and used to determine proteoform abundance in the sample as a whole. This criterion was not applied to TMT reporter ions, the isotopic impurities in which were processed separately. Proteoform abundance was calculated as the fraction of annotated fragment ion intensity attributable to a single proteoform relative to the sum of annotated fragment ion intensity for all proteoforms. For spectra assigned multiple PSMs, intensity from shared fragment ions was divided between the relevant PSMs. Reporter ion signal intensities were channel normalized to the maximum average channel (e.g. TMT-126) signal intensity, averaging over all spectra with a PSM. Mean and Median channel normalization produced broadly similar results (**Table S2-S5**). Channel normalized reporter ion signal intensities were log2 transformed for statistical analysis. Spectra/scans with ≥2 reporter ions per treatment condition were used in quantification, and all such scans attributable to a single proteoform (e.g. H3K14ac/K23ac/K27me2) were grouped. For each treatment condition (drug or vehicle) all log2 transformed reporter ion signals from a proteoform grouping were averaged, and the difference between treatment condition averages reported as the log2-fold change in proteoform abundance. Statistical significance was assessed by two-sided T test (p<0.05) using R stats (4.3.2).

### Bottom-up proteomics

Nuclear acid extracts were resuspended in propionylation buffer (50 mM NH4 HCO3, pH 8.0), to 1 μg/μL, and mixed with 0.5 volumes of 25% propionic anhydride in acetonitrile. The reaction was pH adjusted to 8.0 with ammonium hydroxide, and incubated for 15 minutes at 25 °C. The reaction was mixed with a further 0.5 volumes of fresh 25% propionic anhydride in acetonitrile, and again adjusted to pH 8.0 with ammonium hydroxide. After incubation for 15 minutes at 25 °C the reactions were concentrated by speedvac. The dried, propionylated samples were suspended in digest buffer (50 mM NH_4_ HCO_3_, pH 8.0) to 1 μg/μL and adjusted to pH 8.0 with ammonium hydroxide. Sequencing grade trypsin was added to 2% (mass/mass) of the total histone protein, and incubated overnight at 25 °C. After trypsination samples were concentrated by speedvac, and propionylation was repeated as described above. Dried samples were resuspended in 0.05% trifluoracetic acid, cleaned up using stage tips packed in house, and concentrated by speed vac.

Samples were resuspended in 0.1% acetic acid and (20 ng / injection) analyzed on a Vanquish Neo LC (Thermo Fisher Scientific) coupled to an Orbitrap Ascend Tribrid mass spectrometer equipped with ETD, PTCR, and UVPD (Thermo Fisher Scientific). The LC was operated in trap-and-elute mode using a PepMap™ Neo C18, 100 Å, 5 μm, 0.3 x 5 mm Trap Cartridge (Thermo Fisher Scientific) and an Easy-Spray PepMap Neo C18, 100 Å, 2 μm, 75 μm x 150 mm column (Thermo Fisher Scientific) heated to 40°C. The flow rate was 0.3 μL/min. Solvent A was 0.1% formic acid and Solvent B was 0.1% formic acid in acetonitrile. The gradient consisted of 2 to 32% Solvent B in 46 min and 32 to 42% B in 7 min. The MS was operated in data independent acquisition (DIA) mode. MS1 were collected in the Orbitrap with 120k resolution and scan range 290-1200 m/z. HCD MS2 were performed with collision energy 25% and resolution 15k. The DIA method consisted of 34 fixed-width isolation windows of 24 m/z with 1 m/z overlap and center masses from 307 to 1093.25.

Raw data was analyzed in Skyline and further processed with in-house software EpiProfile.^53^ Each biological replicate of a treatment condition was analyzed separately. For comparison with middle-down results, site-specific PTM abundances from each replicate were averaged across all samples grouped for a single 6plex middle-down analysis. Pearson correlation was used to compare site-specific PTM abundance measurements between bottom-up and middle-down data.

### Peptide synthesis

Histone H3.1 peptides (aa1-34) and oligoglycine tandem mass tag peptides were prepared by Fmoc-based solid phase peptide synthesis with modified amino acids at the relevant positions. Peptides were synthesized using a Prelude Peptide Synthesizer (Gyros Protein Technologies) using a standard deprotection and coupling protocol (vide infra). Modified amino acids were installed by either the standard protocol, or one of the following manual coupling protocols:

Standard deprotection: 20% piperidine in DMF, 2 rounds of 10 minutes, then 7 DMF washes.

Standard coupling: 4 eq. AA, 3.75 eq. HATU, 0.2 M NMM in DMF, 2 rounds of 90 minutes, then 5 DMF washes.

Tandem Mass Tag: Nα-Fmoc-Nβ-ivDde-2,3-diaminopropionic acid; standard coupling, followed by capping with 10 eq. acetic anhydride and 10 eq. DIEA.

Acetylation: Fmoc-K(Ac)-OH; standard coupling.

Dimethylation: Fmoc-K(Me2)-OH; 2 eq., 1.75 eq HATU, 4 eq DIEA, 1 round, 90 minutes, followed by capping with 10 eq. acetic anhydride and 10 eq. DIEA.

Trimethylation – Fmoc-K(Me3)-OH HCl; 2 eq., 1.75 eq HATU, 4 eq DIEA, 2 rounds, 90 minutes, followed by capping with 10 eq. acetic anhydride and 10 eq. DIEA. Acylations – Fmoc-K(alloc)-OH; standard coupling.

Ubiquitination – Fmoc-K(Boc-Cys(Trt))-OH; 2 eq., 1.75 eq HATU, 4 eq DIEA, 2 rounds, 90 minutes, followed by capping with 10 eq. acetic anhydride and 10 eq. DIEA.

In each case Ala1 of the H3 sequence was coupled as Boc-Ala-OH. Deprotection of K(alloc) was accomplished with Pd(PPh_3_)_4_ (0.35 eq) and phenylsilane (20 eq) in anhydrous dichloromethane for 2 rounds of 60 minutes. Subsequent acylation was achieved with 4 eq. carboxylic acid, 3.75 eq. HATU, and 16 eq. DIEA in DMF for 2 rounds of 60 minutes.

In synthesis of the tandem mass tag (TMT) peptide each standard coupling was followed by capping with 4 eq. 3-phenylpropionic acid, 3.75 eq. HATU, 0.2 M NMM in DMF, for one round of 20 minutes. The most N-terminal glycine in the TMT peptide was coupled as Boc-Gly-OH. Deprotection of the 2,3-diaminopropionic acid Nβ-ivDde was accomplished with 5% hydrazine in DMF for ten rounds of 5 minutes, followed by 5 rounds of washing with DMF, and one round of washing with dichloromethane. The resin was subsequently divided into six fractions of approximately equal mass, each of which was treated with 1 eq. of TMT NHS ester reagent resuspended in anhydrous DMF, to which was added 4 eq. of silica plug dried DIEA.

Peptides were cleaved (90% trifluoroacetic acid, 5% water, 5% triisopropylsilane) and ether precipitated. Histone tail peptides were resuspended in 0.05% trilfuoroacetic acid and purified by preparative RP-HPLC (C18; mobile phase A: water + 0.05% TFA; mobile phase B: acetonitrile +0.05% TFA; 7-40% B 40 minute gradient). TMT peptides were resuspended in mobile phase A and purified by analytical RP-HPLC (Eclipse XDB-C18 4.6 x 250 mm, 0% B isocratic 13 minutes, 0-95%B 15 minutes). Fraction purity was assessed by ESI-MS (Thermo Q Exactive), and the purest fractions were pooled. H3 tail peptides were quantified by analytical RP-HPLC against a standard curve prepared from unmodified H3(1-34) peptide.

Ubiquitinated peptides were prepared by denaturing expressed protein ligation of ubiquitin (aa1-75)-mesna thioester (2.5 mM) with H3 (aa1-34) peptide (0.5 mM) in ligation buffer (250 mM HEPES, pH 7.5, 6 M Guanidine HCl, 30 mM TCEP, 30 mM methylthioglycolate) under argon. Reactions were mixed overnight at 37 °C and checked by analytical RP-HPLC followed by ESI-MS and SDS-PAGE. Once judged complete, one volume of desulfurization buffer (250 mM HEPES, pH 7, 6 M guanidine HCl, 500 mM TCEP, 60 mM VA-044) was added, and the reaction was placed under argon before mixing overnight at 37 °C.^54^ Unligated peptide was removed by RP-HPLC (Higgins Analytical, C18; 30-37% B gradient), then fractions containing the ligation product were identified by ESI-MS and SDS-PAGE and pooled. Ligated peptide did not separate from hydrolyzed ubiquitin (aa1-75), and was quantified by SDS-PAGE relative to a BSA standard curve.

### Nucleosome reconstitution and sortase ligation

Into chilled nucleosome reconstitution buffer (final concentrations 2 M KCl, 10 mM Tris, pH 7.9 at 25 °C, 1 mM EDTA, 10 mM DTT) Widom 601 DNA was added to 6 µM, followed by octamer (6 – 10 µM, typically 6.6 µM). Small scale nucleosome reconstitutions (50-100 µL) are carried out with new batches of octamer and DNA to determine the optimal octamer concentration for reconstitution. For both small scale and preparative reconstitutions the mixture of DNA and octamer are incubated for 30 minutes on ice, then transferred to pre-chilled dialysis cassettes. These were transferred to pre-chilled high salt reconstitution buffer (10 mM Tris, pH 7.9 at 25 °C, 2 M KCl, 1 mM EDTA, 1 mM DTT; 500 mL for small scale; 1 L for large scale) and dialyzed over a linear salt gradient created by constant influx (1 mL / min for small scale; 2 mL / min for large scale) of low salt reconstitution buffer (10 mM Tris, pH 7.9 at 25 °C, 150 mM KCl, 0.1 mM EDTA, 1 mM DTT; 2 L for small scale; 5 L for large scale) and efflux of buffer from the dialysis chamber, mediated by a dual channel peristaltic pump.^55^ This exchange occurs with constant vigorous mixing over 30-48 hours, after which dialysis cassettes are transferred to low salt reaction buffer for a final 2 hours of dialysis.

The crude nucleosome reconstitution within the dialysis cassette is directly used in the sortase ligation reaction. Reconstituted nucleosome (1.65 µM), followed by peptide (symmetric ligation: 49.5 µM; first asymmetric ligation: 16.5 µM; second asymmetric ligation: 49.5 µM) and then cW11 sortase (200 µM) are diluted into reaction buffer (40 mM Tris, pH 7.5 at 25 °C, 150 mM NaCl/KCl, 5 mM CaCl_2_, 1 mM DTT final concentration; KCl from the reconstituted nucleosome and NaCl in the buffer are treated as equivalent for the purpose of determining the final concentration of monovalent cation chloride salt). For ligations employing a charged C-terminal auxiliary (e.g. tetra- and penta-acetylated histone tails) the nucleosome and peptide are sequentially added to reaction buffer and incubated for 10 minutes at 25 °C before the addition of cW11 sortase. Reactions are transferred to a 37 °C incubator for at least 4 hours. Overnight reactions are generally well-tolerated, but should be avoided for hyperacetylated substrates. Reactions are quenched by addition of one eq. salmon sperm DNA (mass/mass, relative to nucleosome DNA; Fisher) and NaCl to 250 mM, mixing, then sitting for 5 minutes at 25 °C before purification.

Purification was accomplished by weak anion exchange (Tosoh DEAE 5-pW, 7.5 mm x 7.5 cm) liquid chromatography (Waters 1525 Binary HPLC Pump, 1 mL / min, mobile phase A: 10 mM Tris, pH 7.9 at 25 °C, 150 mM KCl, 0.5 mM EDTA; mobile phase B: 10 mM Tris, pH 7.9 at 25 °C, 600 mM KCl, 0.5 mM EDTA). Purification gradients by ligation type (symmetric v. asymmetric) and DNA length (147 bp v. 185 bp) are as follows:

147 bp, symmetric: 0-22% B (3 min), 22% B (7 min), 22-49% B (1 min), 49-65% B (21 min), 65-100% B (1 min), 100% (10 min), 100-0% B (1 min)

147 bp, asymmetric: 0-22% B (3 min), 22% B (7 min), 22-49% B (1 min), 49-61% B (21 min), 65-100% B (1 min), 100% (10 min), 100-0% B (1 min)

185 bp, symmetric: 0-22% B (3 min), 22% B (7 min), 22-63% B (1 min), 63-80% B (21 min), 65-100% B (1 min), 100% (10 min), 100-0% B (1 min)

185 bp, asymmetric: 0-22% B (3 min), 22% B (7 min), 22-58% B (1 min), 63-75% B (21 min), 65-100% B (1 min), 100% (10 min), 100-0% B (1 min)

Fractions collected during purification are directly diluted with one volume of chilled dilution buffer (10 mM Tris, pH 7.5 at 25 °C, 1 mM DTT) and spin concentrated (Amicon, 10 kDa MWCO). Fractions of purity may be assessed by SDS-PAGE followed by western blotting against histone H3 (25 ng H3 loading, Abcam, ab1791), to confirm conversion of all H3 (aa33-135) to H3 (aa1-135), or blotting against relevant PTMs for asymmetric syntheses. Pooled, concentrated fractions are dialyzed against either dilution buffer (symmetric ligation), or low salt reconstitution buffer (asymmetric ligation). Following dialysis against low salt reconstitution buffer, a second round of ligation and purification are performed as described above (note that the equivalents of peptide increase in the second round of ligation). Following dialysis into dilution buffer, symmetric and asymmetric nucleosomes are dialyzed into storage buffer (10 mM Tris, pH 7.5 at 25 °C, 25 mM NaCl, 1 mM DTT, 20% glycerol) and concentrated to >5 µM before flash freezing and storing at -80 °C.

### Cryo-EM

In-situ fixation of cW11 nucleosome for structural studies was performed according to Worden.^56^ 300 mM nucleosome solution was dialyzed against crosslink buffer (25 mM HEPES pH 7.5, 25 mM NaCl, 1 mM EDTA,1 mM DTT) and mixed with 2.44 ml of 0.14% glutaraldehyde. The crosslinking reaction was incubated on ice for 60 min. The reaction was quenched for 1 hour on ice by adding 1.0 M Tris pH 7.5 to the final concentration of 100 mM. Finally, the sample was dialyzed against crosslink buffer and concentrated using an Amicon Ultra 30K MWCO spin concentrator to a final 0.2 mg/ml concertation, according to DNA absorption at 260 nm.

Cryo-EM grids of the cW11 nucleosome were prepared following an established protocol: 3.0 μL of the samples at 0.2 mg/mL were applied to glow-discharged Quantifoil gold grids (400 mesh, 1.2 μm hole size).^57^ The grids were then blotted for 3 seconds at 4ºC and 100% humidity and plunge-frozen using Vitrobot Mark IV (FEI Company). All sample images were recorded on FEI Talos Arctica operated at 200kV using Gatan K2 Summit direct electron detector camera in counting mode at a nominal magnification of 130,000x (calibrated pixel size of 1.096 Å/pixel). The total accumulated electron exposure was 50.77 electrons per Å2.

Raw Cryo-EM Images were motion corrected using UCSF MotionCor2 v1.2.1; Patch CTF was used to calculate CTF, Blob Picker for particle picking, Extract Micrograph for particle extraction, 2D classification, Ab Initio for Ab Initio reconstruction and hetero, homo and Non-homogeneous refinement for 3D refinement; all implemented on CryoSPARC 4.5.1.^58,59^ This procedure led to obtaining reconstructions of cW11 nucleosome at 4.7 Å (FSC = 0.143).

For the model fitting of the canonical nucleosome into the cryo-EM map, we fitted the X-ray structure of the X. Laevis Nucleosome Core Particle (PDBID:1KX5)^60^ using the map fit feature included in ChimeraX 1.4.^61^ All structure images were generated using ChimeraX 1.4.

### Deacylase kinetics

All deacetylation assays and analyses were performed as previously reported, these include deacetylation of symmetric H3 penta- and mono-acetylated nucleosomes (MiDAC, Sirt2, and Sirt6), deacetylation of asymmetric H3 mono-acetylated/unmodified or H3mono-acetylated/tetra-acetylated nucleosomes (MiDAC, Sirt2, and Sirt6), deacylation of symmetric H3 acylated nucleosomes, and comparative deacetylation of nucleosomes prepared by established methods and the sortase ligation method (LHC and Sirt6). Symmetric nucleosomes (H3K9ac, H3K14ac, H3K18ac, H3K23ac, H3K27ac, H3pentaAc, H3K9pr, H3K9bu, H3K9cro, H3K9hib, H3K9succ) and asymmetric nucleosomes (H3K18ac, H3K23ac, H3K27ac) (final 100 nM) were diluted into HDAC reaction buffer (50 mM HEPES at pH 7.5, 100 mM KCl, 0.2 mg/mL BSA, and 100 μM inositol hexaphosphate (IP6)), or Sirtuin reaction buffer (50 mM HEPES at pH 7.5, 1 mM DTT, 0.2 mg/mL BSA, and 1 mM NAD). The reaction solution was kept on ice until the addition of HDAC complex (LHC, MiDAC, HDAC1 free enzyme) or Sirtuin (Sirt1, Sirt2, Sirt5, Sirt6) then incubated at 37 ºC. Each complex was assayed at a minimum of two enzyme concentrations to ensure consistency in calculated V/[E] values. At each assay time points, replicate aliquots were taken from a single reaction, and quenched. Typically, 6.5 μL aliquots of the reaction were taken and quenched with Dual quenching buffer (6.5 μL of a 1:1 mixture of 4 × Laemmli sample buffer and 40 mM EDTA) such that the final quenched aliquot contains 1 × Laemmli sample buffer with 10 mM EDTA. For typical nucleosome assays, time points were collected at 0, 30, 60, 90 and 120 min. Each sample was then boiled for 3-5 min at 95 ºC and resolved on a 4-20 % gradient SDS-PAGE gel (TGX^TM^, Bio-Rad, 4561096) at 180 Volts for ∼30 min. Gels were then transferred to nitrocellulose membrane (Transfer Stack, Invitrogen, IB301031) for western blot analysis (WB) by iBlot (Invitrogen) with P3 (20 V) for 5.0 min. Membranes were blocked for 60 minutes with 5% BSA in TBST (20 mM Tris pH 7.5 at 25 C, 150 mM NaCl, 0.1% Tween-20) then incubated overnight at 4 ºC with primary antibody in 5% BSA in TBST. Site-specific antibodies for acetylated H3 and acylated H3 (1:2000 unless noted otherwise), such as for anti-H3K9ac (Abcam AB32129), anti-H3K14ac (EMD Millipore 07-353), anti-H3K18ac (EMD Millipore 07-354), anti-H3K23ac (EMD Millipore 07-355), anti-H3K27ac (Cell Signaling 81735),, anti-H3K9bu (PTM Biolabs, PTM-305), anti-H3K9cro (PTM Biolabs, PTM-539), pan anti-Khib (A kind gift from Dr. Yingming Zhao, University of Chicago, 1:1000 dilution), and anti-H3K9succ (A kind gift from Dr. Yingming Zhao, University of Chicago, 1:1000 dilution) were used to blot the corresponding membranes. Meanwhile, anti-H3 (Abcam, #ab1791, 1:2000 dilution) was used to visualize total H3. The specificities of the primary antibodies against H3K9ac, H3K14ac, H3K18ac, H3K23ac, and H3K27ac were validated on both H3 mono-acetylated and H3 penta-acetylated nucleosome substrates.

After washing, membanes were incubated with an HRP-linked secondary antibody (either anti-Rabbit IgG, Cell signaling #7074S; or anti-Mouse IgG, Cell signaling #7076S) for 1 h at room temperature, then washed. Membranes were then treated with ECL substrate reagent (Bio-Rad, #170-5061), and visualized by G:BOX mini gel imager (Syngene). The bands on the membrane were then quantified by ImageJ (Download from imagej.nih.gov/ij/). All intensity values were divided by the intensity value at t=0 to get relative intensity, and then fit to a single-phase exponential decay curve with constrain Y0=1, Plateau=0 (GraphPad Prism 10). Each plotted point represents at least 2 replicate measurements from the same assay. The kinetic parameter V/[E] was calculated, log10 transformed and plotted as heatmaps using GraphPad Prism 10.

### Demethylase kinetics

All demethylase assays and analyses were performed as previously reported.^34^ Three different 185 bp asymmetric nucleosomes with the following modifications (tail modification #1/tail modification #2) were prepared: 1) H3K4me2/H3_unmodified_, 2) H3K4me2/H3K14ac, and 3) H3K4me2-K14ac/H3_unmodified_.The nucleosomes at 100 nM were mixed with LSD1-CoREST1 (LC) at 180 nM in a reaction buffer containing 50 mM HEPES at pH 7.5, 7 mM KCl, 2.1% glycerol, and 0.2 mg/mL BSA at 25°C. At 0 min, 10 min, 30 min, and 60 min timepoints, 18 µL samples were mixed with 12 µL 4X SDS sample loading buffer, followed by heating at 95°C for 1 min. Then, the samples were resolved by SDS PAGE using 4-20% gradient precast Tris-glycine gels (Thermo Fisher Scientific), transferred to nitrocellulose membranes (iBlot, Thermo Fisher Scientific), blocked with a 1X TBST buffer supplemented with 5% BSA, washed with 1X TBST, and blotted with anti-H3K4me2 (1:2000 dilution, Abcam, cat ab32356) and anti-H3 antibodies (1:2000 dilution, Abcam, cat ab1791). After washing the membranes with 1X TBST, secondary antibodies were added and incubated for 1 hour at room temperature (anti-rabbit IgG HRP-linked antibody, 1:1000 dilution, Cell Signaling Technology, cat 7074S). After the incubation, the membranes were washed with 1X TBST. Each membrane was visualized by ECL (Clarity, Bio-Rad) using a gel imager (G:Box mini, SynGene) and the Genesys (v1.8.5.0) software. The density of each anti-H3K4me2 and anti-H3 band was quantified by ImageJ (NIH), the data were normalized by total H3 at each time point. The relative H3K4me2 intensities (normalized by total H3, and relative to T0) at each time point were fitted into an exponential decay function using GraphPad Prism 9 with the constraints of Y0 at 1 and plateau at 0. The extrapolated rates were converted to V/[E], min^-1^. All measurements were done in 4 replicates, and One-way ANOVA analysis with Dunnett’s multiple comparisons test, with a single pooled variance was employed to compare statistical differences in LC complexes.

### Electrophoretic mobility shift assays

Modified 185 bp nucleosomes were thawed on ice and serially diluted in nucleosome storage buffer (10 mM Tris, pH 7.5 at 25 °C, 25 mM NaCl, 1 mM DTT, 20% glycerol) on ice. A solution of sfGFP-RFTS protein in RFTS storage buffer (50 mM Tris, pH 7.5 at 25 °C, 300 mM NaCl, 0.5 mM TCEP, 10% glycerol) was thawed on ice. These were combined in EMSA buffer (15 mM Tris, pH 7.5 at 25 °C, 105 mM NaCl, 0.7 mM DTT, 0.3 mg/mL BSA, 4% glycerol final concentrations) to final concentrations of 20 nM sfGFP-RFTS and 0-160 nM nucleosome, and incubated at 4 °C for 1 h or 24 hours. After incubation, samples were mixed with native sample loading buffer (2.5 x TBE, 50% glycerol), loaded on native 4-20% TBE gels. Gels were run in 0.5 x TBE for 2 hours at 100 V, then sfGFP fluorescence was visualized (Typhoon 5, ex: 488 nm, em: 532 nm). After fluorescent visualization gels were stained with ethidium bromide (30 minutes), rinsed, and DNA was visualized. This titration and visualization was repeated with the concentration of nucleosome fixed at 20 nM, and the concentration of sfGFP-RFTS varied from 0-160 nM. The fraction bound was quantified in ImageJ, and data was fit using GraphPad Prism 10.2.2 (Nonlinear, one site specific binding).

## Acknowledgments

The authors gratefully acknowledge funding from the following sources: NIH GM149229 (to P.A.C.), Leukemia and Lymphoma Society SCOR (to P.A.C.), NSF 2127882 (to P.A.C. and B.A.G.), NIH P01CA196539 (to B.A.G.), R01HD106051 (to B.A.G.), Wellcome Trust Investigator Award 222493/Z/21/Z (to J.W.R.S.), University of Chicago, Nancy and Leonard Florsheim family fund (to Y.Z), NIH AR078555 (to Y.Z), NIH CA251677 (to Y.Z), NIH 2R01GM115882-06 (to K-J.A.), NIH 1R01CA266978-01 (to K-J.A.), NIH 1R01GM144547-01A1 (to K-J.A.), Charles King Trust Postdoctoral Fellowship (to S.D.W.), and Thermo Scientific Tandem Mass Tag Systems Research Award (to S.D.W.). Histone/Nucleosome depiction in figures 1, 2, 3, and 5 are derived from chromatin-structure icon by Pauline Franz https://twitter.com/_paulinefranz is licensed under CC-BY 4.0 Unported https://creativecommons.org/licenses/by/4.0/.

## Author Contributions

The project was conceptualized by P.A.C., B.A.G., M.W., C.L., S.D.W., K.L., and Z.A.W. Data was collected by M.W., C.L., S.D.W, K.L., Z.A.W., E.Z., M.Y-A., E.N., S.D-C., P.D.I., A.A., E.L., and J.J. Mass spectrometry data was analyzed by S.D.W. and E.Z. with input from C.L. and X.S. Enzymology data was analyzed by K.L., Z.A.W., M.Y-A., and J.J. Cryo-EM data was analyzed by P.D.I. Materials were provided by S.D.W., K.L., Z.A.W., M.Y-A., E.N., S.D-C., L.F., P.D.I., X.S., A.A., E.L., K-J.A., Y.Z., J.W.R.S., M.W., B.A.G. and P.A.C. Funding was acquired by K-J.A., Y.Z., J.W.R.S., B.A.G. and P.A.C. The project was supervised by K-J.A., Y.Z., J.W.R.S., M.W., B.A.G. and P.A.C. The draft manuscript was composed by P.A.C. and S.D.W. All authors reviewed the results and approved the final version of the manuscript.

## References

1. Jenuwein, T. & Allis, C. D. Translating the histone code. Science 293, 1074–1080 (2001).

2. Becker, J. S. et al. Genomic and Proteomic Resolution of Heterochromatin and Its Restriction of Alternate Fate Genes. Mol Cell 68, 1023–1037.e15 (2017).

3. Siegenfeld, A. P. et al. Polycomb-lamina antagonism partitions heterochromatin at the nuclear periphery. Nat Commun 13, 1–16 (2022).

4. Zhao, Y. & Garcia, B. A. Comprehensive catalog of currently documented histone modifications. Cold Spring Harb Perspect Biol 7, (2015).

5. Huang, H., Lin, S., Garcia, B. A. & Zhao, Y. Quantitative Proteomic Analysis of Histone Modifications. Chem Rev 115, 2376–2418 (2015).

6. Klein, B. J. et al. Histone H3K23-specific acetylation by MORF is coupled to H3K14 acylation. Nat Commun 10, (2019).

7. Marunde, M. R. et al. Nucleosome conformation dictates the histone code. eLife 13, (2024).

8. Luger, K., Rechsteiner, T. J. & Richmond, T. J. Preparation of nucleosome core particle from recombinant histones. Methods Enzymol 304, 3–19 (1999).

9. Weller, C. E. et al. Aromatic thiol-mediated cleavage of N-O bonds enables chemical ubiquitylation of folded proteins. Nat Commun 7, 1–10 (2016).

10. Li, S. & Shogren-Knaak, M. A. Cross-talk between histone H3 tails produces cooperative nucleosome acetylation. Proc Natl Acad Sci U S A 105, 18243–8 (2008).

11. Lechner, C. C., Agashe, N. D. & Fierz, B. Traceless Synthesis of Asymmetrically Modified Bivalent Nucleosomes. Angew Chemie Int Ed 55, 2903–2906 (2016).

12. Lukasak, B. J. et al. A Genetically Encoded Approach for Breaking Chromatin Symmetry. ACS Cent Sci 8, 176–183 (2022).

13. Mitchener, M. M. & Muir, T. W. Janus Bioparticles: Asymmetric Nucleosomes and Their Preparation Using Chemical Biology Approaches. Acc Chem Res 54, 3215– 3227 (2021).

14. Sidoli, S. & Garcia, B. A. Middle-Down Proteomics: A Still Unexploited Resource for Chromatin Biology. Expert Rev Proteomics 14, 617–626 (2017).

15. Sidoli, S. et al. Middle-down hybrid chromatography/tandem mass spectrometry workflow for characterization of combinatorial post-translational modifications in histones. Proteomics 14, 2200–2211 (2014).

16. Holt, M. V., Wang, T. & Young, N. L. Expeditious Extraction of Histones from Limited Cells or Tissue Samples and Quantitative Top-Down Proteomic Analysis. Curr Protoc 1, (2021).

17. Jain, K. et al. An acetylation-mediated chromatin switch governs H3K4 methylation readwrite capability. eLife 12, (2023).

18. Thompson, A. et al. Tandem Mass Tags: A Novel Quantification Strategy for Comparative Analysis of Complex Protein Mixtures by MS/MS. Anal Chem 75, 1895–1904 (2003).

19. Piotukh, K. et al. Directed Evolution of Sortase A Mutants with Altered Substrate Selectivity Profiles. J Am Chem Soc 133, 17536–17539 (2011).

20. Chen, I., Dorr, B. M. & Liu, D. R. A general strategy for the evolution of bond-forming enzymes using yeast display. Proc Natl Acad Sci U S A 108, 11399– 11404 (2011).

21. Zhulenkovs, D., Jaudzems, K., Zajakina, A. & Leonchiks, A. Enzymatic activity of circular sortase A under denaturing conditions: An advanced tool for protein ligation. Biochem Eng J 82, 200–209 (2014).

22. Musil, M. et al. FireProt: web server for automated design of thermostable proteins. Nucleic Acids Res 45, W393–W399 (2017).

23. Kalin, J. H. et al. Targeting the CoREST complex with dual histone deacetylase and demethylase inhibitors. Nat Commun 9, (2018).

24. Janssen, K. A., Coradin, M., Lu, C., Sidoli, S. & Garcia, B. A. Quantitation of Single and Combinatorial Histone Modifications by Integrated Chromatography of Bottom-up Peptides and Middle-down Polypeptide Tails. J Am Soc Mass Spectrom 30, 2449–2459 (2019).

25. Sidoli, S., Lin, S., Karch, K. R. & Garcia, B. A. Bottom-Up and Middle-Down Proteomics Have Comparable Accuracies in Defining Histone Post-Translational Modification Relative Abundance and Stoichiometry. Anal Chem 87, 3129–3133 (2015).

26. Shim, Y., Duan, M. R., Chen, X., Smerdon, M. J. & Min, J. H. Polycistronic coexpression and nondenaturing purification of histone octamers. Anal Biochem 427, 190–192 (2012).

27. Aparicio Pelaz, D., et al. Examining histone modification crosstalk using immobilized libraries established from ligation-ready nucleosomes. Chem Sci 11, 9218–9225 (2020).

28. Yang, Q., Gao, Y., Liu, X., Xiao, Y. & Wu, M. A General Method to Edit Histone H3 Modifications on Chromatin Via Sortase-Mediated Metathesis. Angew Chemie 134, e202209945 (2022).

29. Wang, Z. A. et al. Histone H2B Deacylation Selectivity: Exploring Chromatin’s Dark Matter with an Engineered Sortase. J Am Chem Soc 144, 3360–3364 (2022).

30. Luger, K., Mäder, A. W., Richmond, R. K., Sargent, D. F. & Richmond, T. J. Crystal structure of the nucleosome core particle at 2.8 Å resolution. Nature 389, 251– 260 (1997).

31. Wang, Z. A. et al. Diverse nucleosome site-selectivity among histone deacetylase complexes. eLife 9, e57663 (2020).

32. Wang, Z. et al. Structural Basis of Sirtuin 6-Catalyzed Nucleosome Deacetylation. J Am Chem Soc 145, 6811–6822 (2023).

33. Wang, Z. A. et al. Structural Basis of Sirtuin 6-Catalyzed Nucleosome Deacetylation. J Am Chem Soc 145, 6811–6822 (2023).

34. Lee, K. et al. Uncoupling Histone Modification Crosstalk by Engineering Lysine Demethylase LSD1. Nat Chem Biol (2024).

35. Turnbull, R. E. et al. The MiDAC histone deacetylase complex is essential for embryonic development and has a unique multivalent structure. Nat Commun 11, 1–15 (2020).

36. Song, Y. et al. Mechanism of Crosstalk between the LSD1 Demethylase and HDAC1 Deacetylase in the CoREST Complex. Cell Rep 30, 2699–2711 (2020).

37. Gates, L. A. et al. Histone butyrylation in the mouse intestine is mediated by the microbiota and associated with regulation of gene expression. Nat Metab 6, 697– 707 (2024).

38. Weinert, B. T. et al. Time-Resolved Analysis Reveals Rapid Dynamics and Broad Scope of the CBP/p300 Acetylome. Cell 174, 231–244.e12 (2018).

39. Stützer, A. et al. Modulations of DNA Contacts by Linker Histones and Post-translational Modifications Determine the Mobility and Modifiability of Nucleosomal H3 Tails. Mol Cell 61, 247–259 (2016).

40. Hao, F. et al. Acetylation-modulated communication between the H3 N-terminal tail domain and the intrinsically disordered H1 C-terminal domain. Nucleic Acids Res 48, 11510–11520 (2020).

41. Oishi, T. et al. Contributions of histone tail clipping and acetylation in nucleosome transcription by RNA polymerase II. Nucleic Acids Res 51, 10364–10374 (2023).

42. Skrajna, A. et al. Comprehensive nucleosome interactome screen establishes fundamental principles of nucleosome binding. Nucleic Acids Res 48, 9415–9432 (2020).

43. Wu, M. et al. Lysine-14 acetylation of histone H3 in chromatin confers resistance to the deacetylase and demethylase activities of an epigenetic silencing complex. eLife 7, 1–19 (2018).

44. Ishiyama, S. et al. Structure of the Dnmt1 Reader Module Complexed with a Unique Two-Mono-Ubiquitin Mark on Histone H3 Reveals the Basis for DNA Methylation Maintenance. Mol Cell 68, 350–360.e7 (2017).

45. Ren, W. et al. Direct readout of heterochromatic H3K9me3 regulates DNMT1-mediated maintenance DNA methylation. Proc Natl Acad Sci U S A 117, 18439– 18447 (2020).

46. Li, Z. et al. The expedient, CAET-assisted synthesis of dual-monoubiquitinated histone H3 enables evaluation of its interaction with DNMT1. Chem Sci 14, 5681– 5688 (2023).

47. Dhall, A., Weller, C. E., Chu, A., Shelton, P. M. M. & Chatterjee, C. Chemically Sumoylated Histone H4 Stimulates Intranucleosomal Demethylation by the LSD1-CoREST Complex. ACS Chem Biol 12, 2275–2280 (2017).

48. Hsu, P. L. et al. Structural Basis of H2B Ubiquitination-Dependent H3K4 Methylation by COMPASS. Mol Cell 76, 712–723.e4 (2019).

49. Currie, M. F., Singh, S. K., Ji, M. & Chatterjee, C. The semisynthesis of site-specifically modified histones and histone-based probes of chromatin-modifying enzymes. Methods 215, 28–37 (2023).

50. Li, W. et al. Rapid reconstitution of ubiquitinated nucleosome using a non-denatured histone octamer ubiquitylation approach. Cell Biosci 14, 1–15 (2024).

51. Shah, N. H., Dann, G. P., Vila-Perelló, M., Liu, Z. & Muir, T. W. Ultrafast protein splicing is common among cyanobacterial split inteins: Implications for protein engineering. J Am Chem Soc 134, 11338–11341 (2012).

52. Abeywardana, M. Y. et al. Multifaceted regulation of Sirtuin 2 (Sirt2) Deacetylase Activity. Journal of Biological Chemistry 0, 107722 (2024).

53. Yuan, Z. F. et al. Epiprofile quantifies histone peptides with modifications by extracting retention time and intensity in high-resolution mass spectra. Mol Cell Proteomics 14, 1696–1707 (2015).

54. Huang, Y. C. et al. Synthesis of l- and d-Ubiquitin by One-Pot Ligation and Metal-Free Desulfurization. Chem Eur J 22, 7623–7628 (2016).

55. Nodelman, I. M., Patel, A., Levendosky, R. F. & Bowman, G. D. Reconstitution and Purification of Nucleosomes with Recombinant Histones and Purified DNA. Curr Protoc Mol Biol 133, e130 (2020).

56. Worden, E. J., Zhang, X. & Wolberger, C. Structural basis for COMPASS recognition of an H2B-ubiquitinated nucleosome. eLife 9, (2020).

57. Li, X. et al. Electron counting and beam-induced motion correction enable near-atomic-resolution single-particle cryo-EM. Nat Methods 10, 584–590 (2013).

58. Zheng, S. Q. et al. MotionCor2: anisotropic correction of beam-induced motion for improved cryo-electron microscopy. Nat Methods 14, 331–332 (2017).

59. Punjani, A., Rubinstein, J. L., Fleet, D. J. & Brubaker, M. A. cryoSPARC: algorithms for rapid unsupervised cryo-EM structure determination. Nat Methods 14, 290–296 (2017).

60. Davey, C. A., Sargent, D. F., Luger, K., Maeder, A. W. & Richmond, T. J. Solvent Mediated Interactions in the Structure of the Nucleosome Core Particle at 1.9 Å Resolution. J Mol Biol 319, 1097–1113 (2002).

61. Pettersen, E. F. et al. UCSF Chimera—A visualization system for exploratory research and analysis. J Comput Chem 25, 1605–1612 (2004).

62. Vijay-Kumar, S., Bugg, C. E. & Cook, W. J. Structure of ubiquitin refined at 1.8 A resolution. J Mol Biol 194, 531–44 (1987).

