## Supplemental_Information for "A circular engineered sortase for interrogating histone H3 in chromatin"

**A circular engineered sortase for interrogating histone H3 in chromatin**
**Supplemental Information**

**Table of contents**

1. Materials
2. Antibodies
3. Cloning
4. Figure S1. Cleavage of purified, recombinant histone H3 by sortase mutants
5. Figure S2. Cleavage of semisynthetic, modified histone H3 by W11
6. Figure S3. SDS-PAGE analysis of sortase reaction in nuclear acid extracts
7. Figure S4. Sortase W11 mutant activity in the presence of co-solvents, chaotropes and detergents
8. Figure S5. Histone tails isolated by trichloroacetic acid precipitation of reaction protein components
9. Table S1. Abundance of individual H3 post-translational modifications in bottom-up and middle-down proteomics data
10. Figure S6. Characterization of H3(33-135) octamer overexpression and purification
11. Figure S7. Western blot analysis of cW11 nucleosome ligation
12. Figure S8. cW11 nucleosome ligation time course chromatograms
13. Figure S9. Mass spectrometric characterization of cW11 nucleosome ligation products
14. Figure S10. SDS-PAGE and TBE native gel characterization of cW11 nucleosome ligation products
15. Figure S11. Mass spectrometric characterization of peptide substrates used in cW11 nucleosome ligation
16. Figure S12. Characterization of asymmetric H3K4me2 H3K14ac ligation intermediate and final products
17. Figure S13. Comparison of Sirt6 deacetylase activity toward H3K9ac nucleosomes prepared by conventional reconstitution and cW11 nucleosome ligation
18. Figure S14. Comparison of LHC deacetylase activity toward H3K9ac nucleosomes prepared by conventional reconstitution and cW11 nucleosome ligation
19. Figure S15. Cryo-electron microscopy characterization of nucleosomes prepared by cW11 nucleosome ligation
20. Figure S16. Western blot measurement of Sirt1 activity by length of H3K9 acylation carbon chain.
21. Figure S17. Western blot measurement of Sirt2 activity by length of H3K9 acylation carbon chain.
22. Figure S18. Western blot measurement of Sirt6 activity by length of H3K9 acylation carbon chain.
23. Figure S19. Western blot measurement of MiDAC activity by length of H3K9 acylation carbon chain.
24. Table S6. Calculated V/[E] values for HDAC activity by length of H3K9 acylation carbon chain
25. Figure S20. Western blot measurement of Sirt1 activity toward four carbon acylations of H3K9
26. Figure S21. Western blot measurement of Sirt2 activity toward four carbon acylations of H3K9
27. Figure S22. Western blot measurement of Sirt6 activity toward four carbon acylations of H3K9
28. Figure S23. Western blot measurement of MiDAC activity toward four carbon acylations of H3K9
29. Figure S24. Western blot measurement of LHC activity toward four carbon acylations of H3K9
30. Figure S25. Western blot measurement of free HDAC activity toward four carbon acylations of H3K9
31. Table S7. Calculated V/[E] values for HDAC activity toward four carbon acylations of H3K9
32. Figure S26. Western blot measurement of Sirtuin5 activity toward acetylated and succinylated H3K9
33. Table S8. Calculated V/[E] values for HDAC activity toward H3K9 succinylation
34. Figure S27. Validation of H3Kac single site antibody specificity
35. Figure S28. Western blot measurement of Sirt2 activity toward mono-acetylated nucleosomes.
36. Figure S29. Western blot measurement of Sirt2 activity toward penta-acetylated nucleosomes.

37. Figure S30. Western blot measurement of Sirt6 activity toward mono- and penta-acetylated nucleosomes.
38. Figure S31. Western blot measurement of MiDAC activity toward mono- and penta-acetylated nucleosomes.
39. Figure S32. Western blot characterization of asymmetric mono/tetra-acetylated nucleosomes
40. Figure S33. Mass spectrometric characterization of asymmetric mono/tetra-acetylated nucleosomes and asymmetric mono-acetylated/unmodified nucleosomes.
41. Figure S34. Western blot measurement of Sirt2 activity toward asymmetrically acetylated nucleosomes
42. Figure S35. Western blot measurement of Sirt6 activity toward asymmetrically acetylated nucleosomes
43. Table S9. Calculated V/[E] values for Sirt2 activity toward mono-, penta-, and asymmetrically acetylated nucleosomes
44. Table S10. Calculated V/[E] values for Sirt6 activity toward mono-, penta-, and asymmetrically acetylated nucleosomes
45. Figure S36. Western blot measurement of MiDAC activity toward asymmetrically acetylated nucleosomes
46. Table S11. Calculated V/[E] values for MiDAC activity toward mono-, penta-, and asymmetrically acetylated nucleosomes
47. Figure S37. Western blot measurement of LC activity toward nucleosomes with asymmetric methylation and acetylation
48. Figure S38. Mass spectrometric characterization of ubiquitinated H3 peptides and ubiquitinated nucleosome
49. Figure S39. Western blot characterization of asymmetric unmodified/ubiquitinated nucleosome
50. Figure S40. Electrophoretic mobility shift assay titrating asymmetric nucleosomes with sfGFP-RFTS fusion
51. Figure S41. Electrophoretic mobility shift assay titrating sfGFP-RFTS fusion with asymmetric nucleosomes
52. References

**Materials**

Chemicals were purchased from Sigma-Aldrich, Fisher Scientific, Chem-Impex, Research Products International, or Ambeed and used without further purification. Fmoc protected amino acids were purchased from Sigma-Aldrich, Chem-Impex, Ambeed, P3 biosystems or ApexBio. Solvents were purchased from Sigma-Aldrich, Fisher Scientific, of EMD-Millipore. HDAC inhibitors MS-275 and Corin were purchased from MedChemExpress. Tandem mass tag reagents were provided by Thermo Scientific as part of the Tandem Mass Tag Systems Research Award. Media for bacterial culture was purchased from Sigma-Aldrich. Media for mammalian cell culture was purchased from Caisson Labs. Restriction enzymes were purchased from New England Biosciences. DNA isolation kits were purchased from Zymo. Polymerases were purchased from Agilent Technologies, Promega or New England Biosciences. Primers were purchased from Integrated DNA Technologies or Quintara Biosciences. Sanger sequencing was performed by Integrated DNA Technologies or Quintara Biosciences. Whole plasmid sequencing was performed by Quintara Biosciences or Plasmidsaurus. Columns for HPLC were purchased from Tosoh Biosciences (DEAE), Agilent technologies (C18), or Higgins Analytical (C18). Columns for FPLC were purchased from Cytiva. Automated solid-phase peptide synthesizer was purchased from Gyros Protein Technologies (Prelude).

**Antibodies**

For demethylase assays, anti-H3K4me2 (1:2000 dilution, Abcam, cat ab32356) and anti-H3 (1:2000 dilution, Abcam, cat ab1791) primary antibodies were used. For deacetylase assays, anti-H3K9ac (1:2000 dilution, Abcam AB32129), anti-H3K14ac (1:2000 dilution, EMD Millipore 07-353), anti-H3K18ac (1:2000 dilution, EMD Millipore 07-354), anti-H3K23ac (1:2000 dilution, EMD Millipore 07-355), anti-H3K27ac (1:2000 dilution, Cell Signaling 81735), anti-H3K9bu (1:2000 dilution, PTM Biolabs, PTM-305 need double check), anti-H3K9cro (1:2000 dilution, PTM Biolabs, PTM-539), pan anti-Khib (1:2000 dilution, A kind gift from Dr. Yingming Zhao, University of Chicago), anti-H3K9succ (1:1000 dilution, A kind gift from Dr. Yingming Zhao, University of Chicago), and anti-H3 (Abcam, #ab1791, 1:2000 dilution) primary antibodies were used. For all assays anti-rabbit IgG (1:1000 dilution, Cell Signaling Technology, cat 7074S) or anti-mouse IgG (1:1000 dilution, Cell Signaling Technology, cat 7076S) secondary antibodies were used.

**Cloning**

Sortase mutants were derived from previously reported variants of S. aureus sortase A (Srt A): F40 sortase mutant, enhanced sortase pentamutant (eSrt), and cyclic sortase.[1–3] Additional mutations were identified using FireProt webserver.[4] DNA encoding S. aureus Srt A (60-206) (A kind gift of Dr. Dirk Schwarzer) was cloned into pET21 vector (Ampicillin resistant). Subsequent mutations were accomplished by sequential rounds of quick-change site directed mutagenesis. The PCR products were purified by PCR cleanup kit (Zymo), treated with DpnI (New England Biolabs), and transformed into DH5anti E. coli. Single colonies were picked and grown over night in 5 mL of Luria Bertani media supplemented with ampicillin (100 µg / mL), then pelleted (5 minutes, 4500 rcf, 4 ˚C). Plasmids were obtained from cell pellets by mini-prep (Plasmid Miniprep Classic, Zymo) and Sanger sequenced, or sequenced by nano-pore.

*Sortase F40* (mutations distinguishing F40 from Srt A are underlined)
MQAKPQIPKDKSKVAGYIEIPDADIKEPVYPGPATPEQLNRGVSFAEENESLDDQNISIAGHTFIDRPNYQF
TNLKAAKMGSMVYFKVGNETRKYKMTSIRDVKPQDVGMHLAEKGKDKQLTLITCDDYNEKTGVWEKRKI
FVATEVKLEHHHHHH

*Sortase W6* (mutations distinguishing F40 from Srt A are underlined, mutations from eSrt are highlighted)
MQAKPQIPKDKSKVAGYIEIPDADIKEPVYPGPATSEQLNRGVSFAEENESLDDQNISIAGHTFIDRPNYQF
TNLKAAKMGSMVYFKVGNETRKYKMTSIRNVKPTAVMHLAEKGKDKQLTLITCDDYNEKTGVWEKRKIFV
ATEVKLEHHHHHH

*Sortase W8* (mutations distinguishing F40 from Srt A are underlined, mutations from eSrt are highlighted)
MQAKPQIPKDKSKVAGYIEIPDADIKEPVYPGPATPEQLNRGVSFAEENESLDDQNISIAGHTFIDRPNYQF
TNLKAAKMGSMVYFKVGNETRKYKMTSIRNVKPQDVMHLAEKGKDKQLTLITCDDYNEETGVWETRKIF
VATEVKLEHHHHHH

*Sortase W9* (mutations distinguishing F40 from Srt A are underlined, mutations from eSrt are highlighted)
MQAKPQIPKDKSKVAGYIEIPDADIKEPVYPGPATSEQLNRGVSFAEENESLDDQNISIAGHTFIDRPNYQF
TNLKAAKMGSMVYFKVGNETRKYKMTSIRNVKPQDVMHLAEKGKDKQLTLITCDDYNEKTGVWETRKIF
VATEVKLEHHHHHH

*Sortase W11* (mutations distinguishing F40 from Srt A are underlined, mutations from eSrt are highlighted)
MQAKPQIPKDKSKVAGYIEIPDADIKEPVYPGPATSEQLNRGVSFAEENESLDDQNISIAGHTFIDRPNYQF
TNLKAAKMGSMVYFKVGNETRKYKMTSIRNVKPQDVMHLAEKGKDKQLTLITCDDYNEETGVWETRKIF
VATEVKLEHHHHHH

*Sortase W12* (mutations distinguishing F40 from Srt A are underlined, mutations from eSrt are highlighted)
MQAKPQIPKDKSKVAGYIEIPDADIKEPVYPGPATSEQLNRGVSFAEENESLDDQNISIAGHTFIDRPNYQF
TNLKAAKMGSMVYFKVGNETRKYKMTSIRNVKPTAVMHLAEKGKDKQLTLITCDDYNEKTGVWETRKIFV
ATEVKLEHHHHHH

*Sortase W13* (mutations distinguishing F40 from Srt A are underlined, mutations from eSrt are highlighted)
MQAKPQIPKDKSKVAGYIEIPDADIKEPVYPGPATSEQLNRGVSFAEENESLDDQNISIAGHTFIDRPNYQF
TNLKAAKMGSMVYFKVGNETRKYKMTSIRNVKPQAVMHLAEKGKDKQLTLITCDDYNEETGVWETRKIFV
ATEVKLEHHHHHH

*Sortase W15* (mutations distinguishing F40 from Srt A are underlined, mutations from eSrt are highlighted)
MQAKPQIPKDKSKVAGYIEIPDADIKEPVYPGPATREQLNRGVSFAEENESLDDQNISIAGHTFIDRPNYQF
TNLKAAKMGSMVYFKVGNETRKYKMTSIRNVKPQDVMHLAEKGKDKQLTLITCDDYNEETGVWETRKIF
VATEVKLEHHHHHH

*Sortase W16* (mutations distinguishing F40 from Srt A are underlined, mutations from eSrt are highlighted)
MQAKPQIPKDKSKVAGYIEIPDADIKEPVYPGPATREQLNRGVSFAEENESLDDQNISIAGHTFIDRPNYQF
TNLKAAKMGSMVYFKVGNETRKYKMTSIRNVKPQAVMHLAEKGKDKQLTLITCDDYNEKTGVWETRKIFV
ATEVKLEHHHHHH

*Sortase W11 E108Q S116V* (mutations distinguishing F40 from Srt A are underlined, mutations from eSrt are highlighted, mutations from FireProt are **bolded**)
MQAKPQIPKDKSKVAGYIEIPDADIKEPVYPGPATSEQLNRGVSFAEEN**Q**SLDDQNI**V**IAGHTFIDRPNYQF
TNLKAAKMGSMVYFKVGNETRKYKMTSIRNVKPQDVMHLAEKGKDKQLTLITCDDYNEETGVWETRKIF
VATEVKLEHHHHHH

*Sortase cW11* (Split intein is *italicized*, linkers are **bolded** and W11 E108Q S116V is underlined)
*MIKIATRKYLGKQNVYDIGVERDHNFALKNGFIASN***CFNGGSS**MQAKPQIPKDKSKVAGYIEIPDADIKEPV
YPGPATSEQLNRGVSFAEENQSLDDQNIVIAGHTFIDRPNYQFTNLKAAKMGSMVYFKVGNETRKYKMTS
IRNVKPQDVMHLAEKGKDKQLTLITCDDYNEETGVWETRKIFVATEVKLEHHHHHH**AEY***CLSYETEILTVE*
*YGLLPIGKIVEKRIECTVYSVDNNGNIYTQPVAQWHDRGEQEVFEYCLEDGSLIRATKDHKFMTVDGQML*
*PIDEIFERELDLMRVDNLPN*

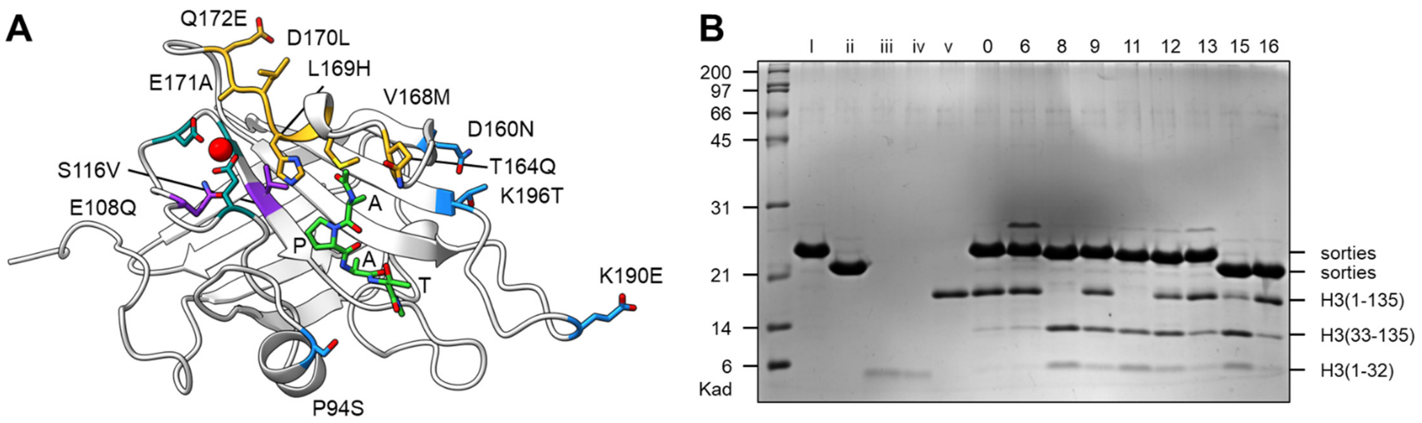

**C**

| # | P94 | D160 | T164 | D165 | K190 | K196 | cleavage yield |
|---|---|---|---|---|---|---|---|
| 0 | - | - | Q | - | - | - | **4%** |
| 6 | S | N | - | A | - | - | **7%** |
| 8 | - | N | Q | - | E | T | **79%** |
| 9 | S | N | Q | - | - | T | **29%** |
| 11 | S | N | Q | - | E | T | **82%** |
| 12 | S | N | - | A | - | T | **39%** |
| 13 | S | N | Q | A | E | T | **16%** |
| 15 | R | N | Q | - | E | T | **55%** |
| 16 | R | N | Q | - | - | T | **20%** |

**Figure S1. Cleavage of purified, recombinant histone H3 by sortase mutants.** (**A**) Sites of mutations found in W11 sortase, with origin indicated by color: F40 (yellow); enhanced sortase (blue); FireProt (purple). Calcium (red) binding residues are depicted in dark green, and bound substrate is depicted in light green. Mutations introduced in pymol (PDBID: 2KID), followed by Rosetta energy minimization. (**B**) SDS-PAGE of sortase cleavage reaction with recombinant, unmodified histone H3: i & ii – sortase mutant standards; iii & iv – H3 tail peptide standards; v – H3 protein standard; 0 – F40 sortase; 6-16 – F40-derived mutants. (**C**) Specific enhanced sortase mutations introduced to F40 in mutants W6 through W16 and resultant H3 tail cleavage yields.

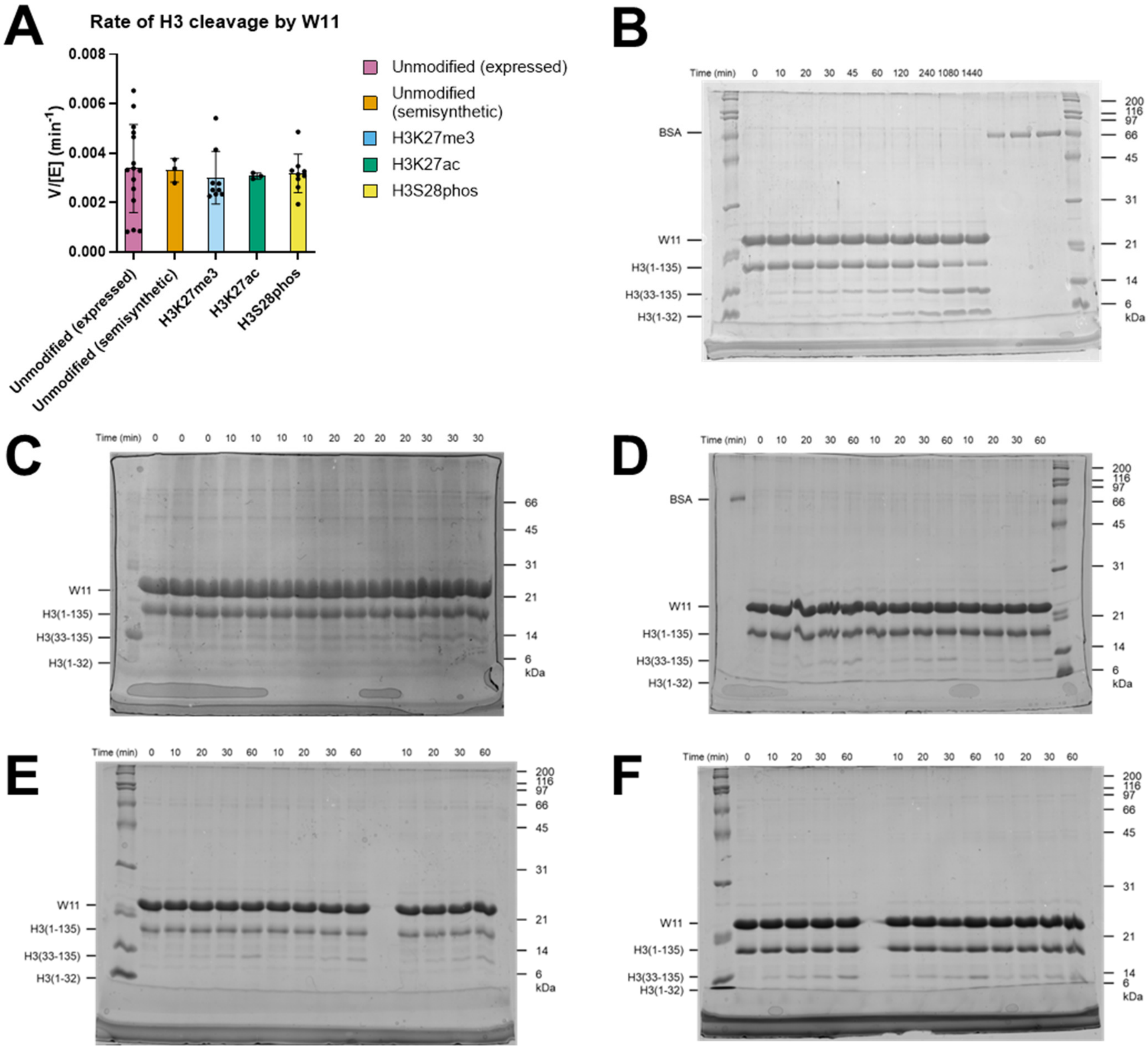

**Figure S2. Cleavage of semisynthetic, modified histone H3 by W11.** (**A**) Rates of H3 cleavage by W11 sortase in the presence of post-translational modifications near the sortase motif (A29-G33). Rates were determined by densitometry (ImageJ) from SDS-PAGE analysis of starting material, H3(1-135), disappearance and product, H3(33-135), formation over time. Data was fitted to a non-linear one-phase decay using GraphPad Prism 10.2.3. (**B-F**) Representative SDS-PAGE gels used for evaluating W11 sortase cleavage rate: (**B**) unmodified, heterologously expressed H3; (**C**) unmodified, semisynthetic H3; (**D**) semisynthetic H3K27me3; (**E**) semisynthetic H3K27ac; (**F**) semisynthetic H3S28phos.
30 μM H3, 100 μM sortase, 1 mM GGG peptide, 20 mM PIPES (pH 7), 1 mM CaCl2, 1 mM DTT

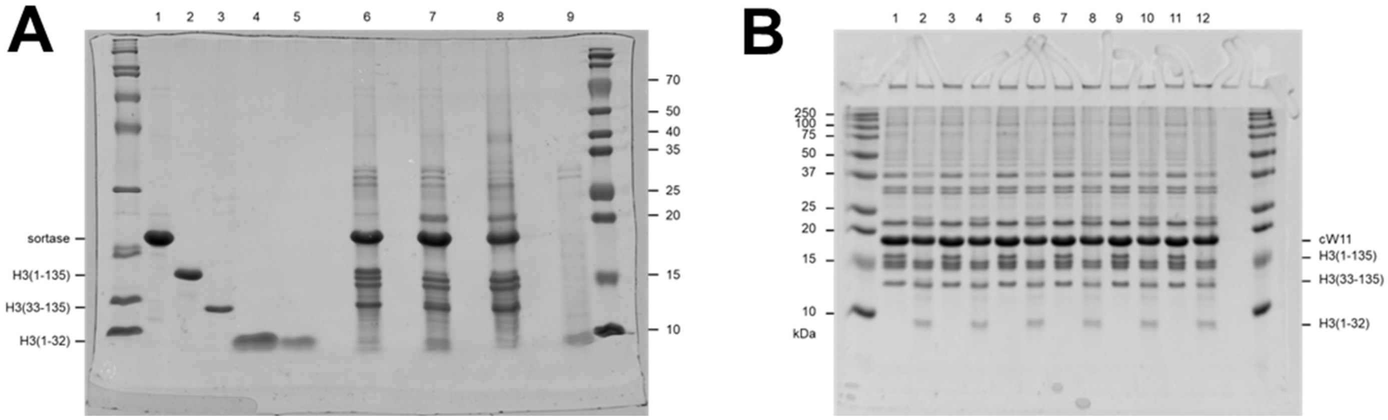

**Figure S3. SDS-PAGE analysis of sortase W11 & cW11 reactions in nuclear acid extracts.** (**A**) W11 reaction with standards: lane 1 – W11; lane 2 – H3(1-135); lane 3 – H3(33-135); lane 4 & 5 – H3(1-32) standard; lane 6 – 0 hr acid extract reaction; lane 7 – 16 hr acid extract reaction; lane 8 – TCA pelleted protein from 16 hr acid extract reaction; lane 9 – buffer exchanged supernatant from TCA precipitation. (**B**) Representative replicate acid extract reactions with cW11: lane 1 – 0 hr acid extract reaction replicate 1; lane 2 – 16 hr acid extract reaction replicate 1; lane 3 – 0 hr acid extract reaction replicate 2; lane 4 – 16 hr acid extract reaction replicate 2; lane 5 – 0 hr acid extract reaction replicate 3; lane 6 – 16 hr acid extract reaction replicate 3; lane 7 – 0 hr acid extract reaction replicate 4; lane 8 – 16 hr acid extract reaction replicate 4; lane 9 – 0 hr acid extract reaction replicate 5; lane 10 – 16 hr acid extract reaction replicate 5; lane 11 – 0 hr acid extract reaction replicate 6; lane 12 – 16 hr acid extract reaction replicate 6. Lanes 1 & 2 reprinted from main text figure 1.

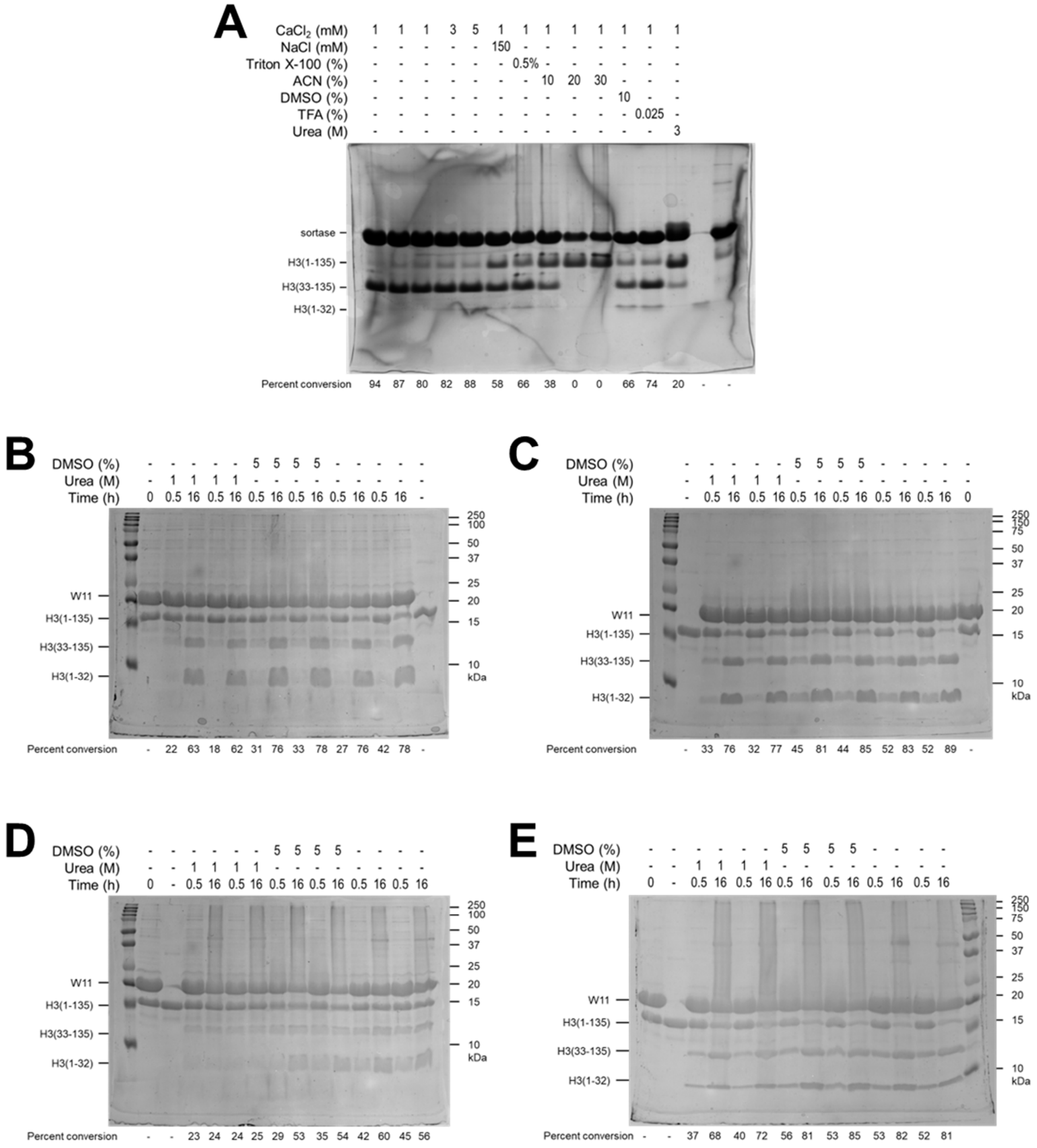

**Figure S4. Sortase W11 mutant activity in the presence of co-solvent, detergent or chaotrope.** (**A**) Endpoint SDS-PAGE analysis of 25 °C sortase W11 reaction run with variant buffer conditions including increased salt, added detergent, organic co-solvent, or chaotrope. (**B** & **C**) Endpoint SDS-PAGE analysis of duplicate 25 °C sortase reaction with W11 (**B**) or FireProt mutant W11(E108Q, S116V) (**C**). (**D** & **E**) Endpoint SDS-PAGE analysis of duplicate 37 °C sortase reaction with W11 (**D**) or FireProt mutant W11(E108Q, S116V) (**E**). In all cases cleavage of H3 was evaluated by densitometry (ImageJ), and percent conversion was calculated as the sum of H3(33-135) and H3(1-32) over the cumulative intensity of all H3-derived bands.

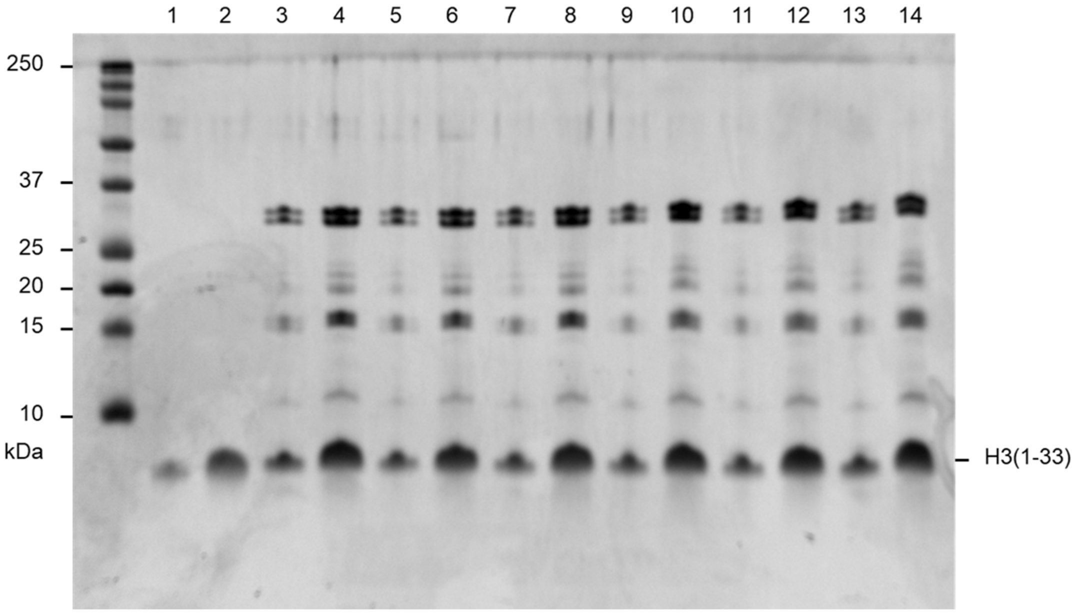

**Figure S5. Histone tails isolated by trichloroacetic acid precipitation of reaction protein components.** Representative histone tails isolated by TCA precipitation from the sortase reaction conducted in nuclear acid extracts. Samples are derived from reactions shown in figure S3b, and a portion (lanes 1-3) of this gel is re-printed in main figure 1. Representative TCA products from replicate acid extract reactions with cW11: lane 1 & 2 – H3(1-32) peptide standards for quantification; lane 3 & 4 – replicate 1 product; lane 5 & 6 – replicate 2 product; lane 7 & 8 – replicate 3 product; lane 9 & 10 – replicate 4 product; lane 11 & 12 – replicate 5 product; lane 13 & 14 – replicate 6 product.

| | Middle-down | | Bottom-up | | | Middle-down | | Bottom-up | |
|---|---|---|---|---|---|---|---|---|---|
| PTM | DMSO & Corin H3.1 & H3.3 | DMSO & MS275 H3.1 & H3.3 | DMSO & Corin H3.1 & H3.3 | DMSO & MS275 H3.1 & H3.3 | PTM | DMSO & Corin H3.1 & H3.3 | DMSO & MS275 H3.1 & H3.3 | DMSO & Corin H3.1 & H3.3 | DMSO & MS275 H3.1 & H3.3 |
| unmodified | 0.02% | 0.10% | 0.00% | 0.00% | K18me1 | 0.08% | 0.10% | 0.01% | 0.00% |
| R2me1 | 0.00% | 0.00% | 0.00% | 0.00% | K18me2 | 0.05% | 0.00% | 1.34% | 1.76% |
| R2me2 | 0.00% | 0.00% | 0.00% | 0.00% | K18me3 | 0.00% | 0.00% | 0.04% | 0.05% |
| T3phos | 0.00% | 0.00% | 0.00% | 0.00% | K18ac | 1.31% | 1.55% | 0.00% | 0.00% |
| K4me1 | 0.21% | 0.05% | 24.06% | 24.24% | K18prop | 0.23% | 0.15% | 0.00% | 0.00% |
| K4me2 | 0.00% | 0.00% | 0.00% | 0.00% | T22phos | 11.18% | 14.69% | 0.00% | 0.00% |
| K4me3 | 0.00% | 0.00% | 0.00% | 0.00% | K23me1 | 0.34% | 0.97% | 0.00% | 0.00% |
| K4ac | 0.00% | 0.00% | 0.00% | 0.00% | K23me2 | 0.41% | 0.37% | 0.00% | 0.00% |
| K4prop | 0.00% | 0.00% | 0.00% | 0.00% | K23me3 | 0.00% | 0.05% | 0.00% | 0.00% |
| T6phos | 0.00% | 0.00% | 0.00% | 0.00% | K23ac | 66.14% | 66.15% | 54.89% | 57.79% |
| R8me1 | 0.20% | 0.00% | 0.00% | 0.00% | K23prop | 4.07% | 3.99% | 0.00% | 0.00% |
| R8me2 | 0.12% | 0.24% | 0.00% | 0.00% | R26me1 | 7.19% | 5.83% | 0.00% | 0.00% |
| K9me1 | 12.84% | 11.10% | 34.21% | 30.82% | R26me2 | 0.94% | 0.72% | 0.00% | 0.00% |
| K9me2 | 32.04% | 32.33% | 17.41% | 15.88% | H3.1_K27me1 | 7.57% | 7.94% | 23.45% | 23.45% |
| K9me3 | 6.40% | 5.91% | 8.81% | 8.04% | H3.1_K27me2 | 67.70% | 68.95% | 44.06% | 44.12% |
| K9ac | 6.48% | 7.63% | 3.54% | 3.54% | H3.1_K27me3 | 0.15% | 0.13% | 14.58% | 14.88% |
| K9prop | 0.84% | 0.81% | 0.00% | 0.00% | H3.1_K27ac | 1.56% | 0.90% | 0.03% | 0.03% |
| S10phos | 0.00% | 0.05% | 0.00% | 0.00% | H3.1_K27prop | 0.20% | 0.14% | 0.00% | 0.00% |
| T11phos | 0.16% | 0.11% | 0.00% | 0.00% | H3.3_K27me1 | 0.00% | 0.00% | 31.08% | 31.16% |
| K14me1 | 0.47% | 0.20% | 0.00% | 0.00% | H3.3_K27me2 | 9.01% | 47.55% | 35.06% | 34.61% |
| K14me2 | 0.11% | 0.15% | 0.00% | 0.00% | H3.3_K27me3 | 0.00% | 0.00% | 14.64% | 14.31% |
| K14me3 | 0.00% | 0.13% | 0.00% | 0.00% | H3.3_K27ac | 0.00% | 0.00% | 0.00% | 0.00% |
| K14ac | 71.65% | 70.81% | 31.28% | 31.15% | H3.3_K27prop | 2.90% | 0.00% | 0.00% | 0.00% |
| K14prop | 1.61% | 2.50% | 0.00% | 0.00% | S28phos | 0.24% | 0.16% | 0.00% | 0.00% |
| R17me1 | 0.06% | 0.28% | 0.00% | 0.00% | S31phos | 0.00% | 0.00% | 0.00% | 0.00% |
| R17me2 | 0.00% | 0.04% | 0.00% | 0.00% | T32phos | 0.00% | 0.00% | 0.00% | 0.00% |
| | Pearson correlation | | | | | | | | |
| middle-down v. bottom-up | | | 75.01% | 80.17% | | | | | |

**Table S1. Abundance of individual H3 post-translational modifications in bottom-up and middle-down proteomics data.**

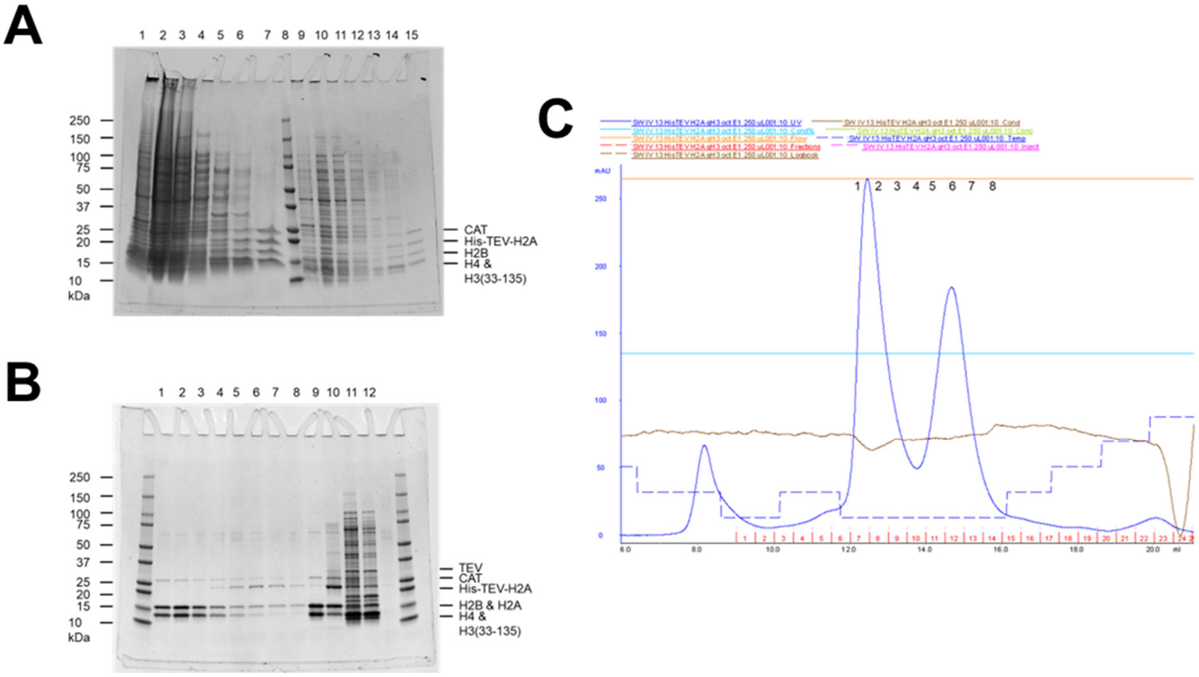

**Figure S6. Characterization of H3(33-135) octamer overexpression and purification.** (**A**) SDS-PAGE evaluation of fractions from IMAC purification of co-overexpressed His-TEV-H2A, H2B, H3(33-135) and H4. Lanes: 1 – lysate pellet; 2 – lysate supernatant; 3 – IMAC column flow through; 4 – lysis buffer wash; 5 – 20 mM imidazole wash 1; 6 – 20 mM imidazole wash 2; 7 – 200 mM imidazole elution; 8 – ladder; 9 – lysate pellet; 10 – lysate supernatant; 11 – IMAC column flow through; 12 – lysis buffer wash; 13 – 20 mM imidazole wash 1; 14 – 20 mM imidazole wash 2; 15 – 200 mM imidazole elution. CAT – chloramphenicol acetyltransferase. (**B**) SDS-PAGE evaluation of TEV cleavage of His-TEV-H2A octamer, and superdex200 fractions from post-TEV purification. Lanes: 1-8 – sequential superdex200 fractions (see **C**); 9 – TEV cleavage endpoint; 10 – TEV cleavage starting point; 11 – IMAC 20 mM imidazole wash 1; 12 – IMAC 20 mM imidazole wash 2. (**C**) Superdex200 chromatogram from purification following TEV cleavage of His-TEV-H2A octamer. Numbering (top) reflects corresponds to gel lanes in figure S6B.

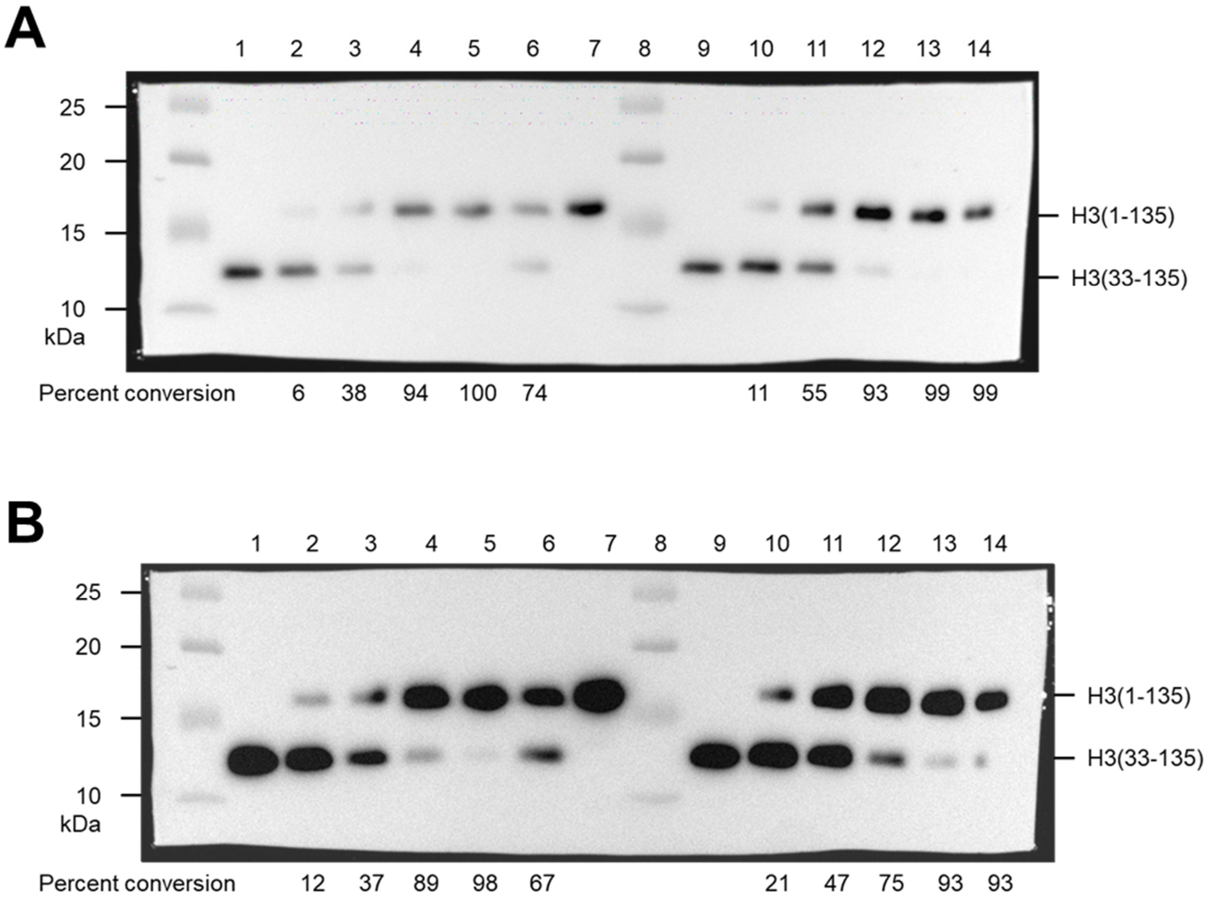

**Figure S7. Western blot analysis of cW11 nucleosome ligation.** (**A**) Anti-H3 western blot (1 minute exposure) of cW11 sortase ligation time-course replicates and DEAE fractions from an un-optimized gradient. Lanes: 1 – H3(33-135) standard; 2 – 0.5 hours; 3 – 1 hour; 4 – 4 hours; 5 – DEAE first peak; 6 – DEAE second peak; 7 – H3(1-135) standard; 8 – ladder; 9 – H3(33-135) standard; 10 – 0.5 hours; 11 – 1 hour; 12 – 4 hours; 13 – DEAE first peak; 14 – DEAE second peak. (**B**) Long exposure (10 minutes) of western blot from **A**. Percent conversion was assessed by densitometry using ImageJ.

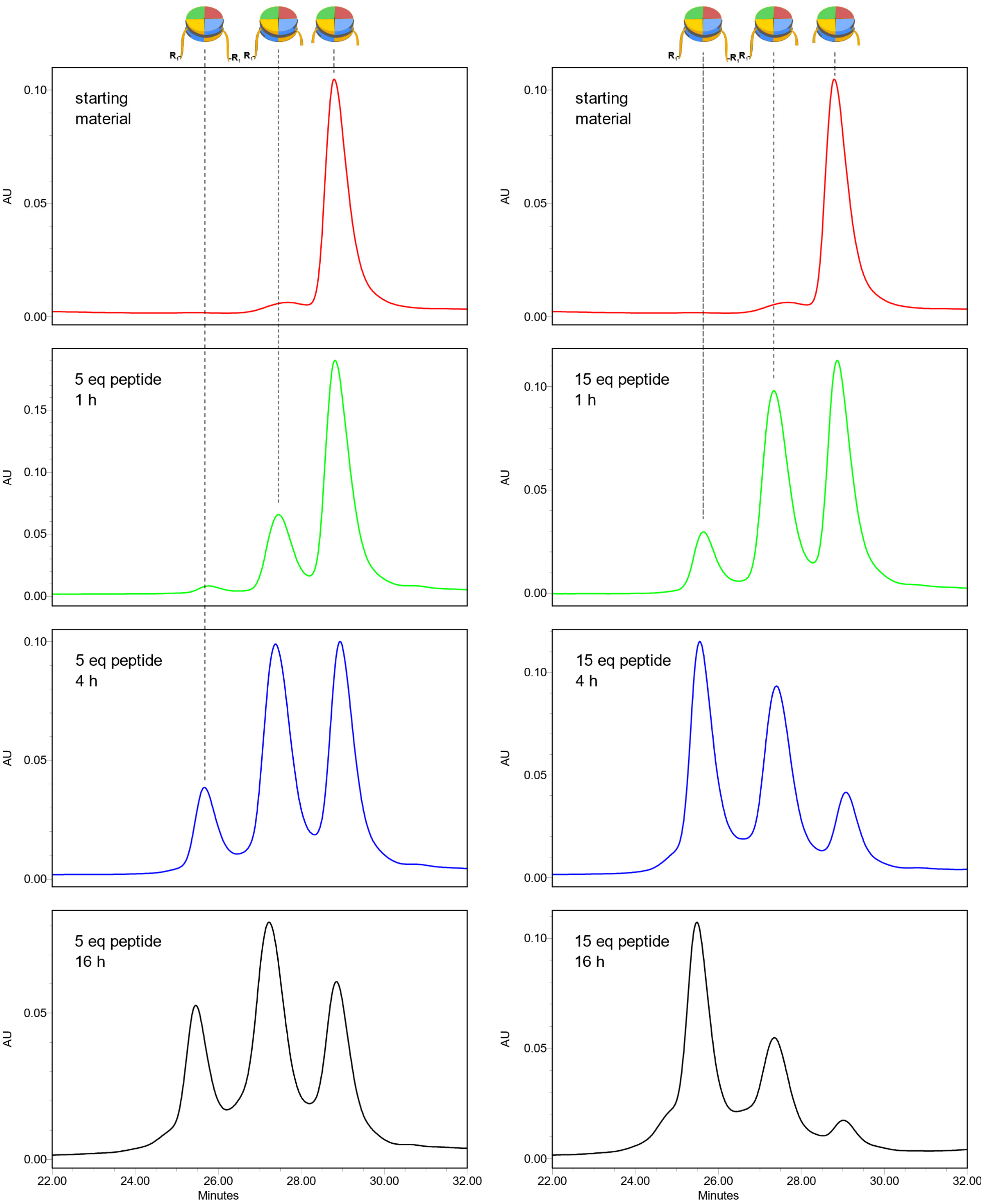

**Figure S8. cW11 nucleosome ligation time course chromatograms.** Change in product distribution over the course of the cW11 sortase ligation using either 5 equivalents of H3 tail peptide (**left**) or 15 equivalents (**right**). Peaks correspond to nucleosome with two copies of full length H3 (**left**), one copy of full length H3 and one copy of N-terminally truncated H3 (aa33-135) (**middle**), and starting material nucleosome with two copies of truncated H3 (aa33-135) (**right**).

**Figure S9. Mass spectrometric characterization of cW11 nucleosome ligation products.** Raw spectra deconvoluted with UniDec.[5]

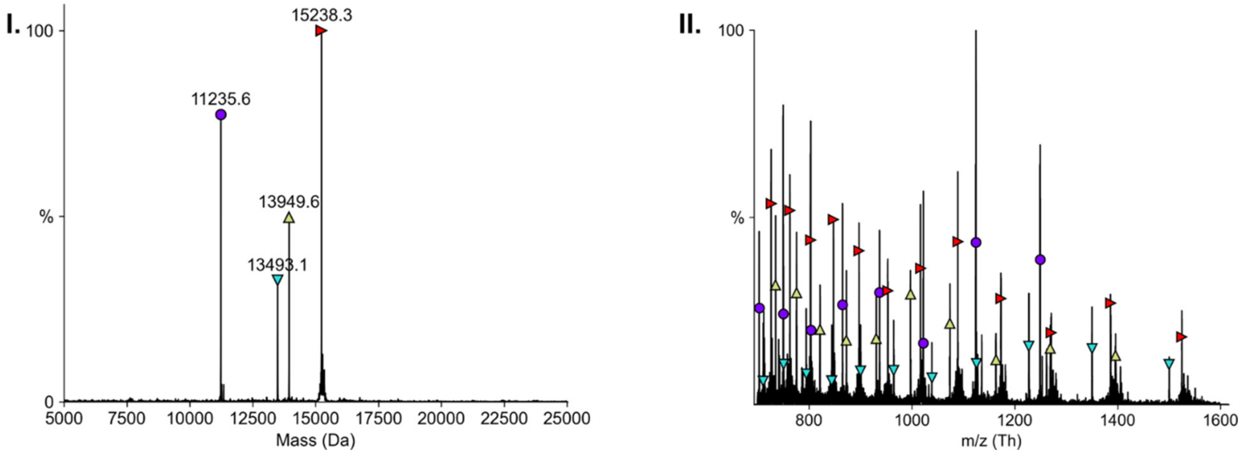

**A. 147 bp unmodified nucleosome.** (**I**) Deconvoluted mass spectrum: H4 (purple circle) calculated mass 11236.15 Da, found: 11235.6 Da; H2B (teal downward pointed triangle) calculated mass 13493.68 Da, found: 13493.2 Da; H2A (green upward pointed triangle) calculated mass 13950.2 Da, found: 13949.6 Da; unmodified H3 (red rightward pointed triangle) calculated mass 15238.61 Da, found: 15238.2 Da. (**II**) Raw mass spectrum: H4 (purple circle); H2B (teal downward pointed triangle); H2A (green upward pointed triangle; H3 (red rightward pointed triangle).

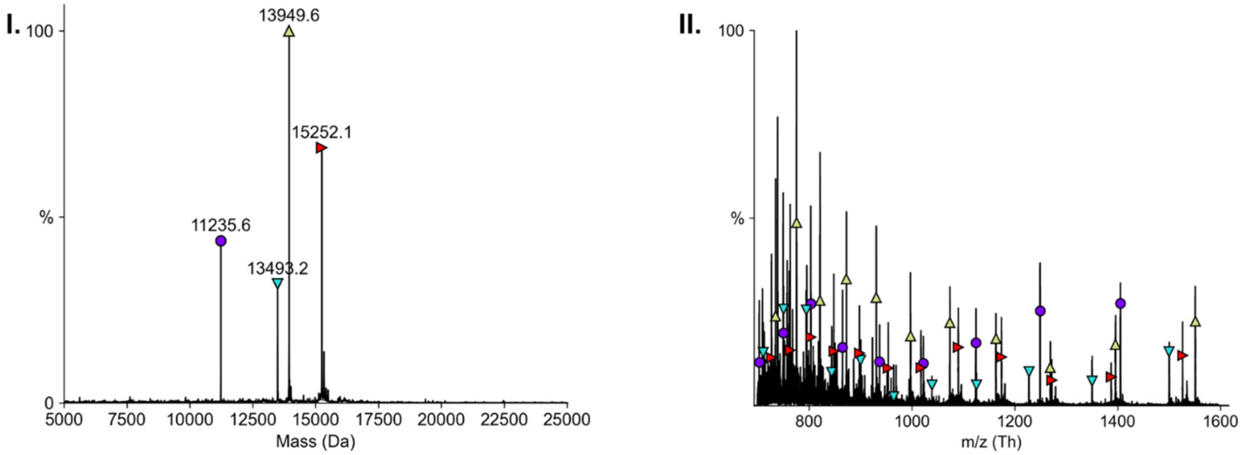

**B. 147 bp H3K4-monomethyl nucleosome.** (**I**) Deconvoluted mass spectrum: H4 (purple circle) calculated mass 11236.15 Da, found: 11235.6 Da; H2B (teal downward pointed triangle) calculated mass 13493.68 Da, found: 13493.2 Da; H2A (green upward pointed triangle) calculated mass 13950.2 Da, found: 13949.6 Da; H3K4me1 (red rightward pointed triangle) calculated mass 15251.63 Da, found: 15252.1 Da. (**II**) Raw mass spectrum: H4 (purple circle); H2B (teal downward pointed triangle); H2A (green upward pointed triangle; H3K4me1 (red rightward pointed triangle).

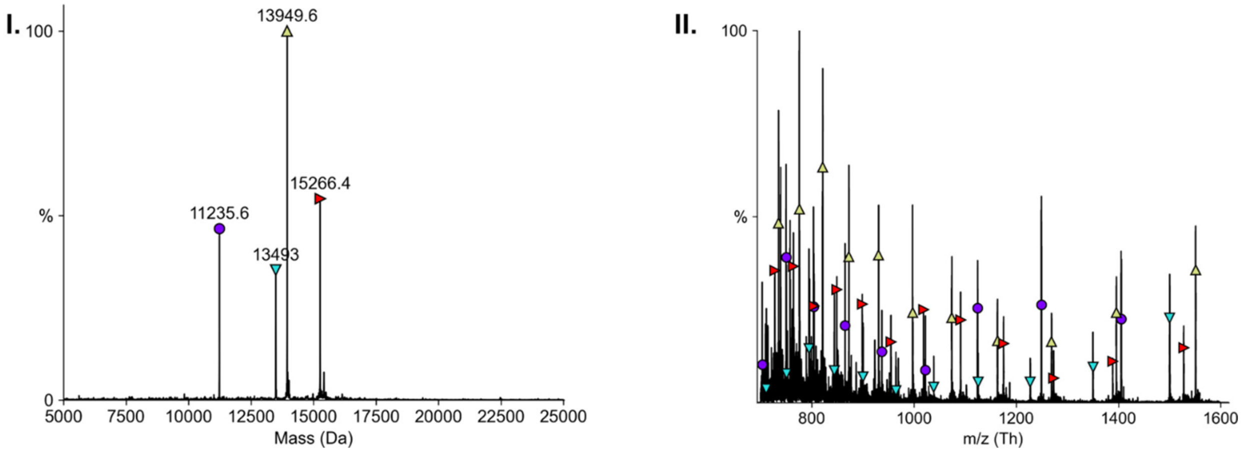

**C. 147 bp H3K4-dimethyl I nucleosome.** (**I**) Deconvoluted mass spectrum: H4 (purple circle) calculated mass 11236.15 Da, found: 11235.4 Da; H2B (teal downward pointed triangle) calculated mass 13493.68 Da, found: 13493.0 Da; H2A (green upward pointed triangle) calculated mass 13950.2 Da, found: 13949.5 Da; H3K4me2 (red rightward pointed triangle) calculated mass 15266.66 Da, found: 15266.4 Da. (**II**) Raw mass spectrum: H4 (purple circle); H2B (teal downward pointed triangle); H2A (green upward pointed triangle; H3K4me2 (red rightward pointed triangle).

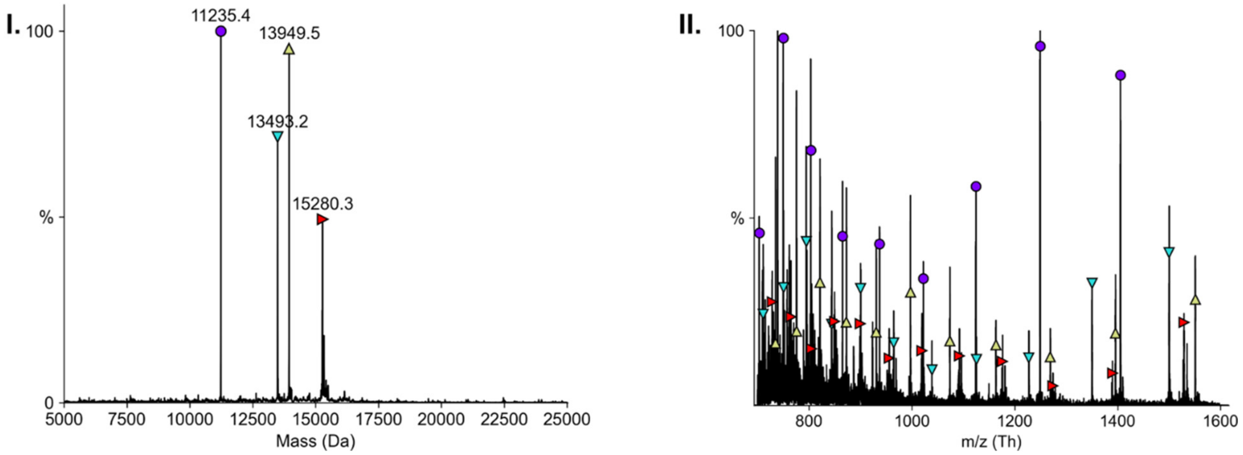

**D. 147 bp H3K4-trimethyl nucleosome.** (**I**) Deconvoluted mass spectrum: H4 (purple circle) calculated mass 11236.15 Da, found: 11235.4 Da; H2B (teal downward pointed triangle) calculated mass 13493.68 Da, found: 13493.0 Da; H2A (green upward pointed triangle) calculated mass 13950.2 Da, found: 13949.5 Da; H3K4me3 (red rightward pointed triangle) calculated mass 15308.70 Da, found: 15308.2 Da. (**II**) Raw mass spectrum: H4 (purple circle); H2B (teal downward pointed triangle); H2A (green upward pointed triangle; H3K4me3 (red rightward pointed triangle).

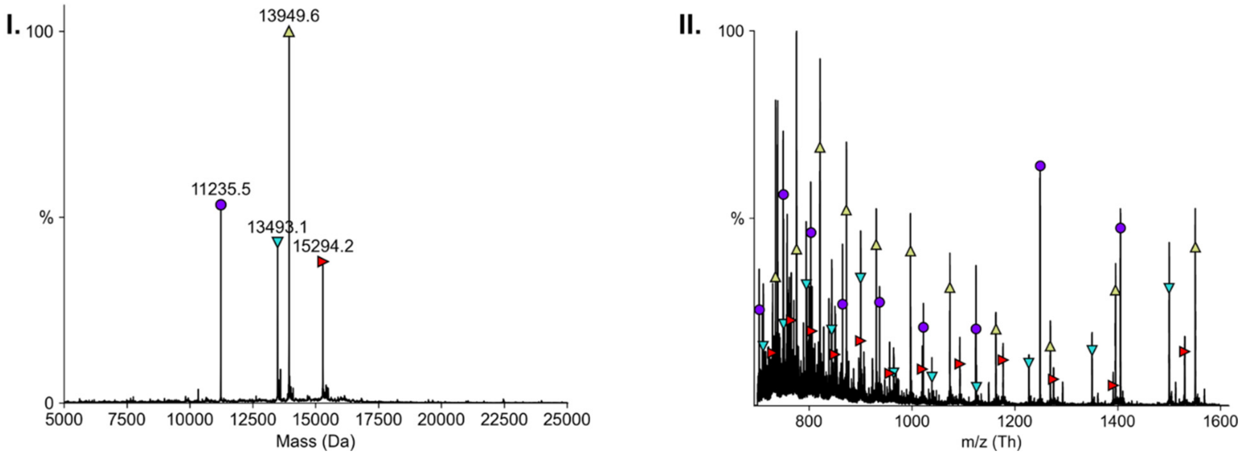

**E. 147 bp H3K9-propionyl nucleosome.** (**I**) Deconvoluted mass spectrum: H4 (purple circle) calculated mass 11236.15 Da, found: 11235.5 Da; H2B (teal downward pointed triangle) calculated mass 13493.68 Da, found: 13493.1 Da; H2A (green upward pointed triangle) calculated mass 13950.2 Da, found: 13949.6 Da; H3K9pr (red rightward pointed triangle) calculated mass 15294.67 Da, found: 15294.2 Da. (**II**) Raw mass spectrum: H4 (purple circle); H2B (teal downward pointed triangle); H2A (green upward pointed triangle; H3K9pr (red rightward pointed triangle).

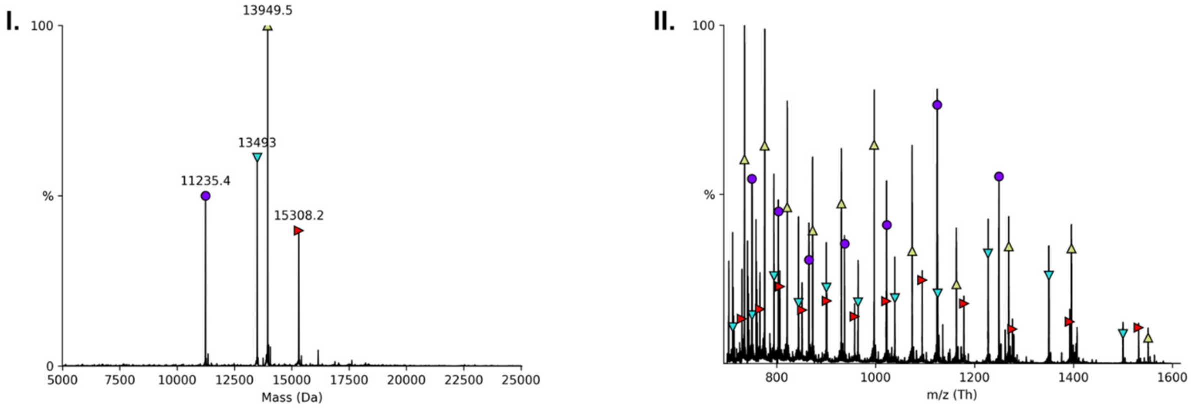

**F. 147 bp H3K9-butyryl nucleosome.** (**I**) Deconvoluted mass spectrum: H4 (purple circle) calculated mass 11236.15 Da, found: 11235.4 Da; H2B (teal downward pointed triangle) calculated mass 13493.68 Da, found: 13493.0 Da; H2A (green upward pointed triangle) calculated mass 13950.2 Da, found: 13949.5 Da; H3K9bu (red rightward pointed triangle) calculated mass 15308.70 Da, found: 15308.2 Da. (**II**) Raw mass spectrum: H4 (purple circle); H2B (teal downward pointed triangle); H2A (green upward pointed triangle; H3K9bu (red rightward pointed triangle).

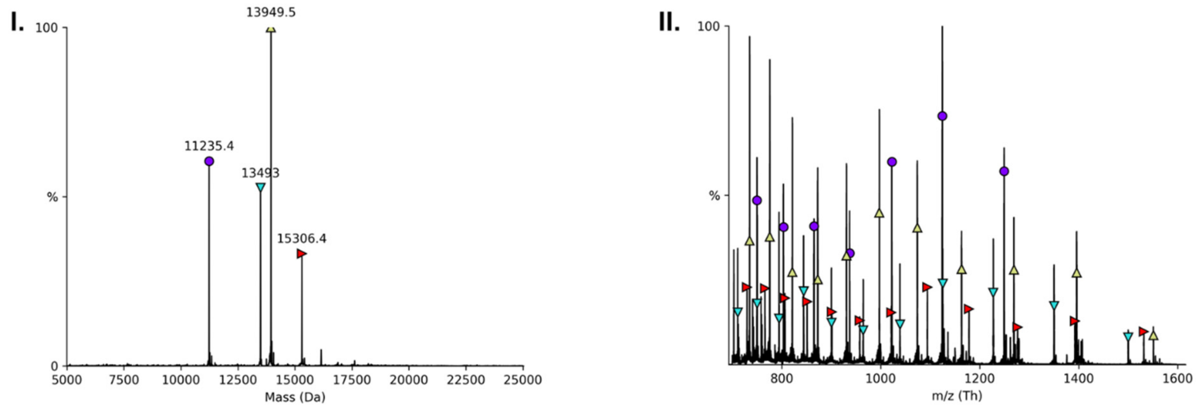

**G. 147 bp H3K9-crotonyl nucleosome.** (**I**) Deconvoluted mass spectrum: H4 (purple circle) calculated mass 11236.15 Da, found: 11235.4 Da; H2B (teal downward pointed triangle) calculated mass 13493.68 Da, found: 13493.0 Da; H2A (green upward pointed triangle) calculated mass 13950.2 Da, found: 13949.5 Da; H3K9cro (red rightward pointed triangle) calculated mass 15306.68, found: 15306.4. (**II**) Raw mass spectrum: H4 (purple circle); H2B (teal downward pointed triangle); H2A (green upward pointed triangle); H3K9cro (red rightward pointed triangle).

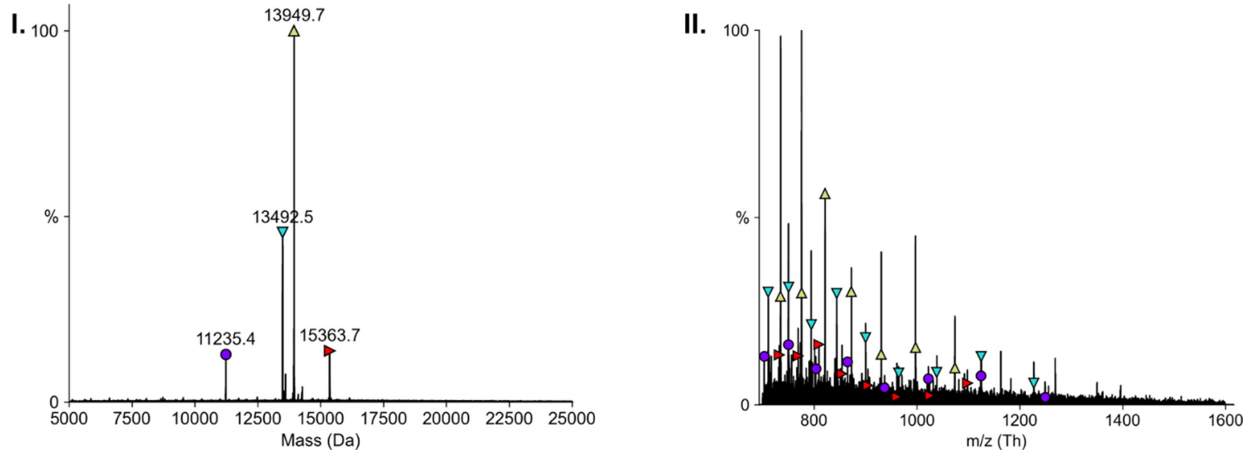

**H. 147 bp H3K9-octanoyl nucleosome.** (**I**) Deconvoluted mass spectrum: H4 (purple circle) calculated mass 11236.15 Da, found: 11235.2 Da; H2B (teal downward pointed triangle) calculated mass 13493.68 Da, found: 13493.3 Da; H2A (green upward pointed triangle) calculated mass 13950.2 Da, found: 13948.7 Da; H3K9oct (red rightward pointed triangle) calculated mass 15363.80, found: 15364.8. (**II**) Raw mass spectrum: H4 (purple circle); H2B (teal downward pointed triangle); H2A (green upward pointed triangle); H3K9oct (red rightward pointed triangle).

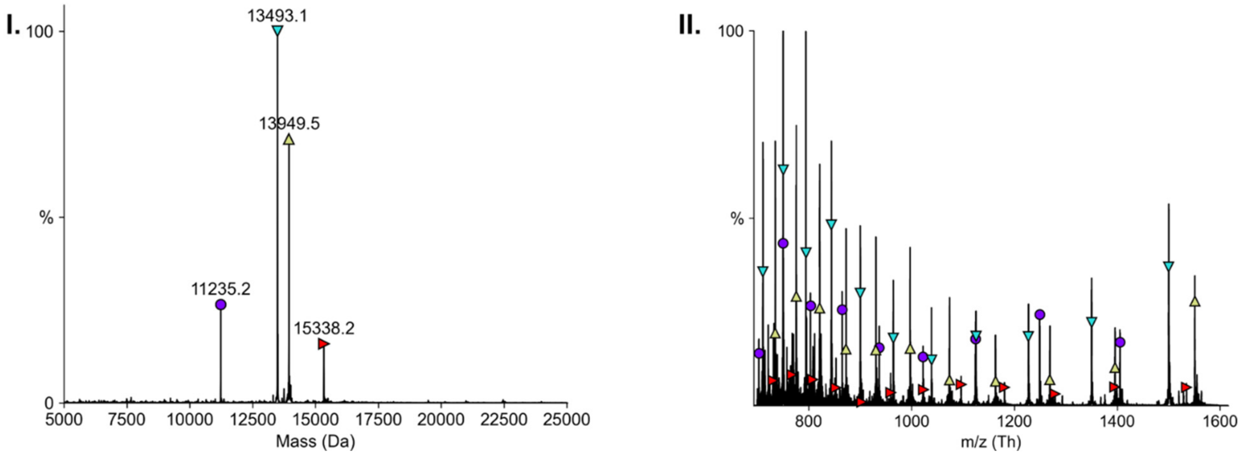

**I. 147 bp H3K9-succinyl nucleosome.** (**I**) Deconvoluted mass spectrum: H4 (purple circle) calculated mass 11236.15 Da, found: 11234.9 Da; H2B (teal downward pointed triangle) calculated mass 13493.68 Da, found: 13942.6 Da; H2A (green upward pointed triangle) calculated mass 13950.2 Da, found: 13949.7 Da; H3K9suc (red rightward pointed triangle) calculated mass 15337.67, found: 15338.6. (**II**) Raw mass spectrum: H4 (purple circle); H2B (teal downward pointed triangle); H2A (green upward pointed triangle); H3K9suc (red rightward pointed triangle).

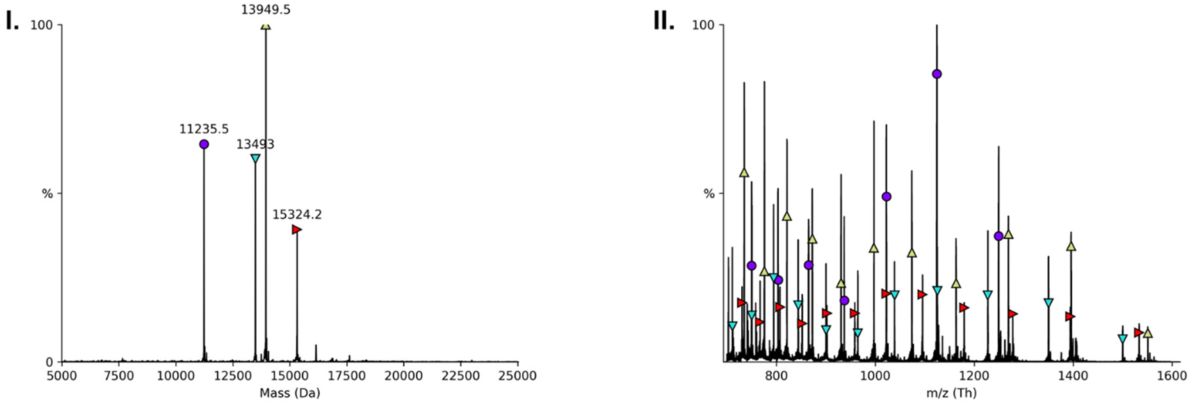

**J. 147 bp H3K9-anti-hydroxyisobutyryl nucleosome.** (**I**) Deconvoluted mass spectrum: H4 (purple circle) calculated mass 11236.15 Da, found: 11235.5 Da; H2B (teal downward pointed triangle) calculated mass 13493.68 Da, found: 13493.0 Da; H2A (green upward pointed triangle) calculated mass 13950.2 Da, found: 13949.5 Da; H3K9hib (red rightward pointed triangle) calculated mass 15324.70, found: 15324.2. (**II**) Raw mass spectrum: H4 (purple circle); H2B (teal downward pointed triangle); H2A (green upward pointed triangle); H3K9hib (red rightward pointed triangle).

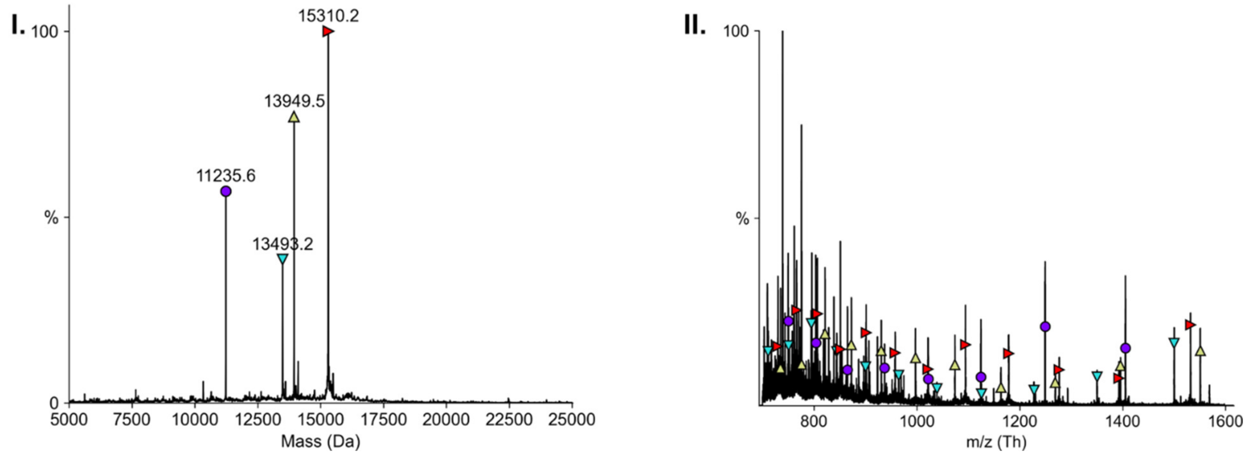

**K. 147 bp H3K9-lactyl nucleosome.** (**I**) Deconvoluted mass spectrum: H4 (purple circle) calculated mass 11236.15 Da, found: 11235.5 Da; H2B (teal downward pointed triangle) calculated mass 13493.68 Da, found: 13493.0 Da; H2A (green upward pointed triangle) calculated mass 13950.2 Da, found: 13949.5 Da; H3K9lac (red rightward pointed triangle) calculated mass 15310.67, found: 15310.2. (**II**) Raw mass spectrum: H4 (purple circle); H2B (teal downward pointed triangle); H2A (green upward pointed triangle); H3K9lac (red rightward pointed triangle).

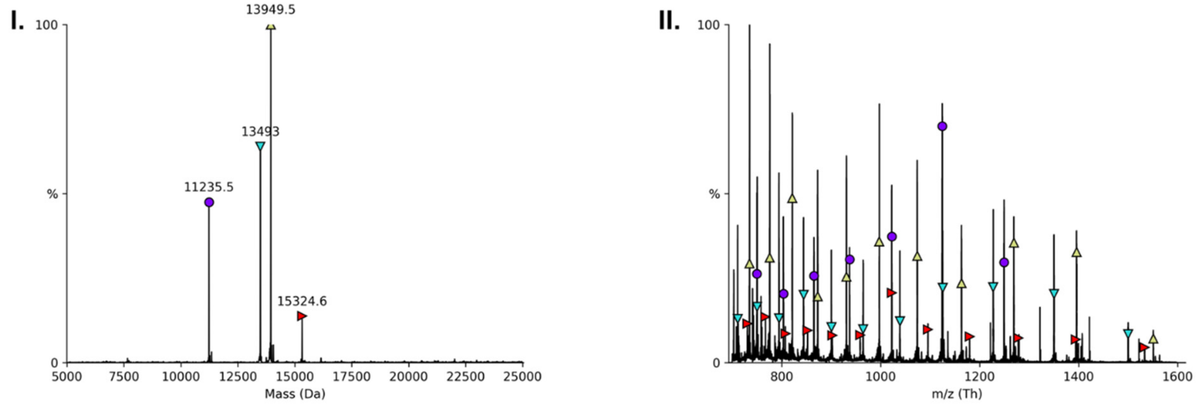

**L. 147 bp H3K9-β-hydroxybutyryl nucleosome.** (**I**) Deconvoluted mass spectrum: H4 (purple circle) calculated mass 11236.15 Da, found: 11235.5 Da; H2B (teal downward pointed triangle) calculated mass 13493.68 Da, found: 13493.0 Da; H2A (green upward pointed triangle) calculated mass 13950.2 Da, found: 13949.5 Da; H3K9bhb (red rightward pointed triangle) calculated mass 15324.70, found: 15324.6. (**II**) Raw mass spectrum: H4 (purple circle); H2B (teal downward pointed triangle); H2A (green upward pointed triangle); H3K9bhb (red rightward pointed triangle).

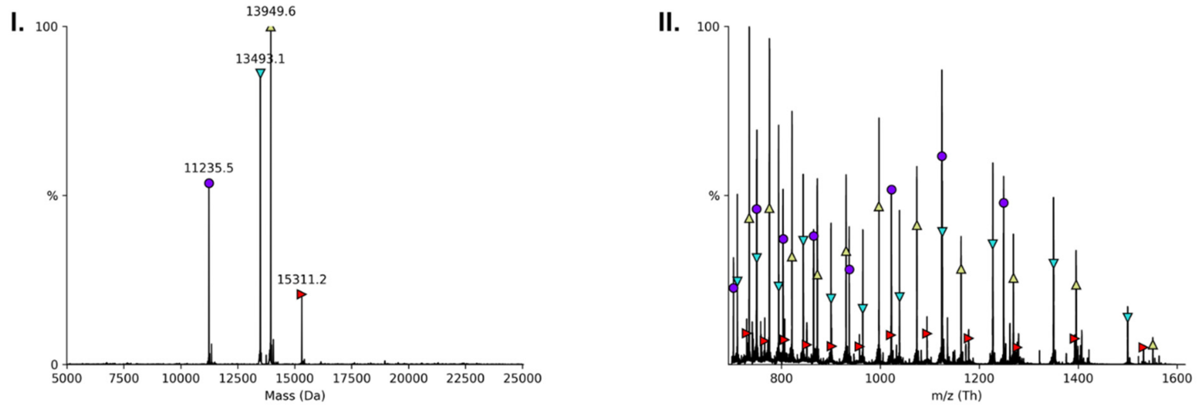

**M. 147 bp H3K9-N-methyl thiourea nucleosome.** (**I**) Deconvoluted mass spectrum: H4 (purple circle) calculated mass 11236.15 Da, found: 11235.5 Da; H2B (teal downward pointed triangle) calculated mass 13493.68 Da, found: 13493.1 Da; H2A (green upward pointed triangle) calculated mass 13950.2 Da, found: 13949.6 Da; H3K9mtu (red rightward pointed triangle) calculated mass 15311.72, found: 15311.2 Da. (**II**) Raw mass spectrum: H4 (purple circle); H2B (teal downward pointed triangle); H2A (green upward pointed triangle); H3K9mtu (red rightward pointed triangle).

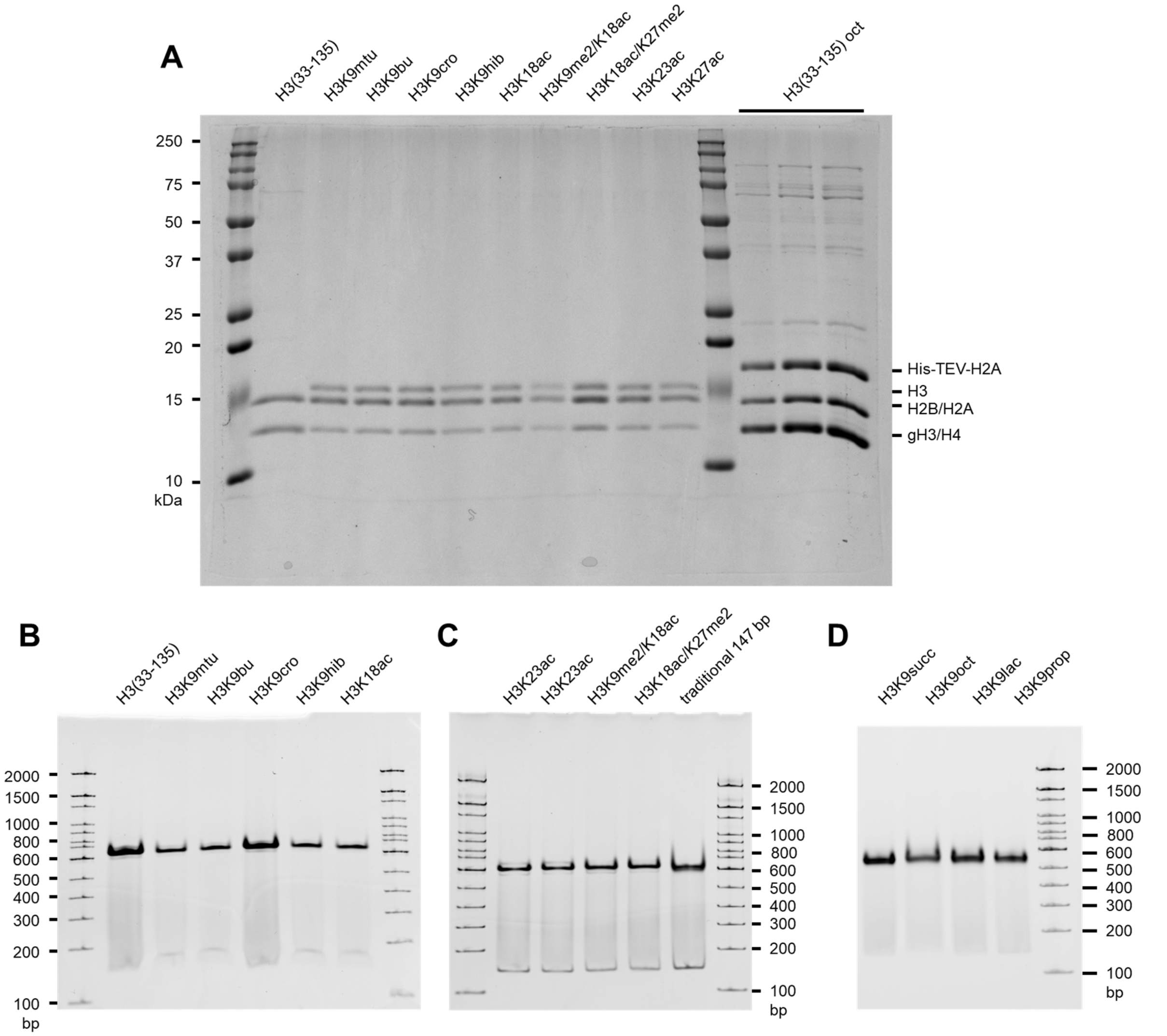

**Figure S10. SDS-PAGE and TBE native gel characterization of cW11 nucleosome ligation products.** (**A**) Representative SDS-PAGE of 147 bp H3(aa33-135) nucleosome starting material, products of the cW11 sortase ligation, and purified H3(aa33-135) octamer prior to TEV cleavage. (**B**) Representative TBE native gel of 147 bp H3(aa33-135) nucleosome starting material, products of the cW11 sortase ligation. (**C**) Representative TBE native gel of 147 bp products of the cW11 sortase ligation, and an unmodified 147 bp nucleosome prepared by traditional nucleosome reconstitution. (**D**) Representative TBE native gel of 147 bp products of the cW11 sortase ligation.

**Figure S11. Mass spectrometric characterization of peptide substrates used in cW11 nucleosome ligation.** Raw spectra deconvoluted with UniDec.[5]

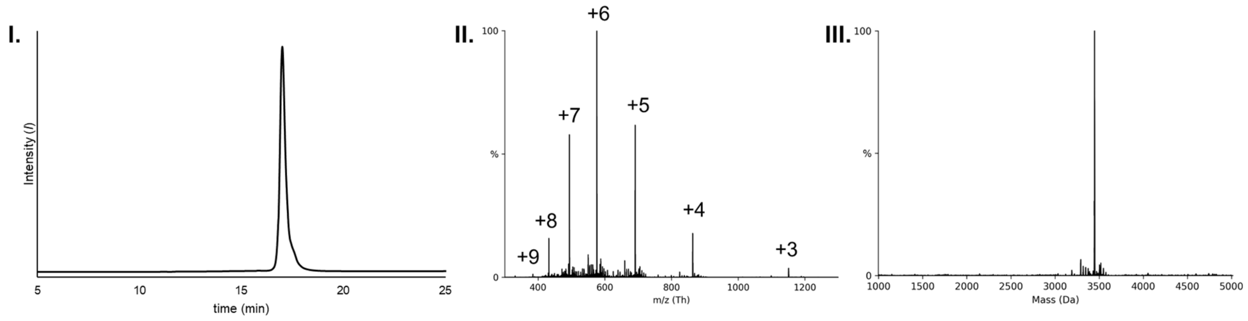

**A. H3(1-34) unmodified T32-G33 amide linkage. I.** Analytical RP-HPLC chromatogram (C18, 0-30% B, 23 min). **II.** Intact peptide ESI-MS. **III.** Deconvoluted peptide ESI-MS (Calculated exact mass for $C_{144}H_{261}N_{55}O_{43}$ $[M]^+$: 3448.99 Da; Observed: 3448.8 Da).

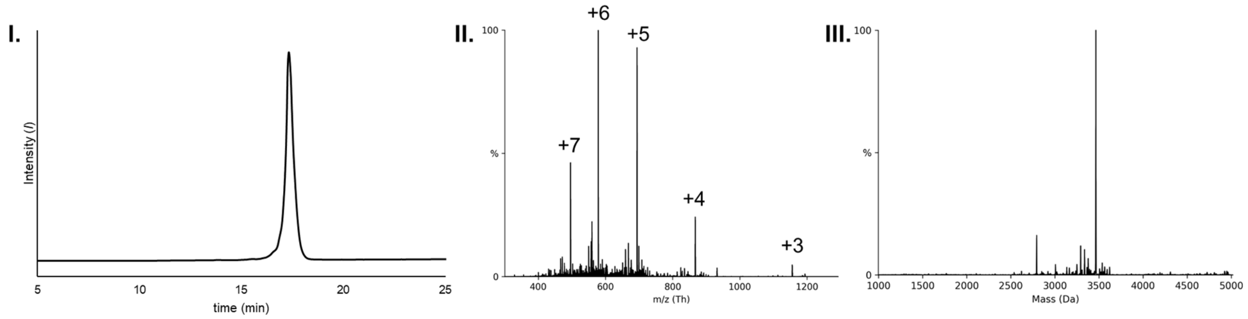

**B. H3(1-34) K4me1 T32-G33 amide linkage. I.** Analytical RP-HPLC chromatogram (C18, 0-30% B, 23 min). **II.** Intact peptide ESI-MS. **III.** Deconvoluted peptide ESI-MS (Calculated exact mass for $C_{145}H_{263}N_{55}O_{43}$ $[M]^+$: 3463.01 Da; Observed: 3463.8 Da).

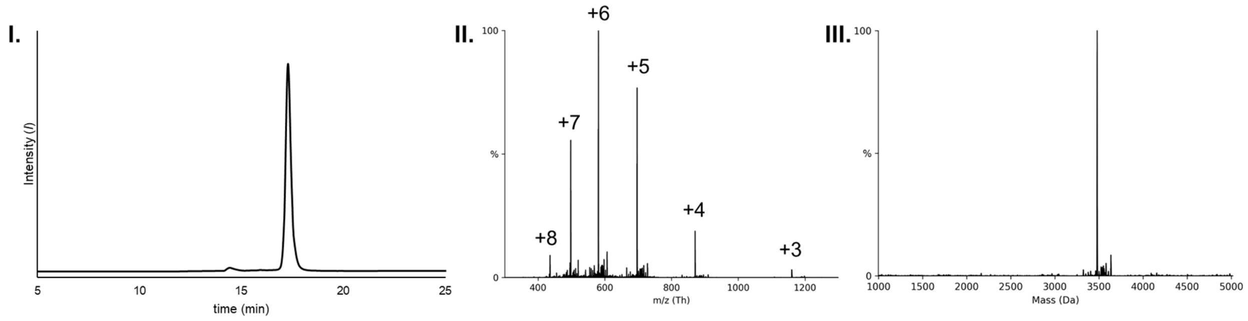

**C. H3(1-34) K4me2 T32-G33 amide linkage. I.** Analytical RP-HPLC chromatogram (C18, 0-30% B, 23 min). **II.** Intact peptide ESI-MS. **III.** Deconvoluted peptide ESI-MS (Calculated exact mass for $C_{146}H_{265}N_{55}O_{43}$ $[M]^+$: 3477.02 Da; Observed: 3476.9 Da).

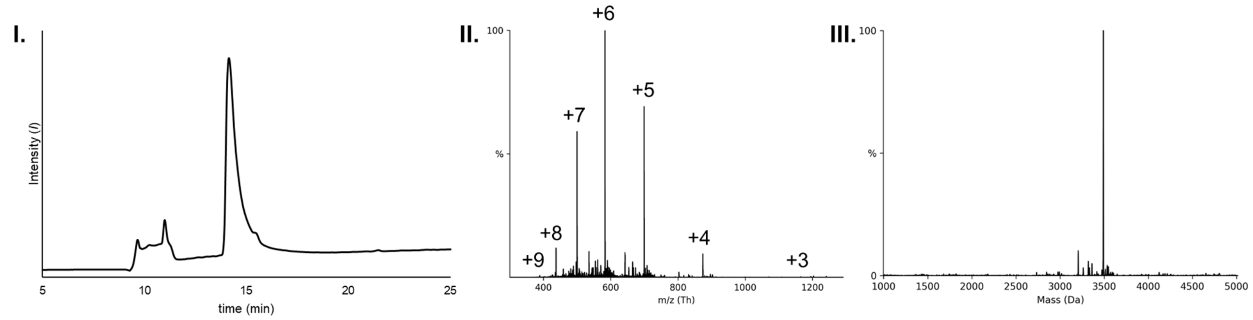

**D. H3(1-34) K4me3 T32-G33 amide linkage. I.** Analytical RP-HPLC chromatogram (C18, 7-30% B, 30 min). **II.** Intact peptide ESI-MS. **III.** Deconvoluted peptide ESI-MS (Calculated exact mass for $C_{148}H_{266}N_{54}O_{46}$ $[M]^{+}$: 3492.05 Da; Observed: 3491.1 Da).

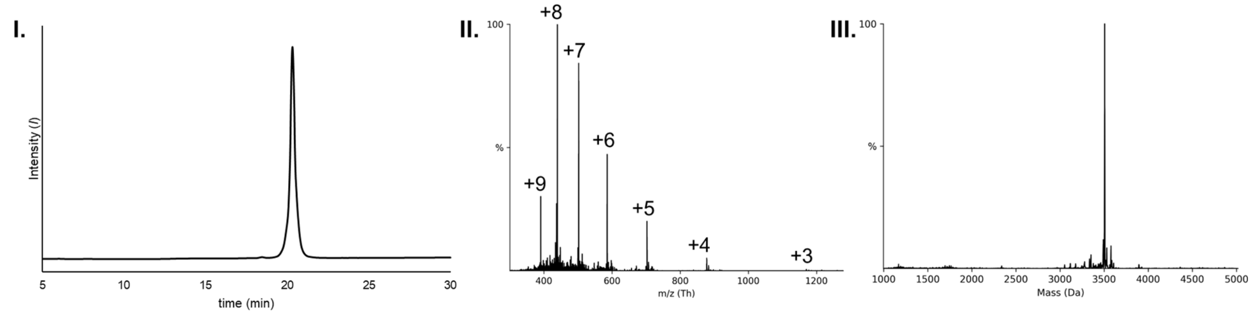

**E. H3(1-34) K9-propionyl T32-G33 depsipeptide. I.** Analytical RP-HPLC chromatogram (C18, 7-30% B, 30 min). **II.** Intact peptide ESI-MS. **III.** Deconvoluted peptide ESI-MS (Calculated exact mass for $C_{148}H_{266}N_{54}O_{45}$ $[M]^{+}$: 3520.02 Da; Observed: 3519.7 Da).

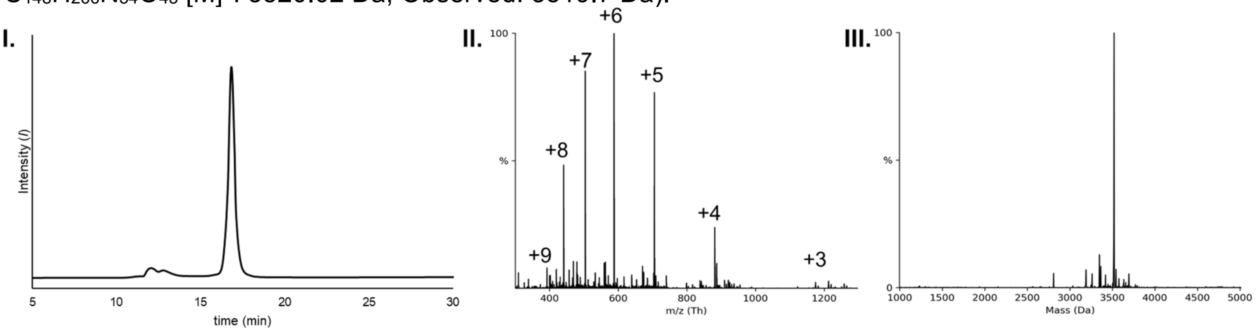

**F. H3(1-34) K9-butyryl T32-G33 depsipeptide. I.** Analytical RP-HPLC chromatogram (C18, 7-30% B, 30 min). **II.** Intact peptide ESI-MS. **III.** Deconvoluted peptide ESI-MS (Calculated exact mass for $C_{148}H_{266}N_{54}O_{45}$ $[M]^{+}$: 3520.02 Da; Observed: 3519.7 Da).

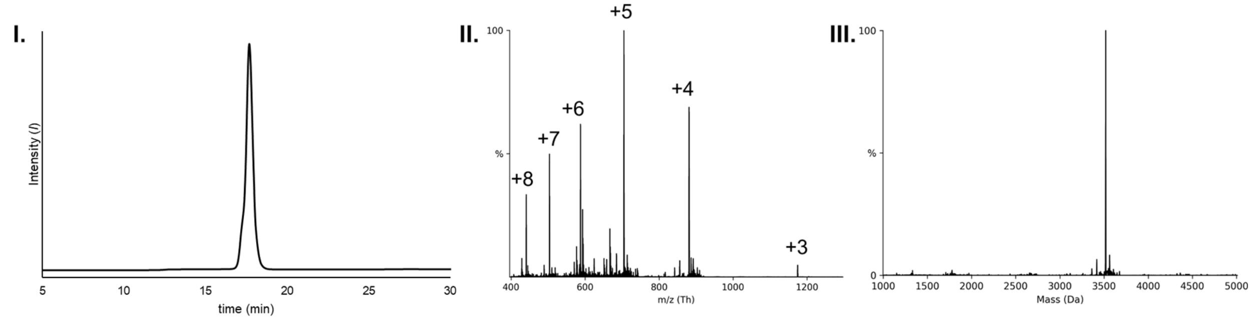

**G. H3(1-34) K9-crotonyl T32-G33 depsipeptide. I.** Analytical RP-HPLC chromatogram (C18, 7-30% B, 30 min). **II.** Intact peptide ESI-MS. **III.** Deconvoluted peptide ESI-MS (Calculated exact mass for $C_{148}H_{264}N_{54}O_{45}$ $[M]^{+}$: 3518.00 Da; Observed: 3518.7 Da).

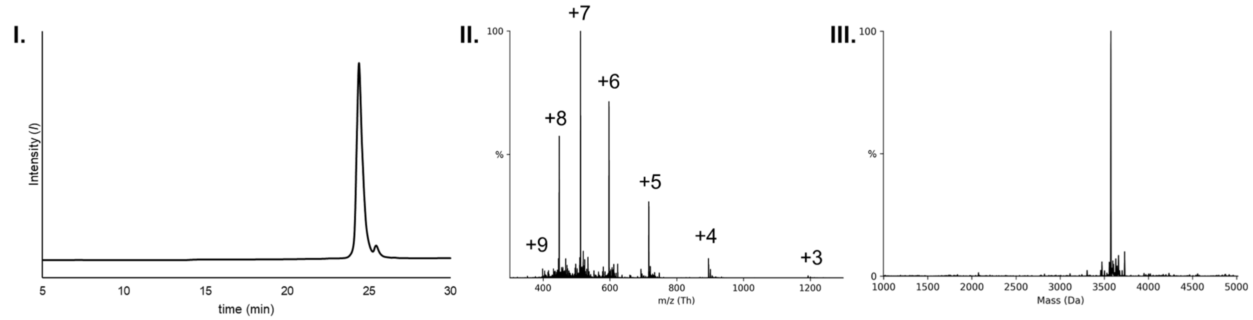

**H. H3(1-34) K9-octanoyl T32-G33 depsipeptide. I.** Analytical RP-HPLC chromatogram (C18, 0-30% B, 23 min). **II.** Intact peptide ESI-MS. **III.** Deconvoluted peptide ESI-MS (Calculated exact mass for $C_{152}H_{274}N_{54}O_{45}$ $[M]^{+}$: 3576.08 Da; Observed: 3575.7 Da).

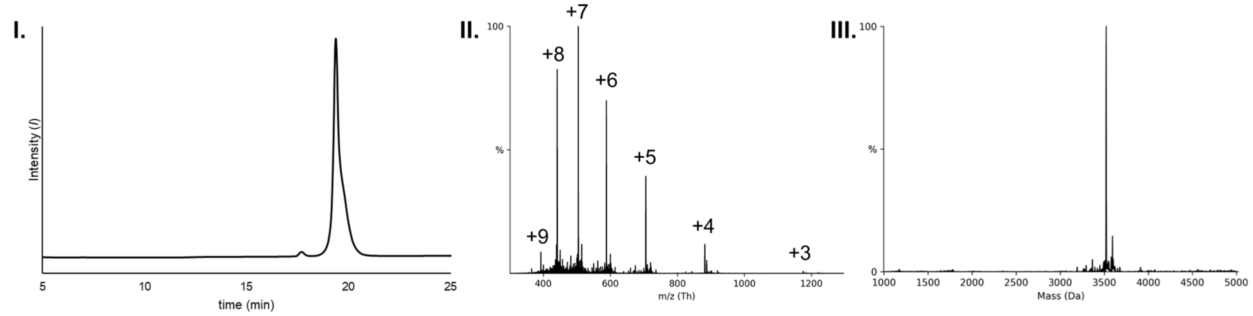

**I. H3(1-34) K9-lactyl T32-G33 depsipeptide. I.** Analytical RP-HPLC chromatogram (C18, 0-30% B, 23 min). **II.** Intact peptide ESI-MS. **III.** Deconvoluted peptide ESI-MS (Calculated exact mass for $C_{147}H_{264}N_{54}O_{46}$ $[M]^{+}$: 3522.00 Da; Observed: 3521.5 Da).

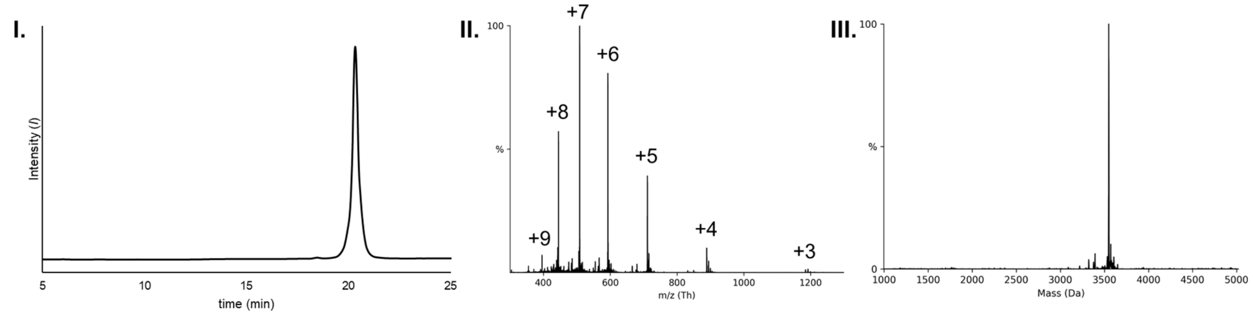

**J. H3(1-34) K9-succinyl T32-G33 depsipeptide. I.** Analytical RP-HPLC chromatogram (C18, 0-30% B, 23 min). **II.** Intact peptide ESI-MS. **III.** Deconvoluted peptide ESI-MS (Calculated exact mass for $C_{148}H_{264}N_{54}O_{47}$ $[M]^{+}$: 3549.99 Da; Observed: 3549.4 Da).

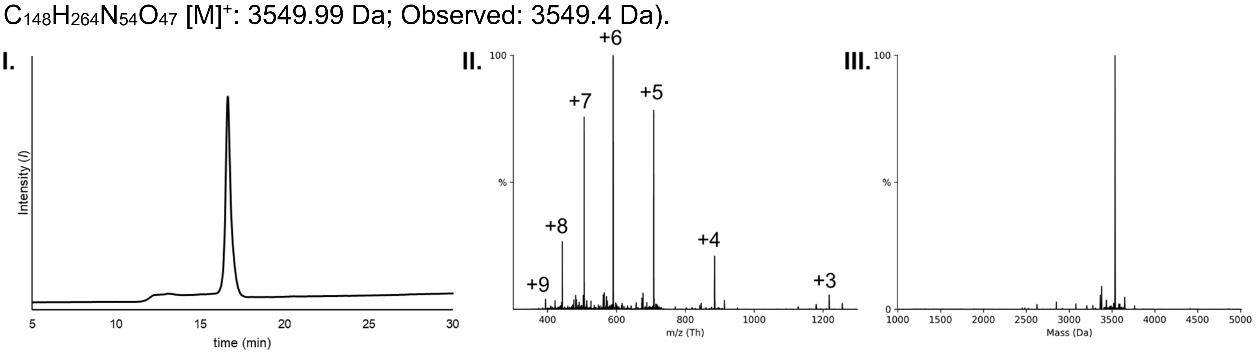

**K. H3(1-34) K9-anti-hydroxyisbutyryl T32-G33 depsipeptide. I.** Analytical RP-HPLC chromatogram (C18, 7-30% B, 30 min). **II.** Intact peptide ESI-MS. **III.** Deconvoluted peptide ESI-MS (Calculated exact mass for $C_{148}H_{266}N_{54}O_{46}$ $[M]^{+}$: 3536.01 Da; Observed: 3535.8 Da).

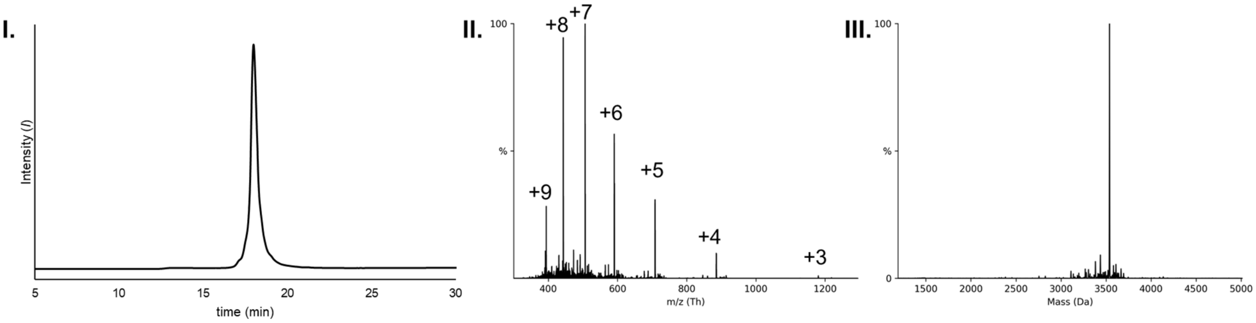

**L. H3(1-34) K9-β-hydroxybutyryl T32-G33 depsipeptide. I.** Analytical RP-HPLC chromatogram (C18, 7-30% B, 30 min). **II.** Intact peptide ESI-MS. **III.** Deconvoluted peptide ESI-MS (Calculated exact mass for $C_{148}H_{266}N_{54}O_{46}$ $[M]^{+}$: 3536.01 Da; Observed: 3536.3 Da).

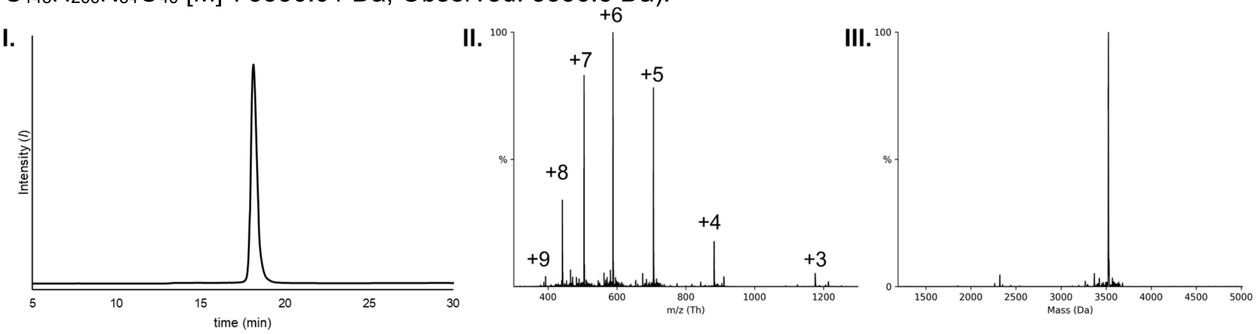

**M. H3(1-34) K9-N-methylthiourea T32-G33 depsipeptide. I.** Analytical RP-HPLC chromatogram (C18, 7-30% B, 30 min). **II.** Intact peptide ESI-MS. **III.** Deconvoluted peptide ESI-MS (Calculated exact mass for $C_{146}H_{263}N_{55}O_{44}S$ $[M]^{+}$: 3522.98 Da; Observed: 3523.7 Da).

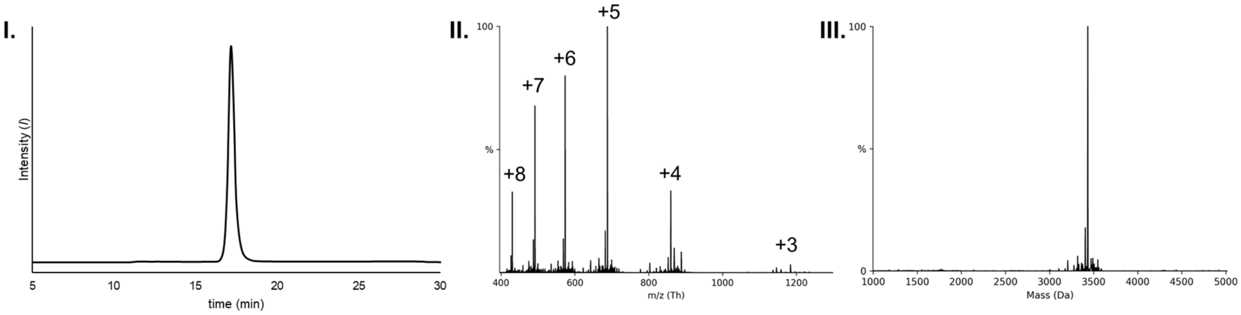

**N. H3(1-34) K9norleucine T32-G33 depsipeptide. I.** Analytical RP-HPLC chromatogram (C18, 7-30% B, 30 min). **II.** Intact peptide ESI-MS. **III.** Deconvoluted peptide ESI-MS (Calculated exact mass for $C_{144}H_{259}N_{53}O_{44}$ $[M]^{+}$: 3433.98 Da; Observed: 3433.9 Da).

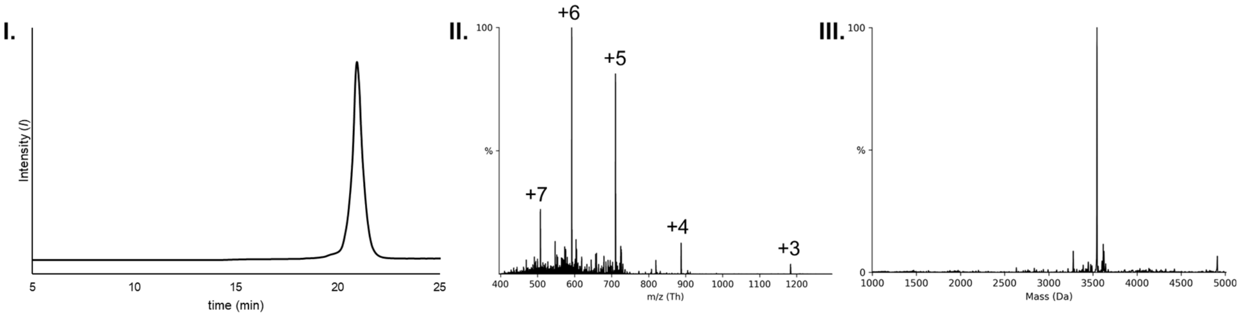

**O. H3(1-34) K9-myristoyl T32-G33 depsipeptide. I.** Analytical RP-HPLC chromatogram (C18, 7-30% B, 30 min). **II.** Intact peptide ESI-MS. **III.** Deconvoluted peptide ESI-MS (Calculated exact mass for $C_{158}H_{286}N_{54}O_{45}$ $[M]^{+}$: 3660.18 Da; Observed: 3659.8 Da).

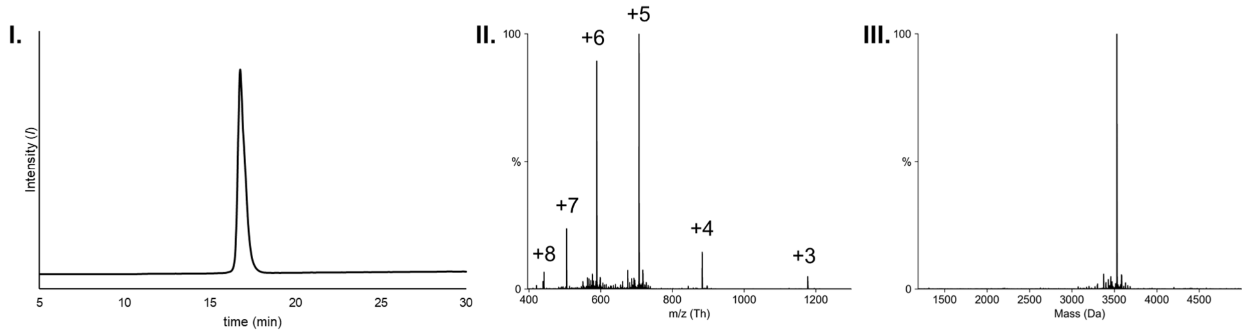

**P. H3(1-34) S28-phosphoryl T32-G33 depsipeptide. I.** Analytical RP-HPLC chromatogram (C18, 0-65% B, 30 min). **II.** Intact peptide ESI-MS. **III.** Deconvoluted peptide ESI-MS (Calculated exact mass for $C_{144}H_{260}N_{54}O_{47}P$ $[M]^{+}$: 3528.94 Da; Observed: 3529.0 Da).

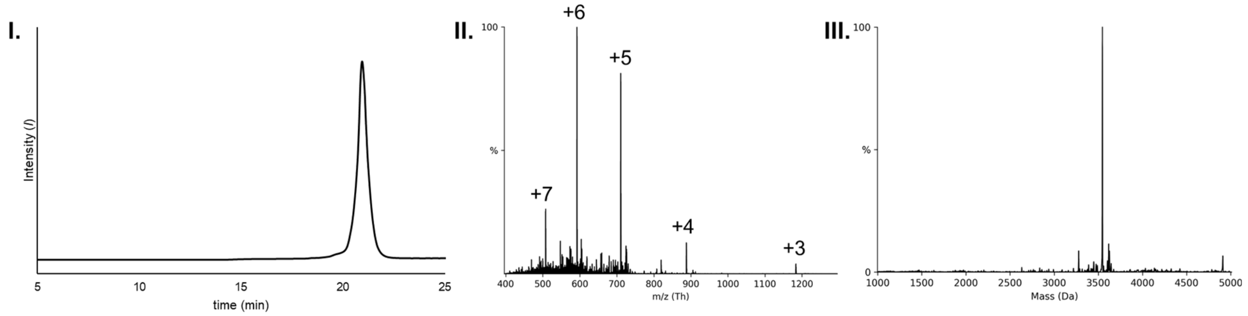

**Q. H3(1-34) S31-phosphoryl T32-G33 depsipeptide. I.** Analytical RP-HPLC chromatogram (C18, 7-30% B, 30 min). **II.** Intact peptide ESI-MS. **III.** Deconvoluted peptide ESI-MS (Calculated exact mass for $C_{144}H_{260}N_{54}O_{48}P$ $[M]^{+}$: 3544.93 Da; Observed: 3546.1 Da).

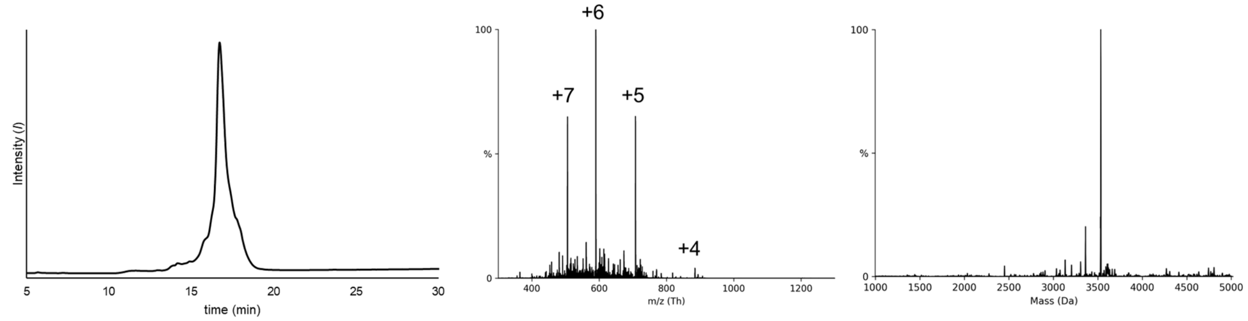

**R. H3(1-34) K4-trimethyl K9-acetyl T32-G33 depsipeptide. I.** Analytical RP-HPLC chromatogram (C18, 7-30% B, 30 min). **II.** Intact peptide ESI-MS. **III.** Deconvoluted peptide ESI-MS (Calculated exact mass for $C_{144}H_{260}N_{54}O_{48}P$ $[M]^{+}$: 3544.93 Da; Observed: 3546.1 Da).

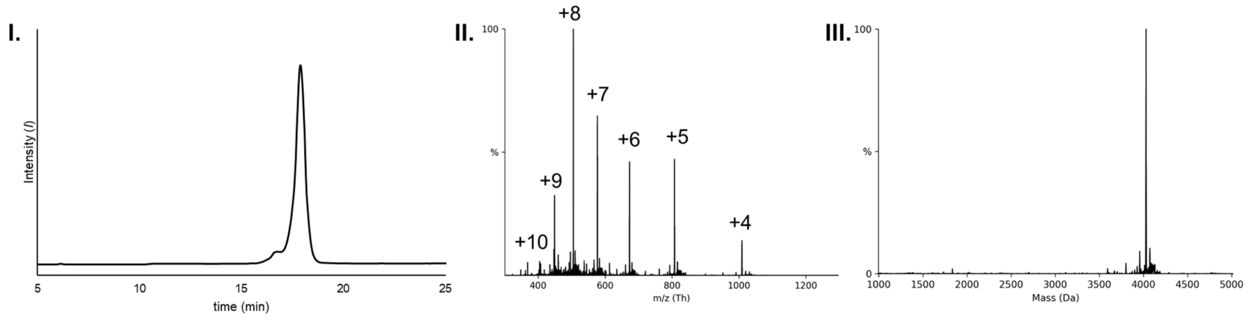

**S. H3(1-34) K9-acetyl K14-acetyl K18-acetyl K27-acetyl T32-G33 depsipeptide (aa35-37) KRK. I.** Analytical RP-HPLC chromatogram (C18, 7-30% B, 30 min). **II.** Intact peptide ESI-MS. **III.** Deconvoluted peptide ESI-MS (Calculated exact mass for $C_{170}H_{305}N_{62}O_{51}$ $[M]^{+}$: 4030.31 Da; Observed: 4029.1 Da).

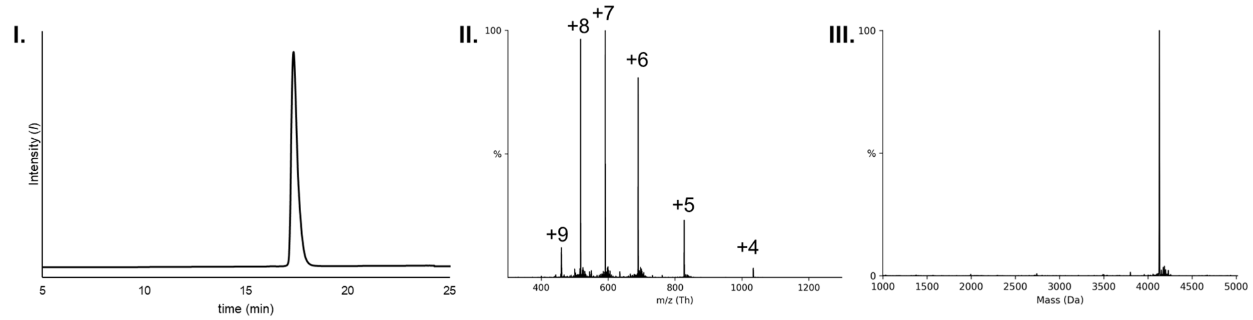

**T. H3(1-34) K9-acetyl K14-acetyl K18-acetyl K23-acetyl T32-G33 depsipeptide (aa35-38) KKKK. I.** Analytical RP-HPLC chromatogram (C18, 7-30% B, 30 min). **II.** Intact peptide ESI-MS. **III.** Deconvoluted peptide ESI-MS (Calculated exact mass $C_{176}H_{316}N_{62}O_{52}$ $[M]^+$: 4130.40 Da; Observed: 4130.3 Da).

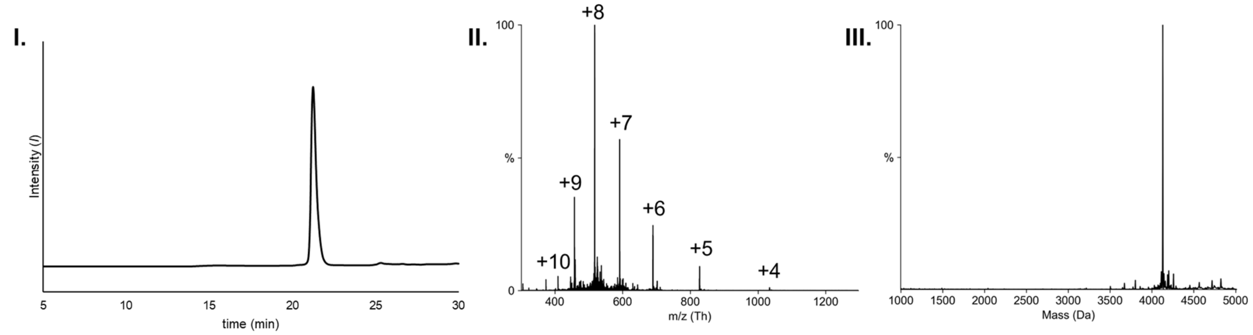

**U. H3(1-34) K9-acetyl K14-acetyl K23-acetyl K27-acetyl T32-G33 depsipeptide (aa35-38) KKKK. I.** Analytical RP-HPLC chromatogram (C18, 7-30% B, 30 min). **II.** Intact peptide ESI-MS. **III.** Deconvoluted peptide ESI-MS (Calculated exact mass $C_{176}H_{316}N_{62}O_{52}$ $[M]^+$: 4130.40 Da; Observed: 4130.0 Da).

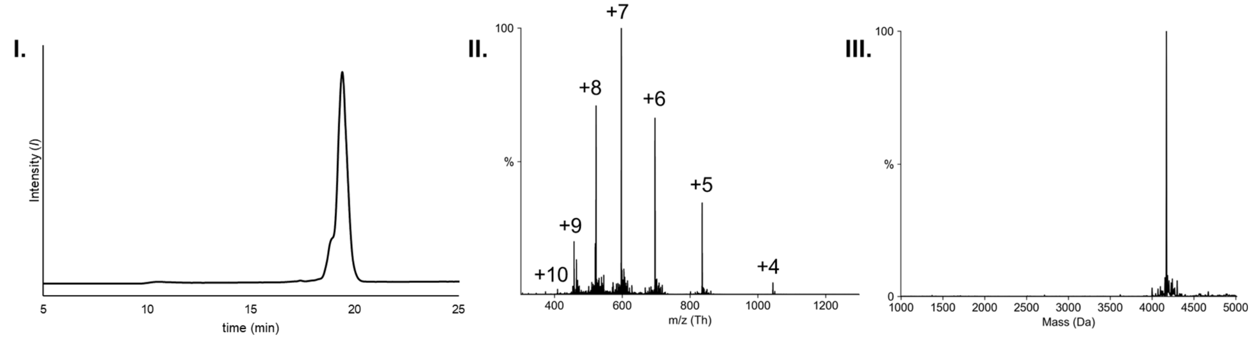

**V. H3(1-34) K9-acetyl K14-acetyl K18-acetyl K23-acetyl K27-acetyl T32-G33 depsipeptide (aa35-38) KKKK. I.** Analytical RP-HPLC chromatogram (C18, 7-30% B, 30 min). **II.** Intact peptide ESI-MS. **III.** Deconvoluted peptide ESI-MS (Calculated exact mass $C_{178}H_{318}N_{62}O_{53}$ $[M]^+$: 4172.41 Da; Observed: 4172.0 Da)

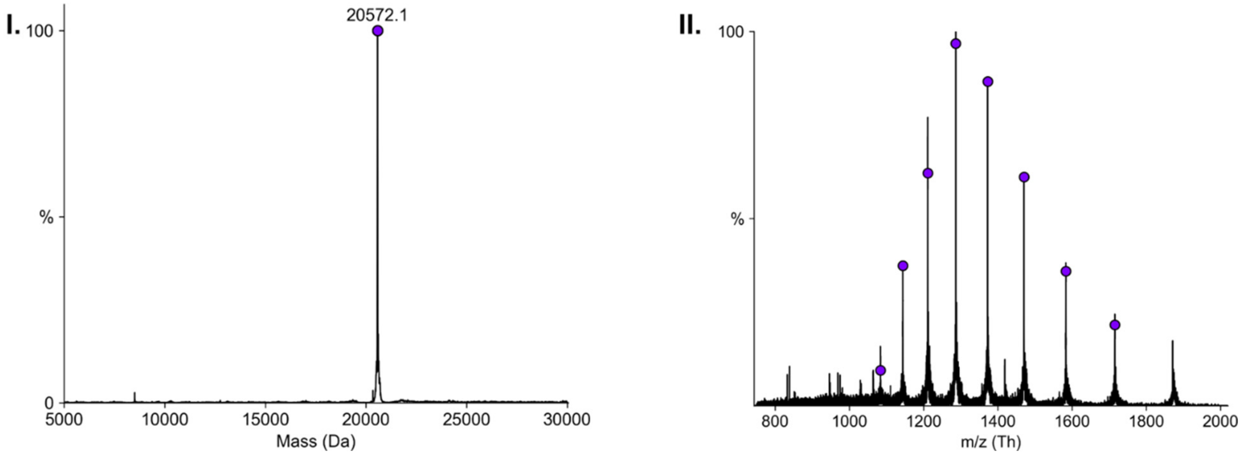

**W. H3(1-34) K18-Ubiquitin(G76A) K23-Ubiquitin(G76A) T32-G33 amide. I.** Deconvoluted peptide ESI-MS (Calculated exact mass $C_{902}H_{1519}N_{265}O_{277}S_2$ $[M]^+$: 20572.45 Da; Observed: 20572.1 Da). **II.** Intact peptide ESI-MS.

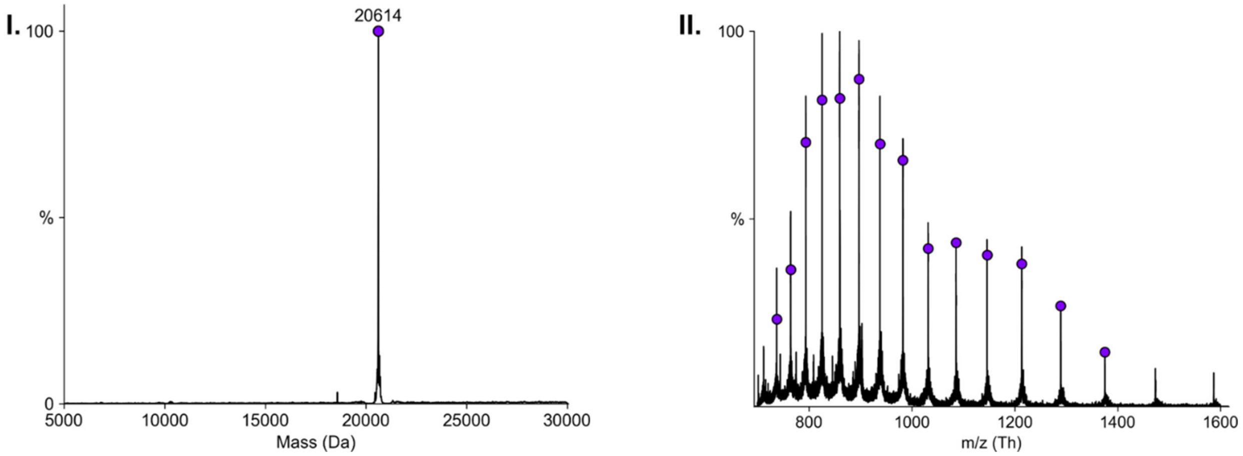

**X. H3(1-34) K9-trimethyl K18-Ubiquitin(G76A) K23-Ubiquitin(G76A) T32-G33 amide. I.** Deconvoluted peptide ESI-MS (Calculated exact mass for $C_{905}H_{1525}N_{265}O_{277}S_2$ $[M]^+$: 20614.53 Da; Observed: 20614.0 Da). **II.** Intact peptide ESI-MS.

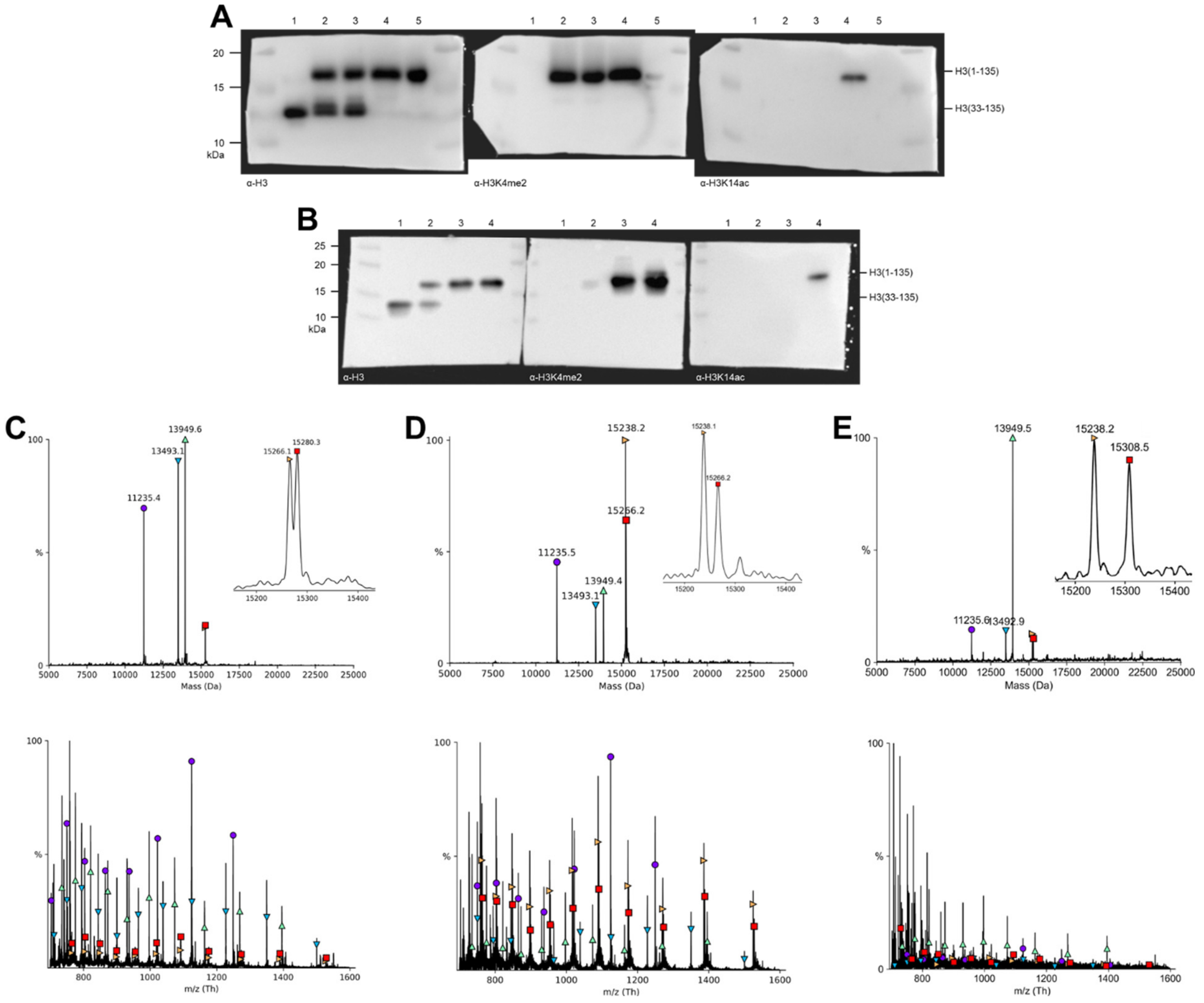

**Figure S12. Characterization of asymmetric H3K4me2 H3K14ac ligation intermediate and final products.** (**A**) Anti-H3 (left), anti-H3K4me2 (middle), and anti-H3K14ac (right) western blot visualization of asymmetric K4me2 nucleosome synthesis starting material, intermediates, and final products: H3 (aa33-135) starting material; (2) asymmetric H3K4me2 & H3 (aa33-135) intermediate; (3) asymmetric H3K4me2 & H3 (aa33-135) intermediate; (4) asymmetric H3K4me2 & H3K14ac product; (5) asymmetric H3K4me2 & unmodified H3 product. (**B**) Anti-H3 (left), anti-H3K4me2 (middle), and anti-H3K14ac (right) western blot visualization of asymmetric K4me2 nucleosome synthesis starting material, intermediates, and final products: H3 (aa33-135) starting material; (2) asymmetric H3K4me2 & H3 (aa33-135) intermediate; (3) asymmetric H3K4me2 & unmodified H3 product (4) asymmetric H3K4me2 & H3K14ac product. (**C**) Deconvoluted mass spectrum (top) and raw mass spectrum (bottom) of 185 bp asymmetric H3K4me2 & H3K14ac product: H4 (purple circle) calculated mass 11236.15 Da, found: 11235.4 Da; H2B (teal downward pointed triangle) calculated mass 13493.68 Da, found: 13493.1 Da; H2A (green upward pointed triangle) calculated mass 13950.2 Da, found: 13949.6 Da; H3K4me2 (orange rightward pointed triangle) calculated mass 15267.62 Da, found: 15266.1 Da; H3K14ac (red square) calculated mass 15281.61 Da, found: 15280.3 Da. (**D**) Deconvoluted mass spectrum (top) and raw mass spectrum (bottom) of 185 bp asymmetric unmodified H3 & H3K4me2 product: H4 (purple circle) calculated mass 11236.15 Da, found: 11235.5 Da; H2B (teal downward pointed triangle) calculated mass 13493.68 Da, found: 13493.1 Da; H2A (green upward pointed triangle) calculated mass 13950.2 Da, found: 13949.4 Da; H3 (orange rightward pointed triangle) calculated mass 15238.61 Da, found: 15238.1 Da; H3K4me2 (red square) calculated mass 15266.66 Da, found: 15266.2 Da. (**E**) Deconvoluted mass spectrum (top) and raw mass spectrum (bottom) of 185 bp asymmetric unmodified H3 & H3K4me2K14ac product: H4 (purple circle) calculated mass 11236.15 Da, found: 11235.6 Da; H2B (teal downward pointed triangle) calculated mass 13493.68 Da, found: 13492.9 Da; H2A (green upward pointed triangle) calculated mass 13950.2 Da, found: 13949.5 Da; H3 (orange rightward pointed triangle) calculated mass 15238.61 Da, found: 15238.2 Da; H3K4me2K14ac (red square) calculated mass 15309.71 Da, found: 15308.5 Da. Raw spectra deconvoluted with UniDec.[5]

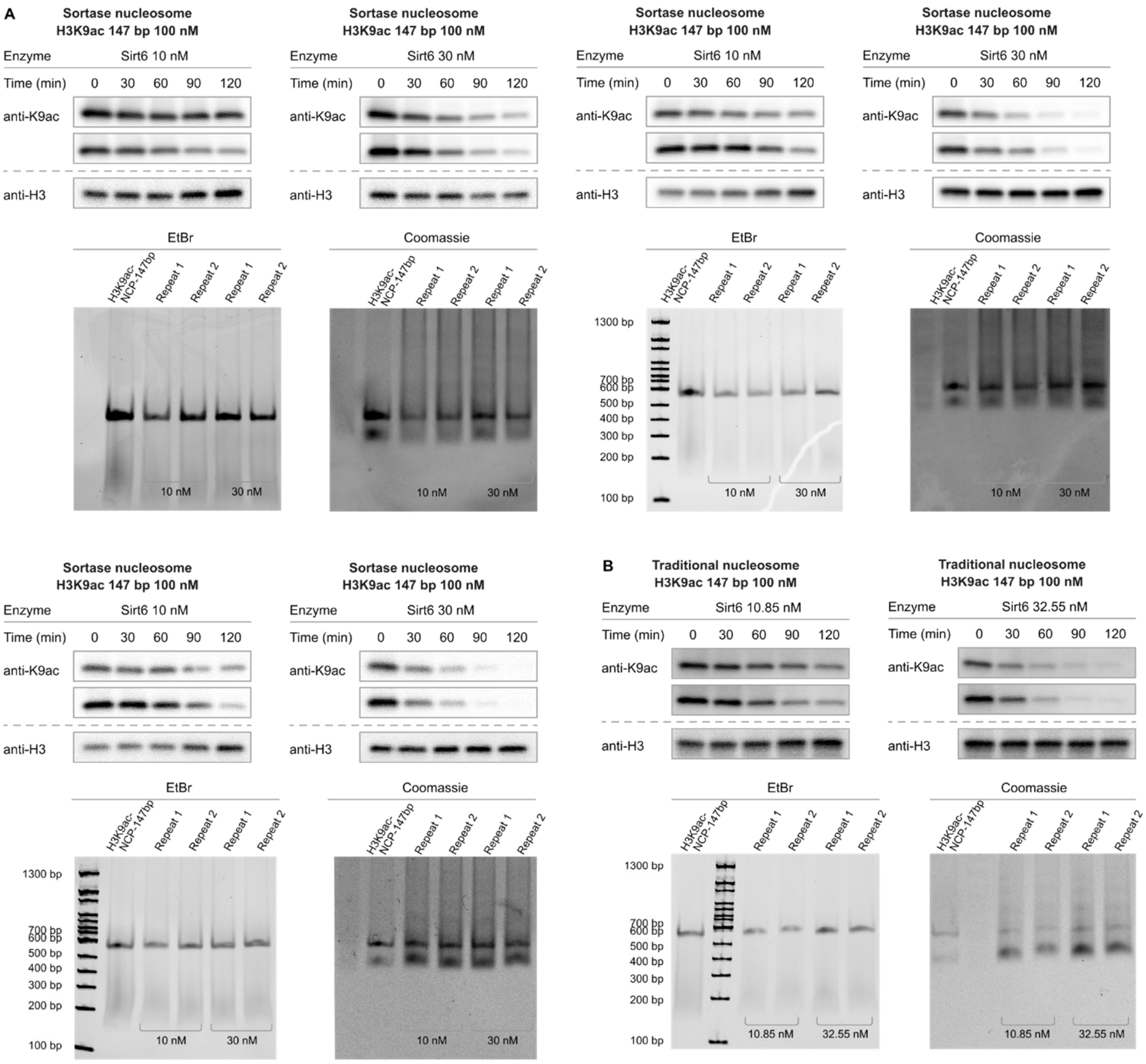

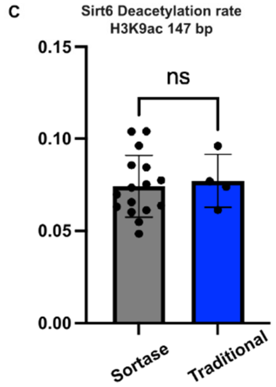

D

| Sirt6 Nucleosome | Sortase nucleosome | Traditional nucleosome |
|---|---|---|
| H3K9ac | 0.075±0.0044 | 0.076±0.0030 |

**Figure S13. Comparison of Sirt6 activity toward nucleosomes prepared by conventional reconstitution and cW11 nucleosome ligation.** (**A**) Representative western blots (top) and native TBE gels (bottom) of Sirt6 deacetylation assays with 147 bp H3K9ac nucleosome prepared by sortase ligation.

(**B**) Western blots (top) and native TBE gels (bottom) of Sirt6 deacetylation assays with 147 bp H3K9ac nucleosome prepared by traditional nucleosome reconstitution. (**C**) for H3K9ac nucleosomes prepared by the cW11 sortase ligation (Sortase) and nucleosomes prepared by literature protocols (Traditional). (**D**) Average V/[E] ± SEM values for Sirt6 deacetylation of H3K9ac nucleosomes prepared by either cW11 sortase ligation or literature protocols.

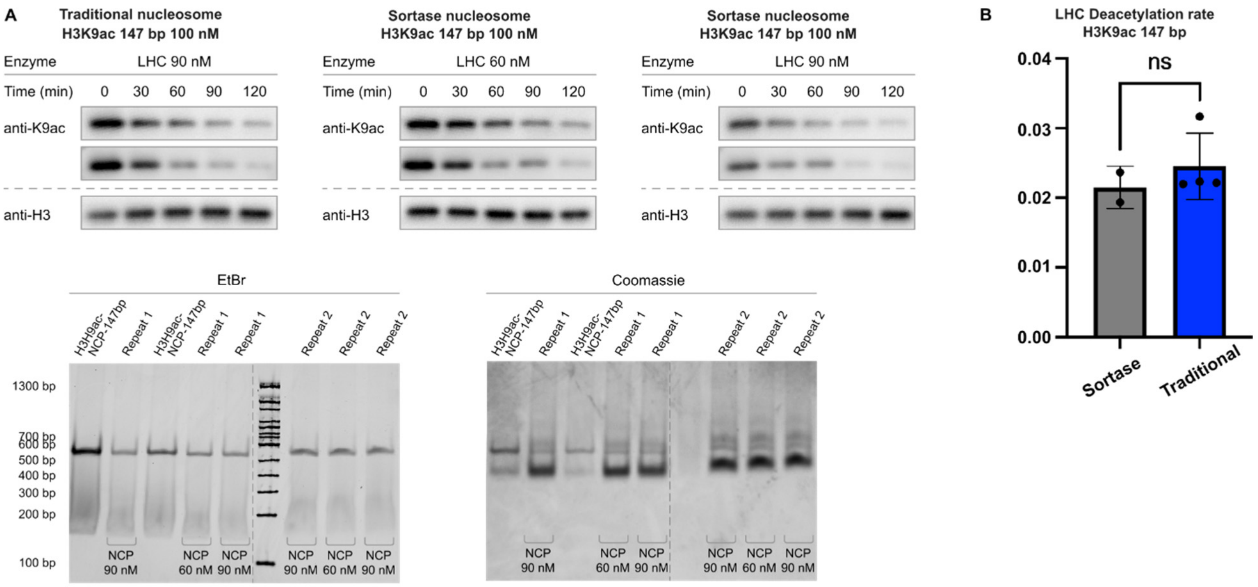

C

| LHC Nucleosome | Sortase nucleosome | Traditional nucleosome |
|---|---|---|
| H3K9ac | 0.021±0.0011 | 0.026±0.0021 |

**Figure S14. Comparison of LSD1/HDAC1/CoREST activity toward nucleosomes prepared by conventional reconstitution and cW11 nucleosome ligation.** (**A**) Western blots (top) and native TBE gels (bottom) of LSD1/HDAC1/CoREST (LHC) deacetylation assays with 147 bp H3K9ac nucleosome prepared by the sortase ligation (top right) or traditional nucleosome reconstitution (top left). (**B**) LHC complex deacetylation rates (V/[E]) for H3K9ac nucleosomes prepared by the cW11 sortase ligation (Sortase) and nucleosomes prepared by literature protocols (Traditional). (**C**) Average V/[E] ± SEM values for Sirt6 and LHC deacetylation of H3K9ac nucleosomes prepared by either cW11 sortase ligation or literature protocols.

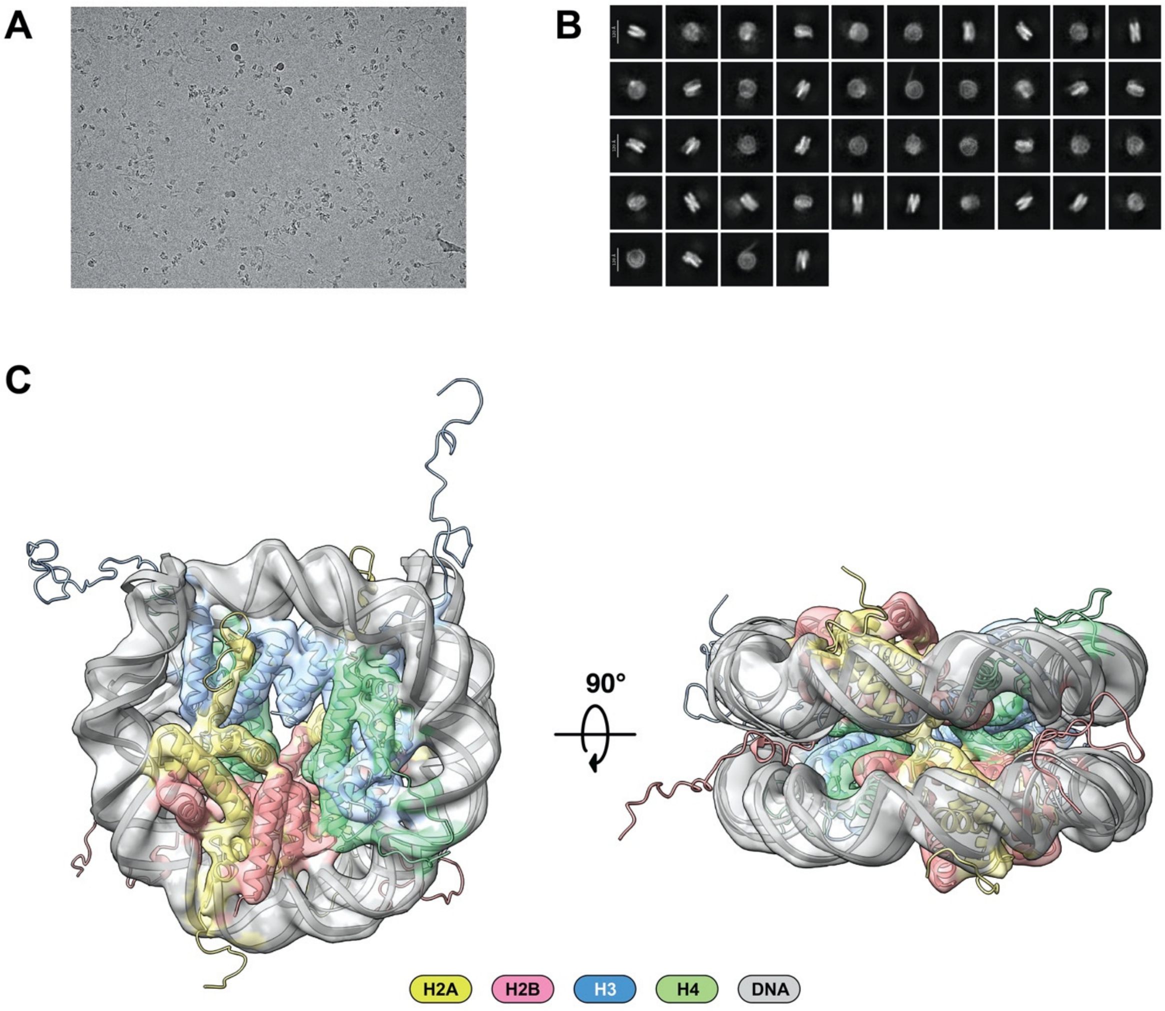

**Figure S15. Cryo electron microscopy characterization of nucleosomes prepared by cW11 nucleosome ligation.** (**A**) Raw cryo-EM images of cW11 nucleosome. (**B**) Representative 2D Class averages from 14300 particles. (**C**) 4.7-Å cryo-EM reconstruction map of cW11 nucleosome displayed in two separate views related by 90°. A structural model of canonical *X. laevis* nucleosome (PDBID: 1KX5) is docked in to the EM density. The EM reconstruction and the model were colored the following way: DNA in gray, histone H2A in pale yellow, histone H2B in red salmon, histone H3 in marine blue, and histone H4 in lime green.

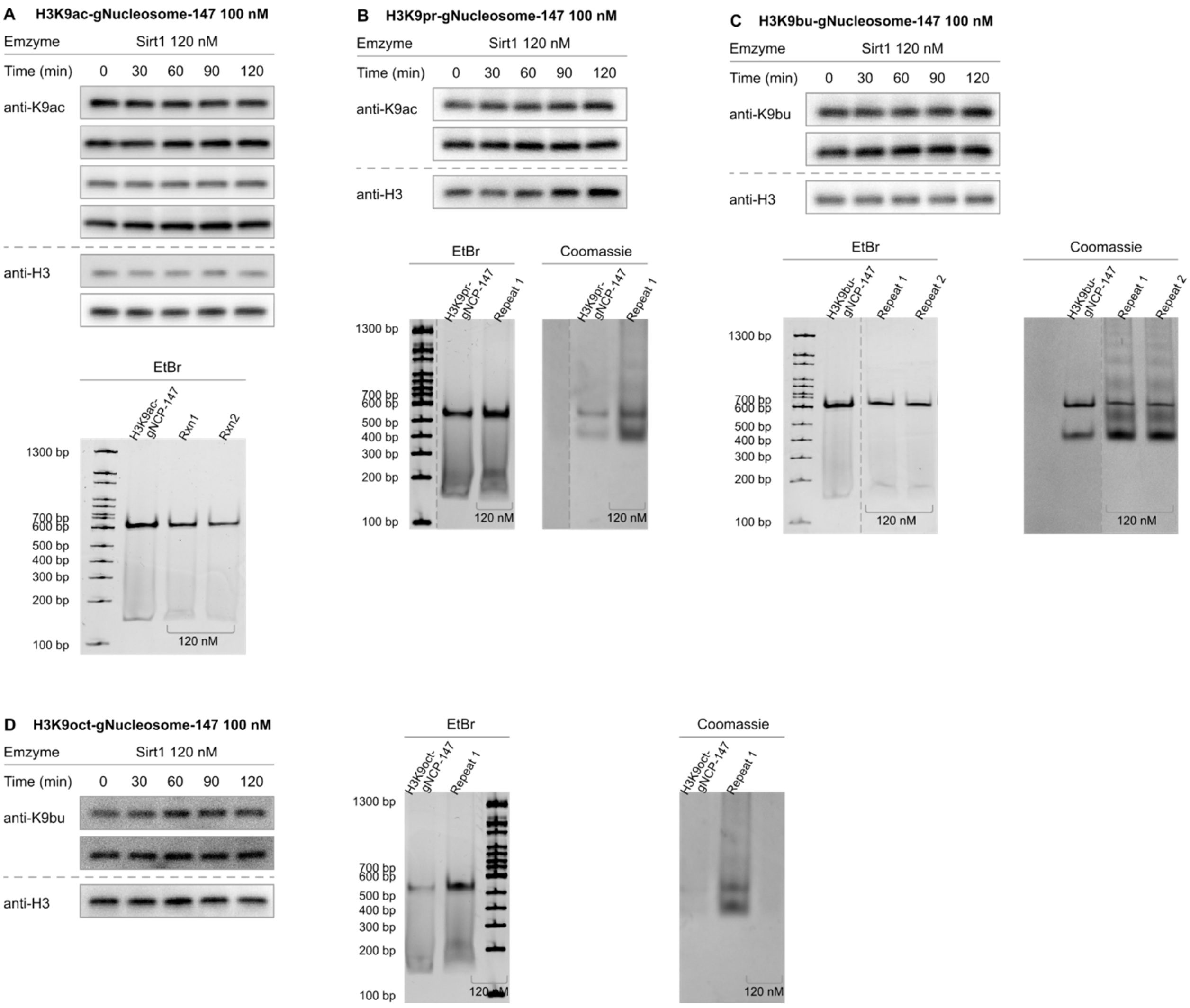

**Figure S16. Western blot measurement of Sirt1 activity by length of H3K9 acylation carbon chain.** (**A**) Western blots (top; anti-H3K9ac) and native TBE gels (bottom) of Sirt1 deacetylation assays with 147bp H3K9acetyl (H3K9ac) nucleosomes. (**B**) Western blots (top; anti-H3K9ac antibody) and native TBE gels (bottom) of Sirt1 deacetylation assays with 147 bp H3K9propionyl (H3K9pr) nucleosomes. (**C**) Western blots (top; anti-H3K9bu) and native TBE gels (bottom) of Sirt1 deacetylation assays with 147 bp H3K9butyryl (H3K9bu) nucleosomes. (**D**) Western blots (left; anti-H3K9but) and native TBE gels (right) of Sirt1 deacetylation assays with 147 bp H3K9octanoyl (H3K9oct) nucleosomes.

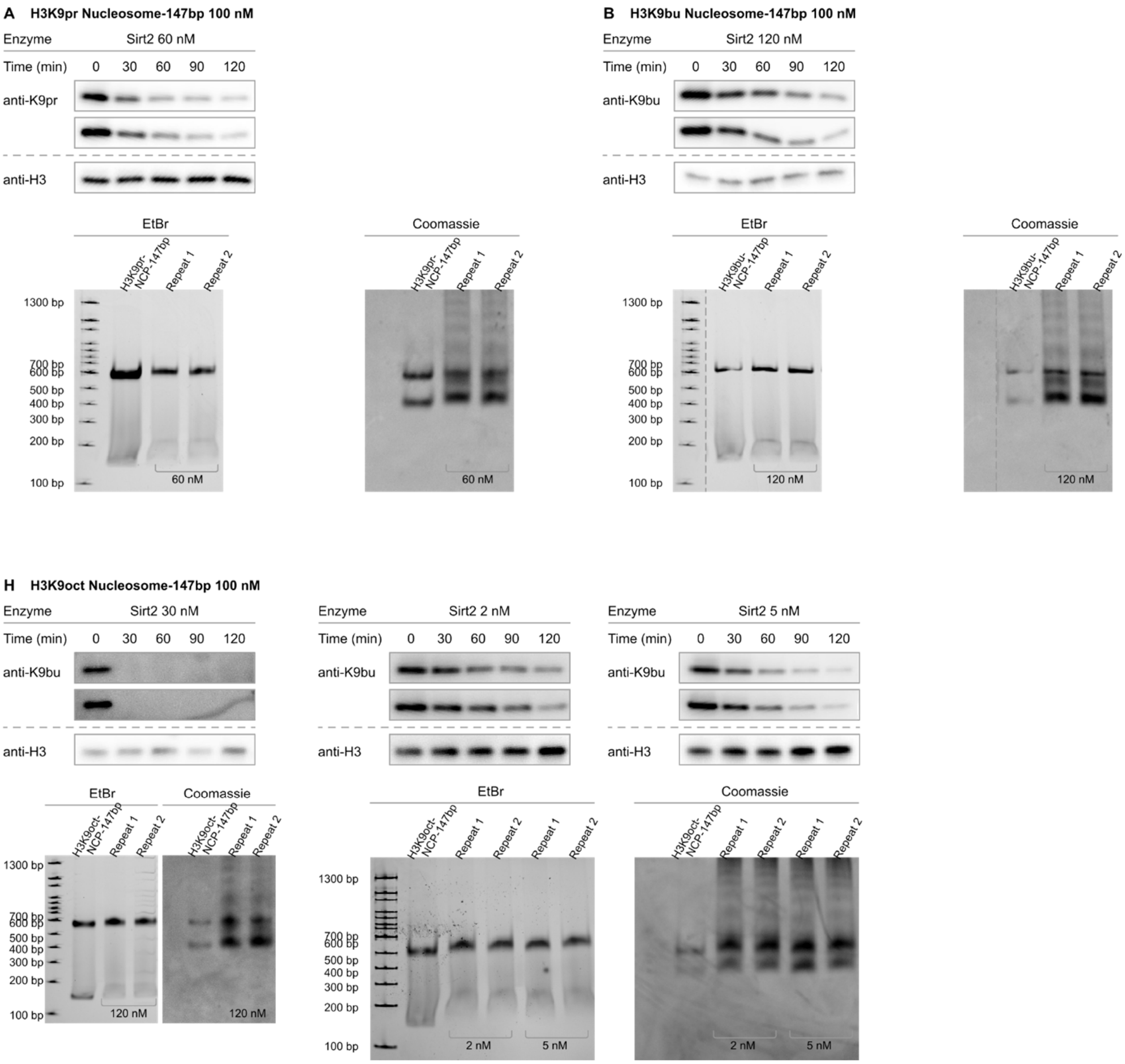

**Figure S17. Western blot measurement of Sirt2 activity by length of H3K9 acylation carbon chain.** (**A**) Western blots (top; anti-H3K9ac antibody) and native TBE gels (bottom) of Sirt2 deacetylation assays with H3K9propionyl (H3K9pr) nucleosomes. (**B**) Western blots (top) and native TBE gels (bottom) of Sirt2 deacetylation assays with H3K9butyryl (H3K9bu) nucleosomes. (**C**) Western blots (top; anti-H3K9bu) and native TBE gels (bottom) of Sirt2 deacetylation assays with H3K9octanoyl (H3K9oct) nucleosomes.

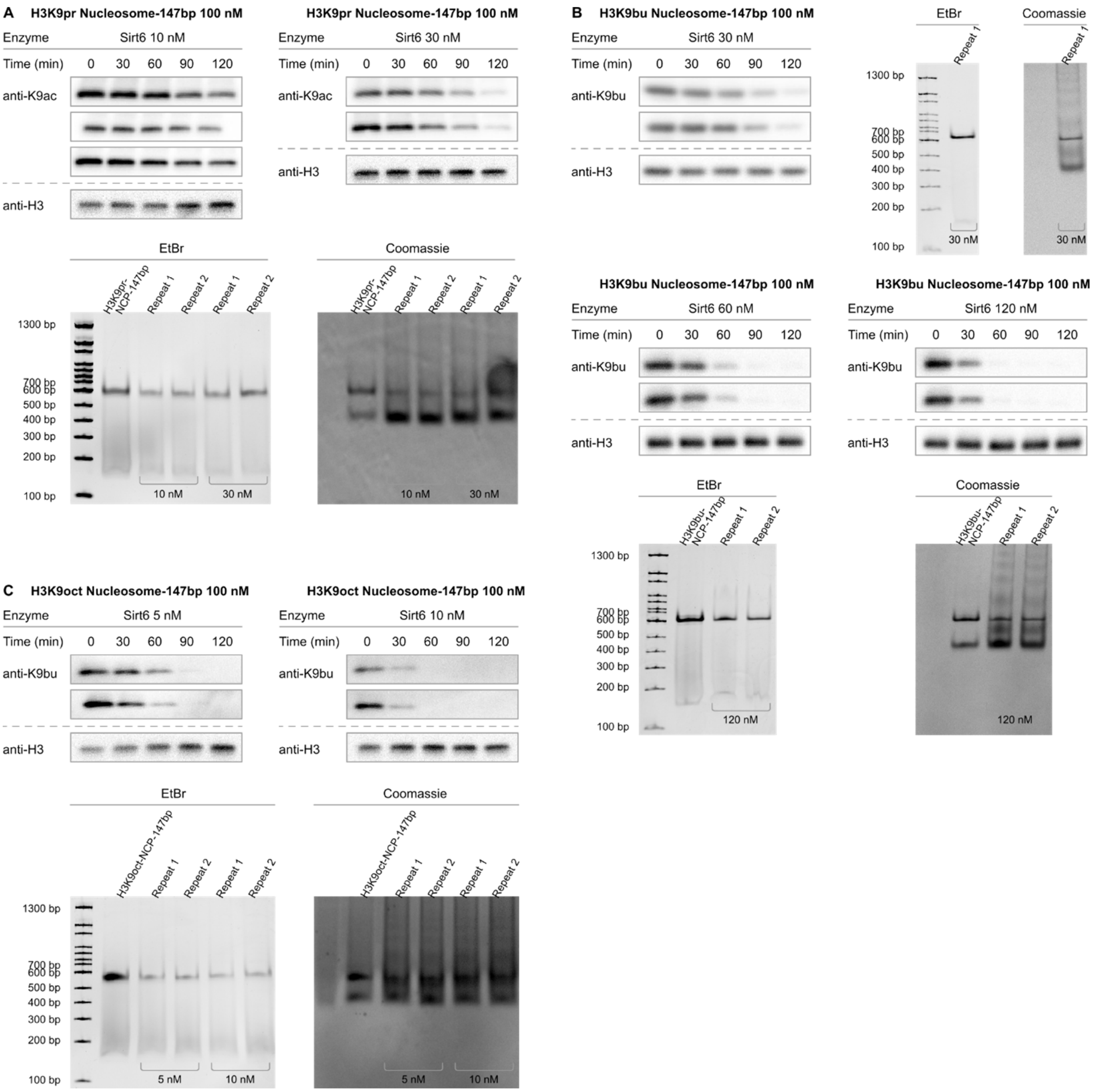

**Figure S18. Western blot measurement of Sirt6 activity by length of H3K9 acylation carbon chain.** (**A**) Western blots (top; anti-H3K9ac antibody) and native TBE gels (bottom) of Sirt6 deacetylation assays with 147 bp H3K9propionyl (H3K9pr) nucleosomes. (**B**) Western blots (top, middle anti-H3K9bu) and native TBE gels (right, bottom) of Sirt6 deacetylation assays with 147 bp H3K9butyryl (H3K9bu) nucleosomes. (**C**) Western blots (top; anti-H3K9bu) and native TBE gels (bottom) of Sirt6 deacetylation assays with 147 bp H3K9octanoyl (H3K9oct) nucleosomes.

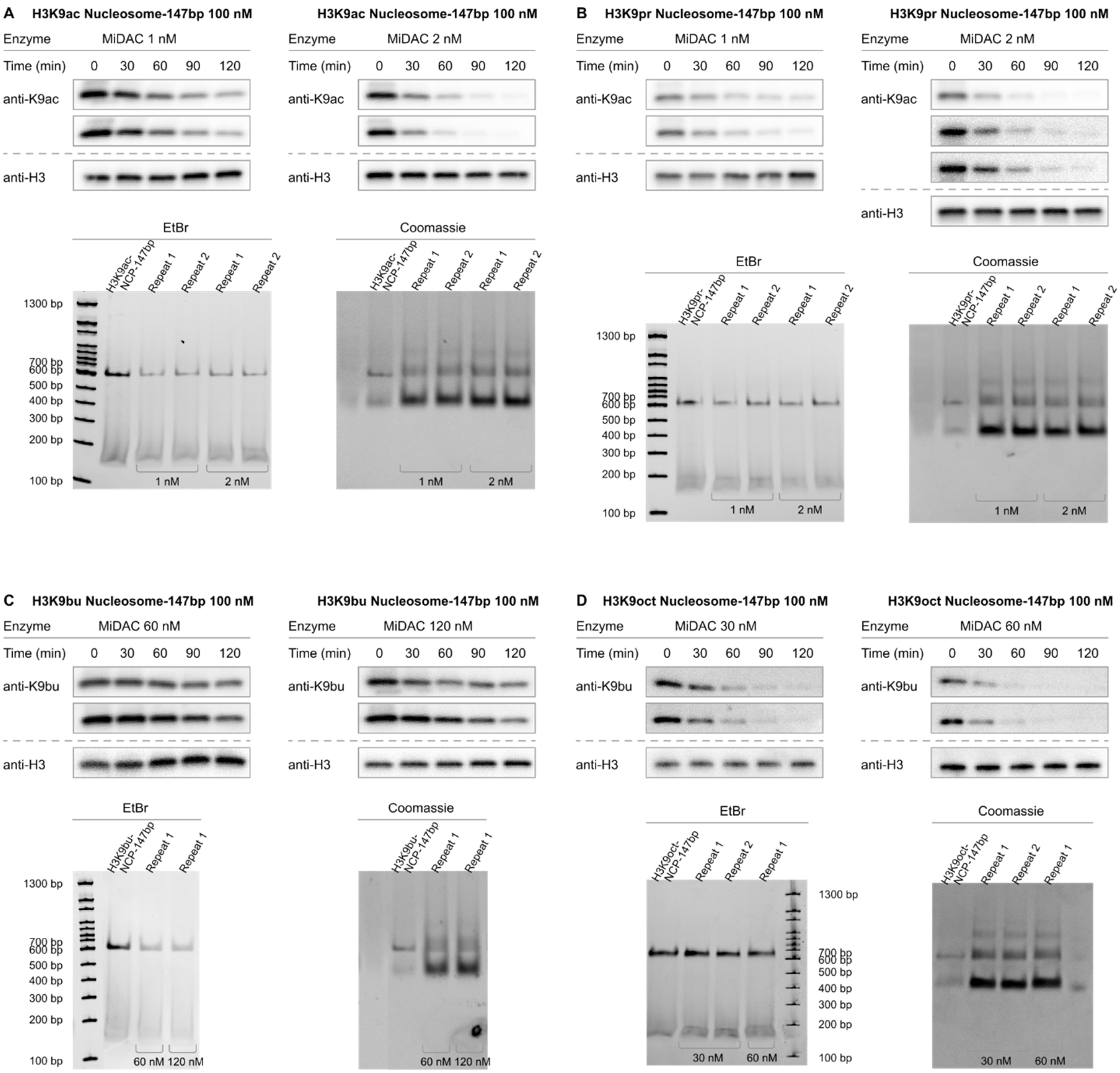

**Figure S19. Western blot measurement of MiDAC activity by length of H3K9 acylation carbon chain.** (**A**) Western blots (top; anti-H3K9ac) and native TBE gels (bottom) of MiDAC deacetylation assays with 147 bp H3K9acetyl (H3K9ac) nucleosomes. (**B**) Western blots (top; anti-H3K9ac antibody) and native TBE gels (bottom) of MiDAC deacetylation assays with 147 bp H3K9propionyl (H3K9pr) nucleosomes. (**C**) Western blots (top; anti-H3K9bu) and native TBE gels (bottom) of MiDAC deacetylation assays with 147 bp H3K9butyryl (H3K9bu) nucleosomes. (**D**) Western blots (top; anti-H3K9bu) and native TBE gels (bottom) of MiDAC deacetylation assays with 147 bp H3K9octanoyl (H3K9oct) nucleosomes.

| Acylated Nucleosome | Sirt 1 | Sirt2 | Sirt6 | MiDAC |
|---|---|---|---|---|
| H3K9ac | <0.002 | 0.026±0.0015* | 0.075±0.0044 | 1.4±0.046 |
| H3K9pr | <0.002 | 0.036±0.0022 | 0.041±0.0051 | 1.9±0.13 |
| H3K9bu | <0.002 | 0.013±0.00050 | 0.044±0.0049 | 0.0077±0.00053 |
| H3K9oct | <0.002 | 0.49±0.026 | 0.46±0.063 | 0.081±0.0051 |

**Table S6. Calculated V/[E] values for HDAC activity by length of H3K9 acylation carbon chain.** Average V/[E] ± SEM values for Sirt1, Sirt2, Sirt6 and MiDAC deacylation of deacylation of nucleosomes with linear acylations of increasing length. Values marked with asterisks are re-printed from prior reports.[6]

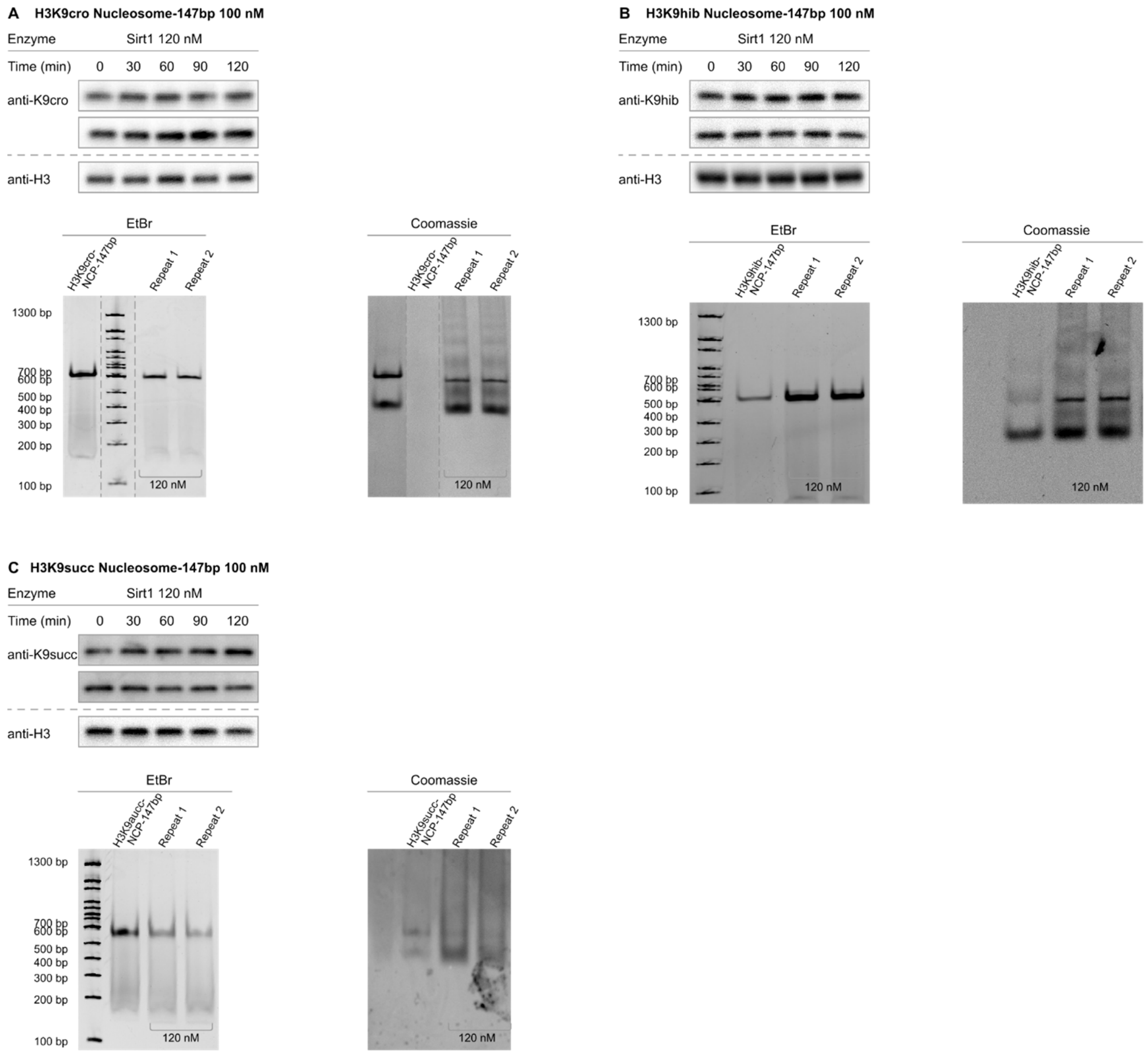

**Figure S20. Western blot measurement of Sirt1 activity toward four carbon acylations of H3K9.** (**A**) Western blots (top; anti-H3K9cro) and native TBE gels (bottom) of Sirt1 deacetylation assays with 147 bp H3K9crotonyl (H3K9cro) nucleosomes. (**B**) Western blots (top; anti-H3K9hib) and native TBE gels (bottom) of Sirt1 deacetylation assays with 147 bp H3K9anti-hydroxyisobutyryl nucleosomes. (**C**) Western blots (top; anti-H3K9succ) and native TBE gels (bottom) of Sirt1 deacetylation assays with 147 bp H3K9succinyl nucleosomes.

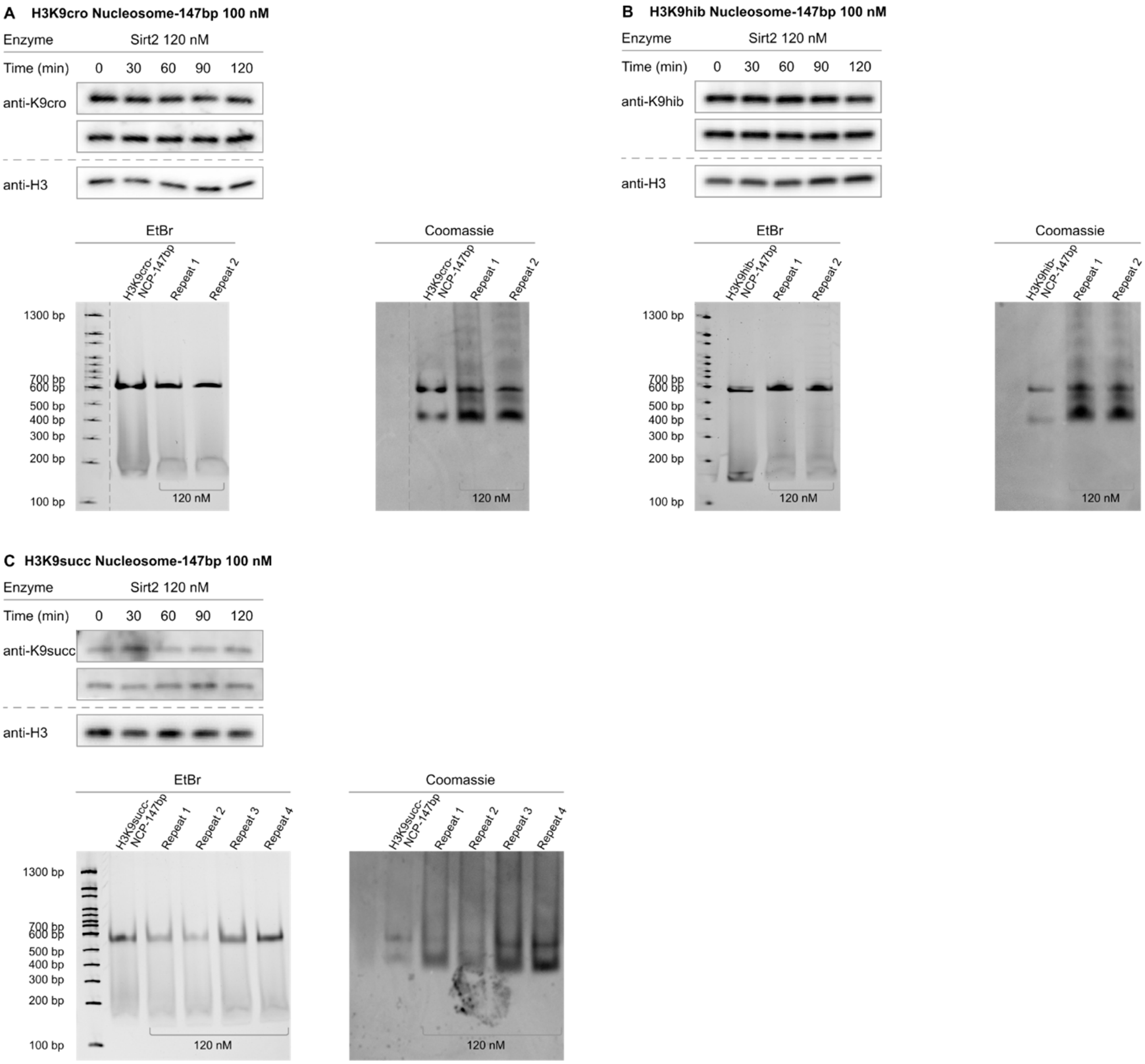

**Figure S21. Western blot measurement of Sirt2 activity toward four carbon acylations of H3K9.** (**A**) Western blots (top; anti-H3K9cro) and native TBE gels (bottom) of Sirt2 deacetylation assays with 147 bp H3K9crotonyl (H3K9cro) nucleosomes. (**B**) Western blots (top; anti-H3K9hib) and native TBE gels (bottom) of Sirt2 deacetylation assays with 147 bp H3K9anti-hydroxyisobutyryl nucleosomes. (**C**) Western blots (top; anti-H3K9succ) and native TBE gels (bottom) of Sirt6 deacetylation assays with 147 bp H3K9succinyl nucleosomes.

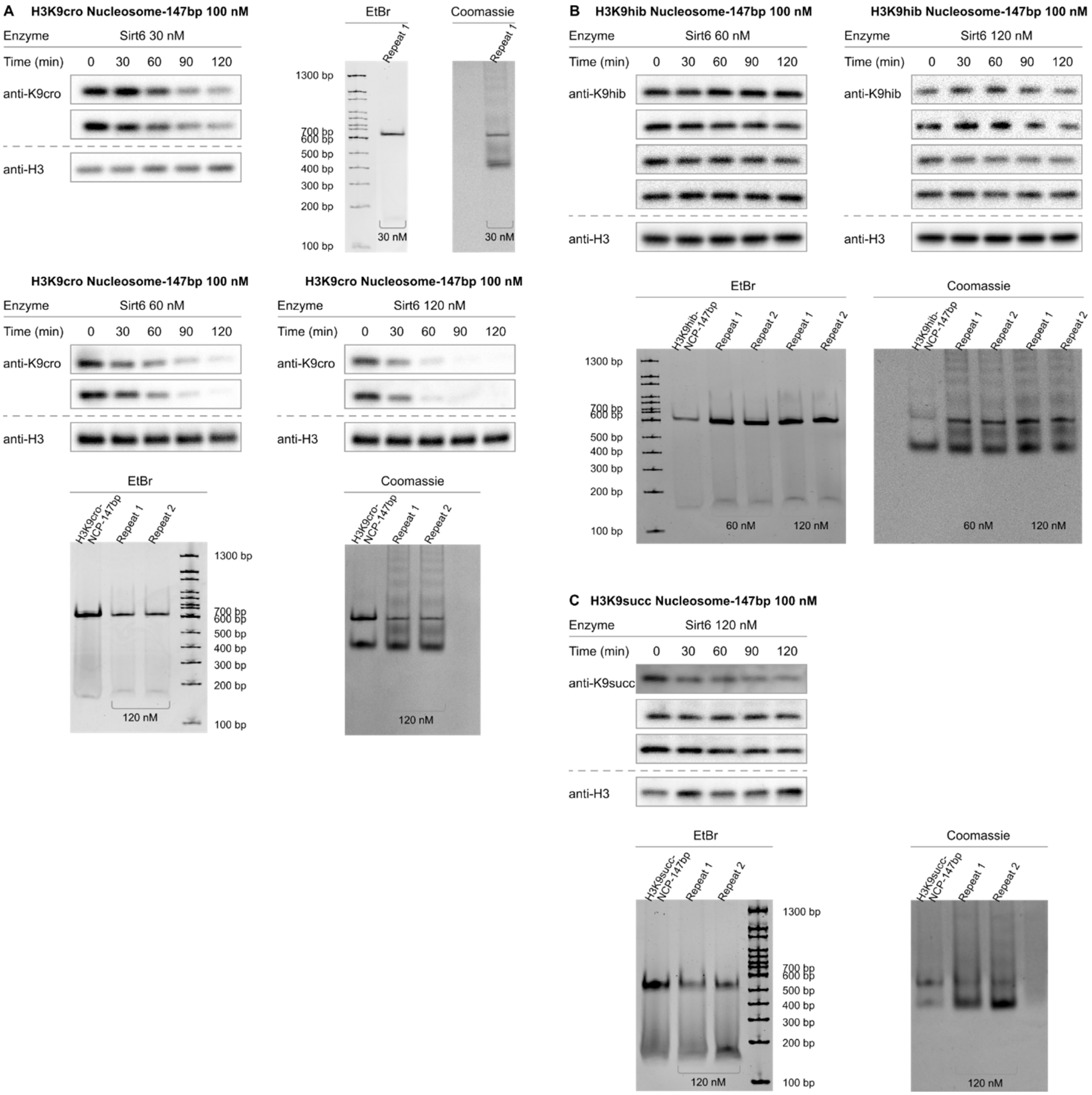

**Figure S22. Western blot measurement of Sirt6 activity toward four carbon acylations of H3K9.** (**A**) Western blots (left, middle; anti-H3K9cro) and native TBE gels (right) of Sirt6 deacetylation assays with 147 bp H3K9crotonyl (H3K9cro) nucleosomes. (**B**) Western blots (top; anti-H3K9hib) and native TBE gels (bottom) of Sirt6 deacetylation assays with 147 bp H3K9anti-hydroxyisobutyryl nucleosomes. (**C**) Western blots (top; anti-H3K9succ) and native TBE gels (bottom) of Sirt6 deacetylation assays with 147 bp H3K9succinyl nucleosomes.

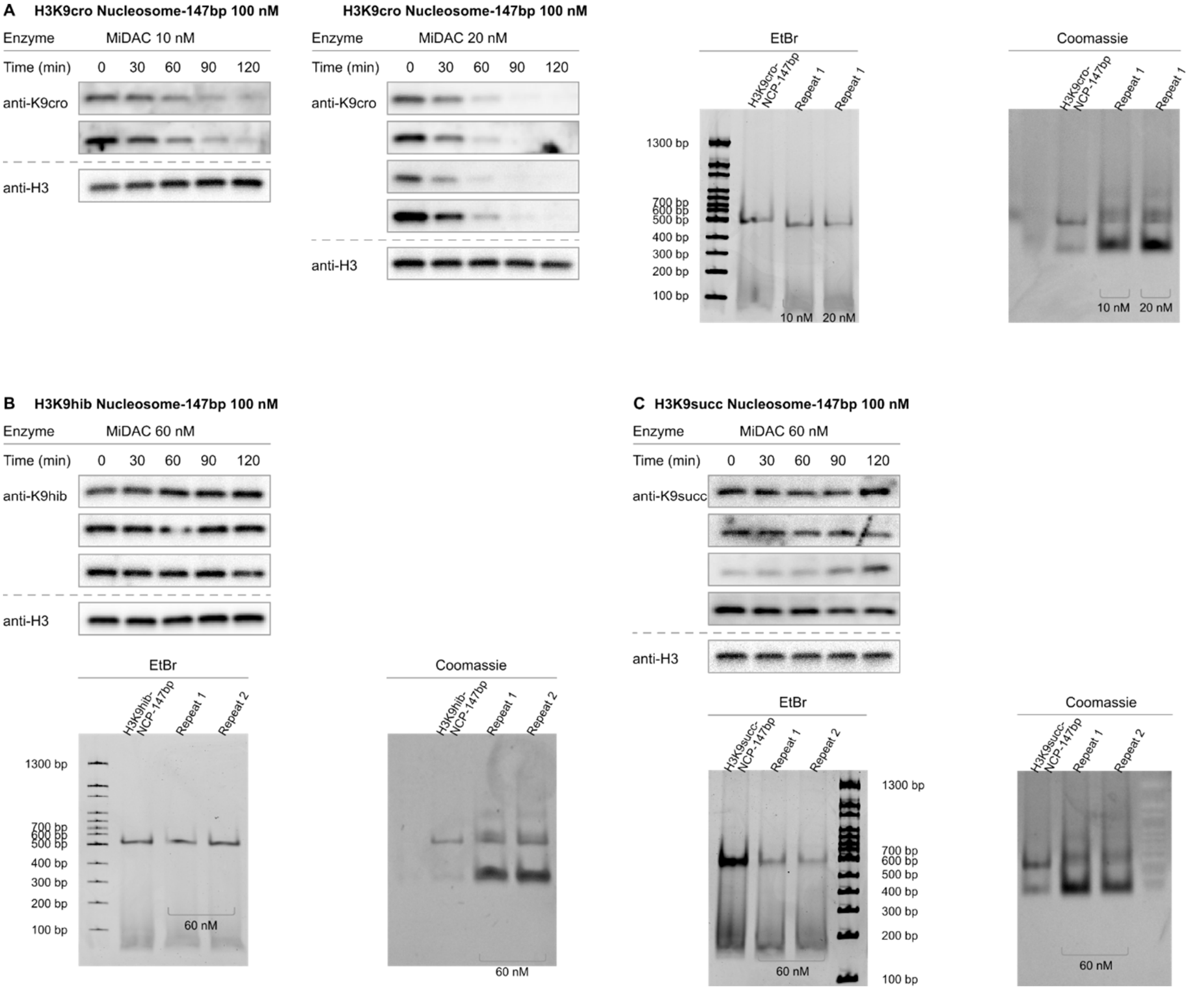

**Figure S23. Western blot measurement of MiDAC activity toward four carbon acylations of H3K9.** (**A**) Western blots (left; anti-H3K9cro) and native TBE gels (right) of MiDAC deacetylation assays with 147 bp H3K9crotonyl (H3K9cro) nucleosomes. (**B**) Western blots (top; anti-H3K9hib) and native TBE gels (bottom) of MiDAC deacetylation assays with 147 bp H3K9anti-hydroxyisobutyryl (H3K9hib) nucleosomes. (**C**) Western blots (top; anti-H3K9hib) and native TBE gels (bottom) of MiDAC deacetylation assays with 147 bp H3K9succinyl (H3K9succ) nucleosomes.

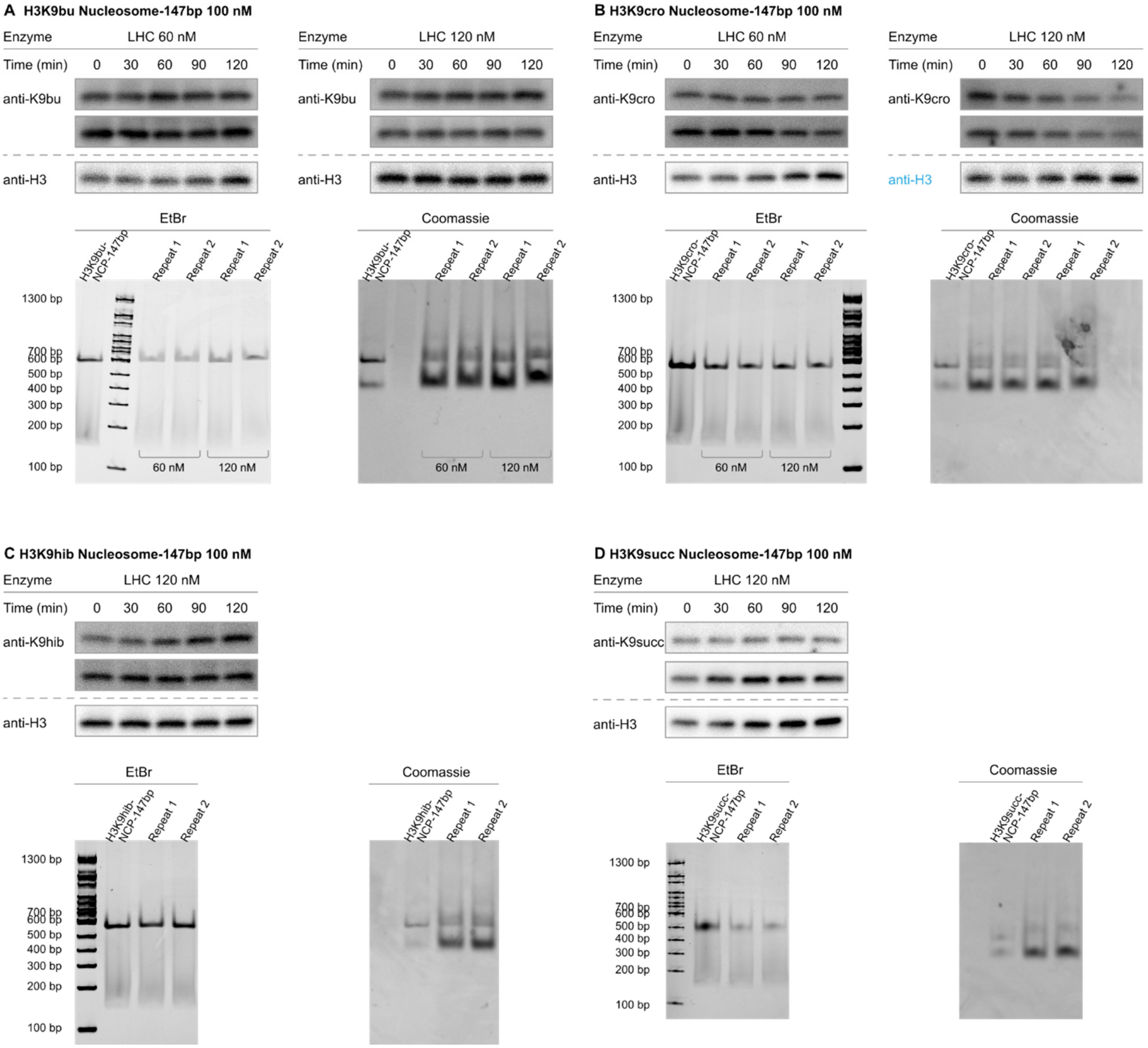

**Figure S24. Western blot measurement of LHC activity toward four carbon acylations of H3K9.** (**A**) Western blots (top; anti-H3K9bu) and native TBE gels (bottom) of LHC deacetylation assays with 147 bp H3K9butyryl (H3K9bu) nucleosomes. (**B**) Western blots (top; anti-H3K9cro) and native TBE gels (bottom) of LHC deacetylation assays with 147 bp H3K9crotonyl (H3K9cro) nucleosomes. (**C**) Western blots (top; anti-H3K9hib) and native TBE gels (bottom) of LHC deacetylation assays with 147 bp H3K9anti-hydroxyisobutyryl (H3K9hib) nucleosomes. (**D**) Western blots (top; anti-H3K9hib) and native TBE gels (bottom) of LHC deacetylation assays with 147 bp H3K9succinyl (H3K9succ) nucleosomes.

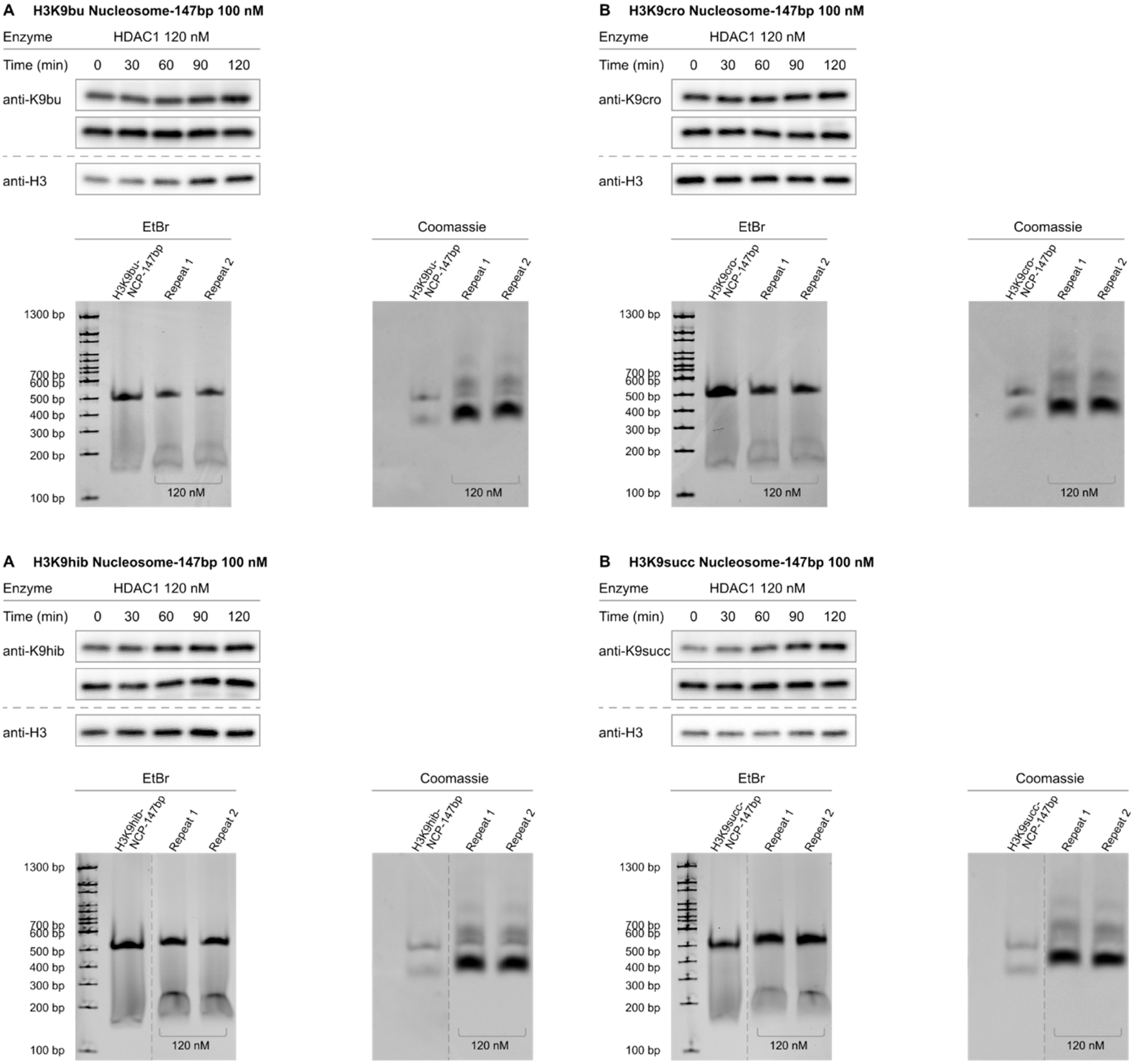

**Figure S25. Western blot measurement of free HDAC1 activity toward four carbon acylations of H3K9.** (**A**) Western blots (top; anti-H3K9bu) and native TBE gels (bottom) of HDAC deacetylation assays with 147 bp H3K9butyryl (H3K9bu) nucleosomes. (**B**) Western blots (top; anti-H3K9cro) and native TBE gels (bottom) of HDAC deacetylation assays with 147 bp H3K9crotonyl (H3K9cro) nucleosomes. (**C**) Western blots (top; anti-H3K9hib) and native TBE gels (bottom) of HDAC deacetylation assays with 147 bp H3K9anti-hydroxyisobutyryl (H3K9hib) nucleosomes. (**D**) Western blots (top; anti-H3K9hib) and native TBE gels (bottom) of HDAC deacetylation assays with 147 bp H3K9succinyl (H3K9succ) nucleosomes.

| Acylated Nucleosome | Sirt 1 | Sirt2 | Sirt6 | MiDAC | LHC | HDAC1 |
|---|---|---|---|---|---|---|
| H3K9ac | <0.002 | 0.026±0.0015* | 0.075±0.0044 | 1.4±0.046 | 0.021±0.0011 | <0.002 |
| H3K9bu | <0.002 | 0.013±0.00050 | 0.044±0.0049 | 0.0077±0.00053 | <0.002 | <0.002 |
| H3K9cro | <0.002 | <0.002 | 0.033±0.0021 | 0.16±0.011 | 0.0086±0.00039 | <0.002 |
| H3K9hib | <0.002 | <0.002 | 0.0013±0.00049 | <0.002 | <0.002 | <0.002 |
| H3K9succ | <0.002 | <0.002 | 0.0044±0.00050 | <0.002 | <0.002 | <0.002 |

**Table S7. Calculated V/[E] values for HDAC activity toward four carbon acylations of H3K9.** Average V/[E] ± SEM values for Sirt1, Sirt2, Sirt6, MiDAC, LHC and free HDAC1 deacylation of 147 bp nucleosomes with different 4 carbon acylations. Western blot bands were quantified by ImageJ (imagej.nih.gov/ij/). Intensity values were normalized to intensity at t=0 and then fit to a single-phase exponential decay curve with constrain Y0=1, Plateau=0 (GraphPad Prism 10). Values marked with asterisks are re-printed from prior reports.[6]

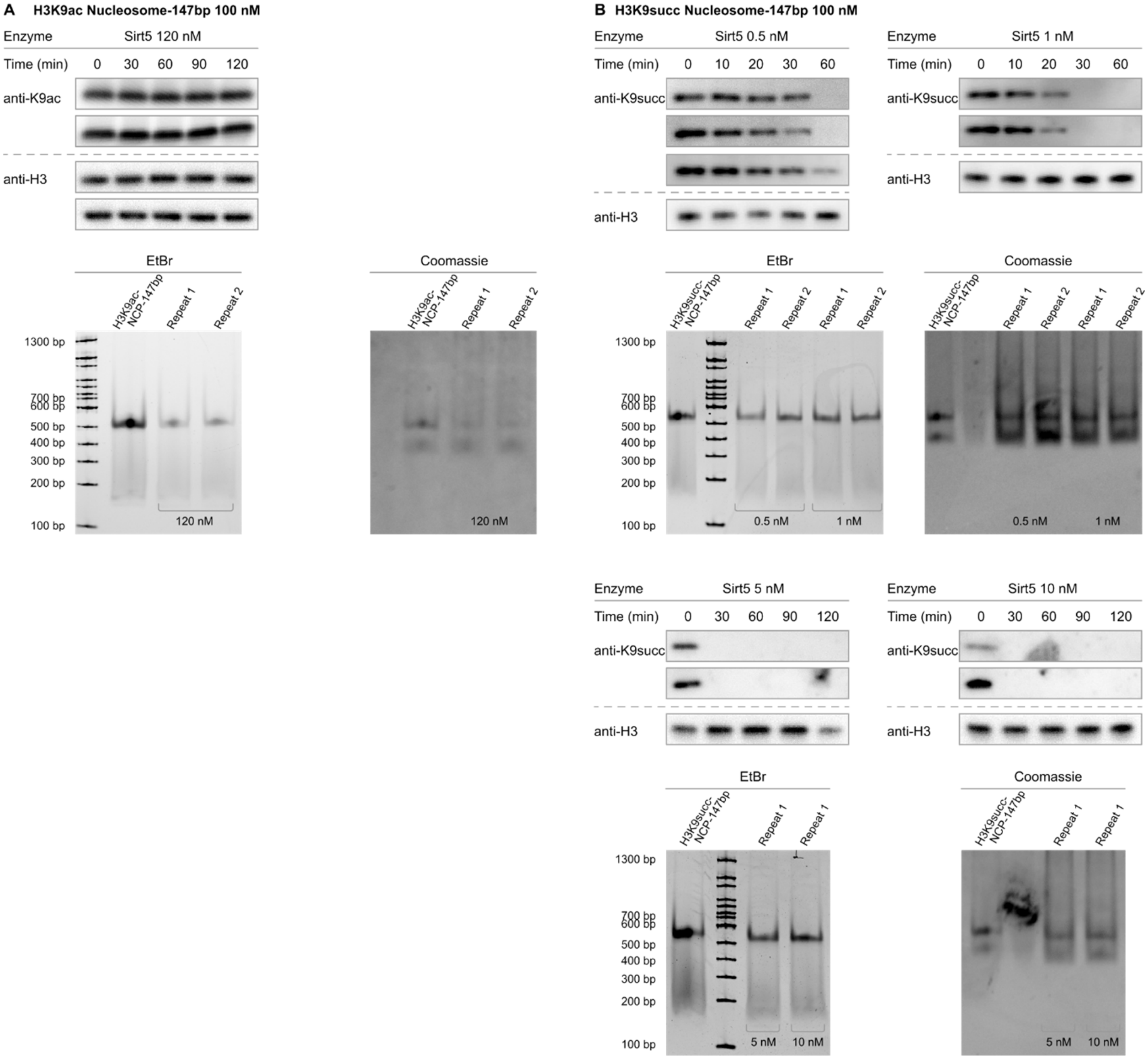

**Figure S26. Western blot measurement of Sirtuin5 activity toward acetylated and succinylated H3K9.** (**A**) Western blots (top; anti-H3K9ac) and native TBE gels (bottom) of Sirt5 deacetylation assays with 147 bp H3K9acetul (H3K9ac) nucleosomes. (**B**) Western blots (top, middle; anti-H3K9succ) and native TBE gels (middle, bottom) of Sirt5 deacetylation assays with 147 bp H3K9succinyl (H3K9succ) nucleosomes.

| Acylated Nucleosome | Sirt 1 | Sirt2 | Sirt6 | MiDAC | LHC | HDAC1 | Sirt5 |
|---|---|---|---|---|---|---|---|
| H3K9ac | <0.002 | 0.026±0.0015* | 0.075±0.0044 | 1.4±0.046 | 0.021±0.0011 | <0.002 | <0.002 |
| H3K9succ | <0.002 | <0.002 | 0.0044±0.00050 | <0.002 | <0.002 | <0.002 | 5.7±0.64 |

**Table S8. Calculated V/[E] values for HDAC activity toward H3K9 succinylation.** Average V/[E] ± SEM values for Sirt1, Sirt2, Sirt6, MiDAC, LHC, free HDAC1 and Sirt5 deacylation of 147 bp nucleosomes with H3K9 succinylation. Western blot bands were quantified by ImageJ (imagej.nih.gov/ij/). Intensity values were normalized to intensity at t=0 and then fit to a single-phase exponential decay curve with constrain Y0=1, Plateau=0 (GraphPad Prism 10). Values marked with asterisks are re-printed from prior reports.[6]

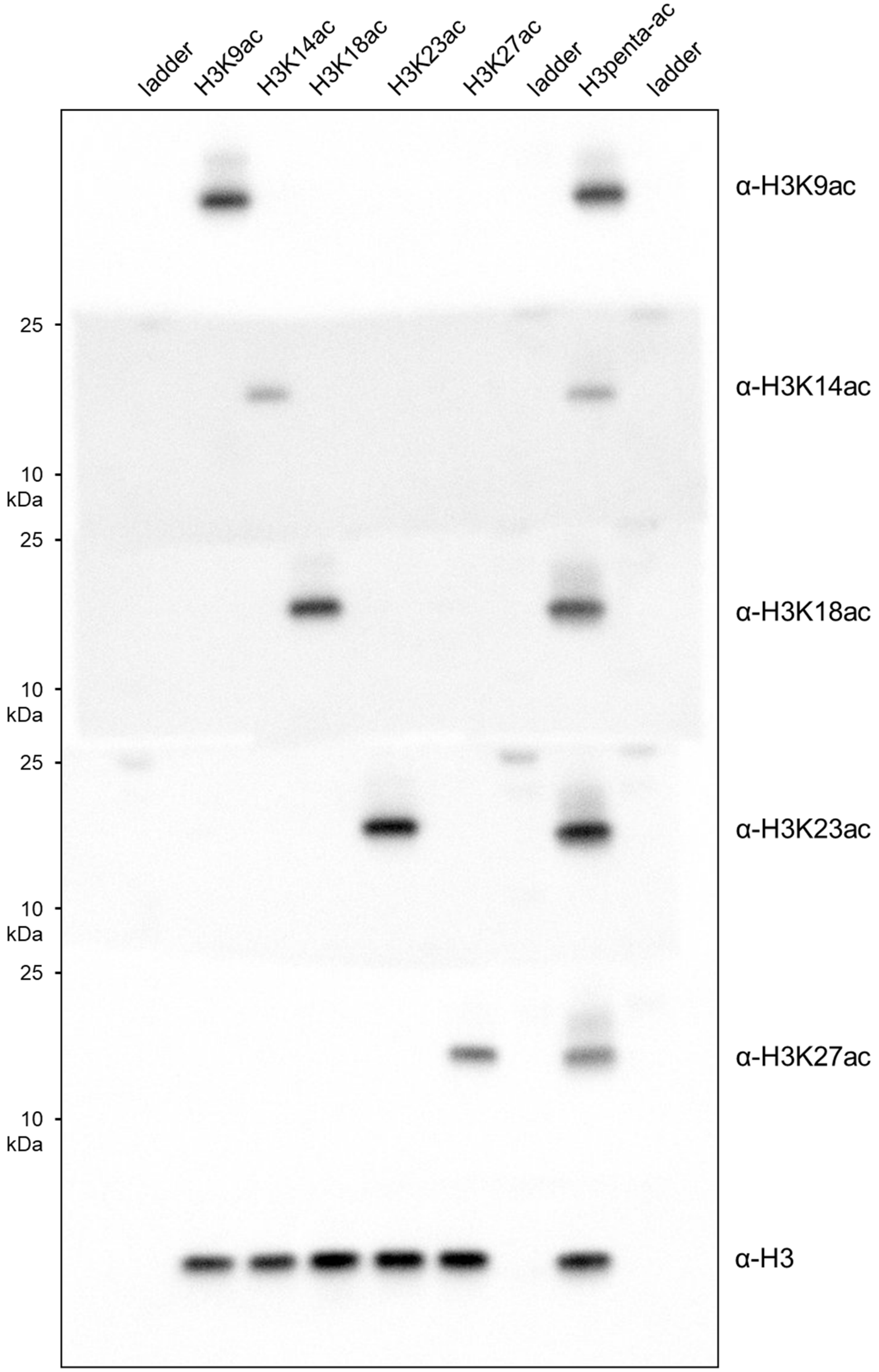

**Figure S27. Validation of H3Kac single site antibody specificity.** Western blot analysis of synthetic nucleosomes with site-specific acetylations at (left-to-right) H3K9, H3K14, H3K18, H3K23, H3K27, or all five positions (H3penta-ac) with site-specific antibodies toward (top-to-bottom) H3K9ac, H3K14ac, H3K18ac, H3K23ac or H3K27ac. Molecular weight markers are visible in blots for H3K14ac, H3K18ac, H3K23ac and H3K27ac.

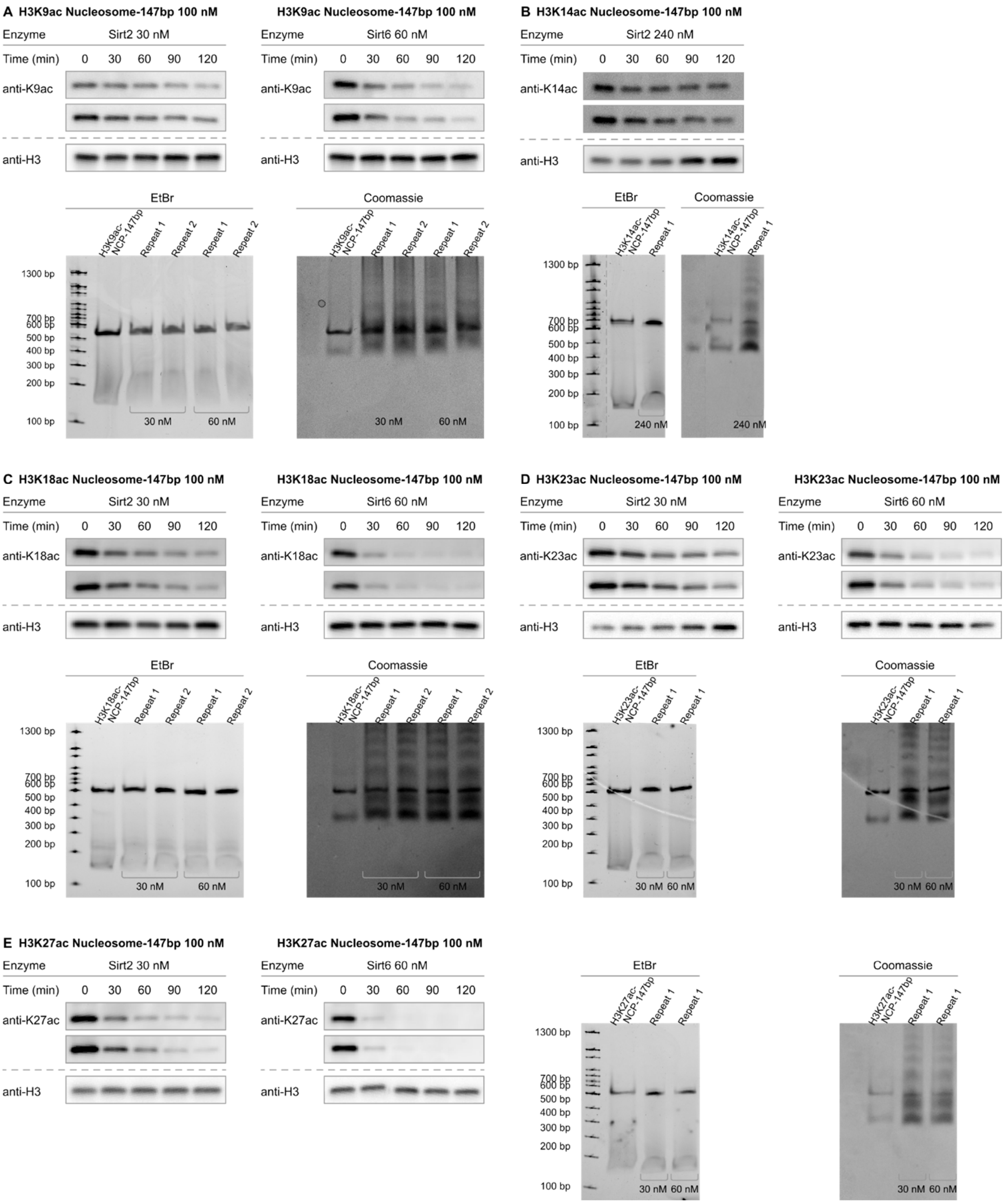

**Figure S28. Western blot measurement of Sirt2 activity toward mono-acetylated nucleosomes.** (**A**) Western blots (top) and native TBE gels (bottom) of Sirt2 deacetylation assays with 147 bp H3K9acetyl (H3K9ac) nucleosomes. (**B**) Western blots (top) and native TBE gels (bottom) of Sirt2 deacetylation assays with 147 bp H3K14acetyl (H3K14ac) nucleosomes. (**C**) Western blots (top) and native TBE gels (bottom) of Sirt2 deacetylation assays with 147 bp H3K18acetyl (H3K8ac) nucleosomes. (**D**) Western blots (top) and native TBE gels (bottom) of Sirt2 deacetylation assays with 147 bp H3K13acetyl

(H3K13ac) nucleosomes. (**A**) Western blots (left) and native TBE gels (right) of Sirt2 deacetylation assays with 147 bp H3K27acetyl (H3K27ac) nucleosomes.

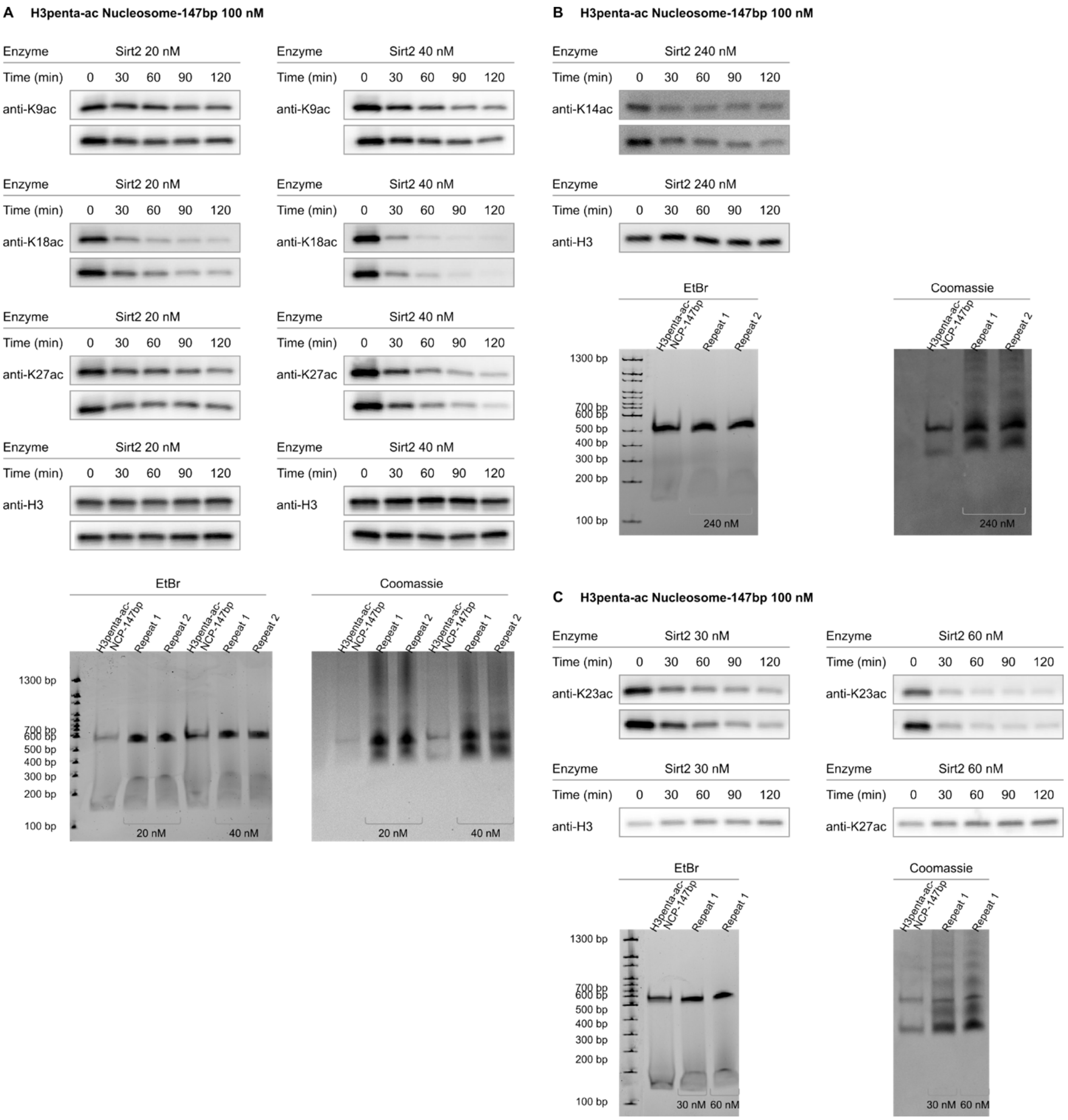

**Figure S29. Western blot measurement of Sirt2 activity toward penta-acetylated nucleosomes.** Western blots (top; anti-H3K9ac, anti-H3K18ac, anti-H3K27ac) and native TBE gels (bottom) of Sirt2 deacetylation assays with 147 bp H3K9ac/K14ac/K18ac/K23ac/K27ac (H3Kpenta-ac) nucleosomes. (**B**) Western blots (top; anti-H3K14ac) and native TBE gels (bottom) of Sirt2 deacetylation assays with 147 bp H3K9ac/K14ac/K18ac/K23ac/K27ac (H3Kpenta-ac) nucleosomes. (**C**) Western blots (top; anti-H3K23ac, anti-H3K27ac) and native TBE gels (bottom) of Sirt2 deacetylation assays with 147 bp H3K9ac/K14ac/K18ac/K23ac/K27ac (H3Kpenta-ac) nucleosomes.

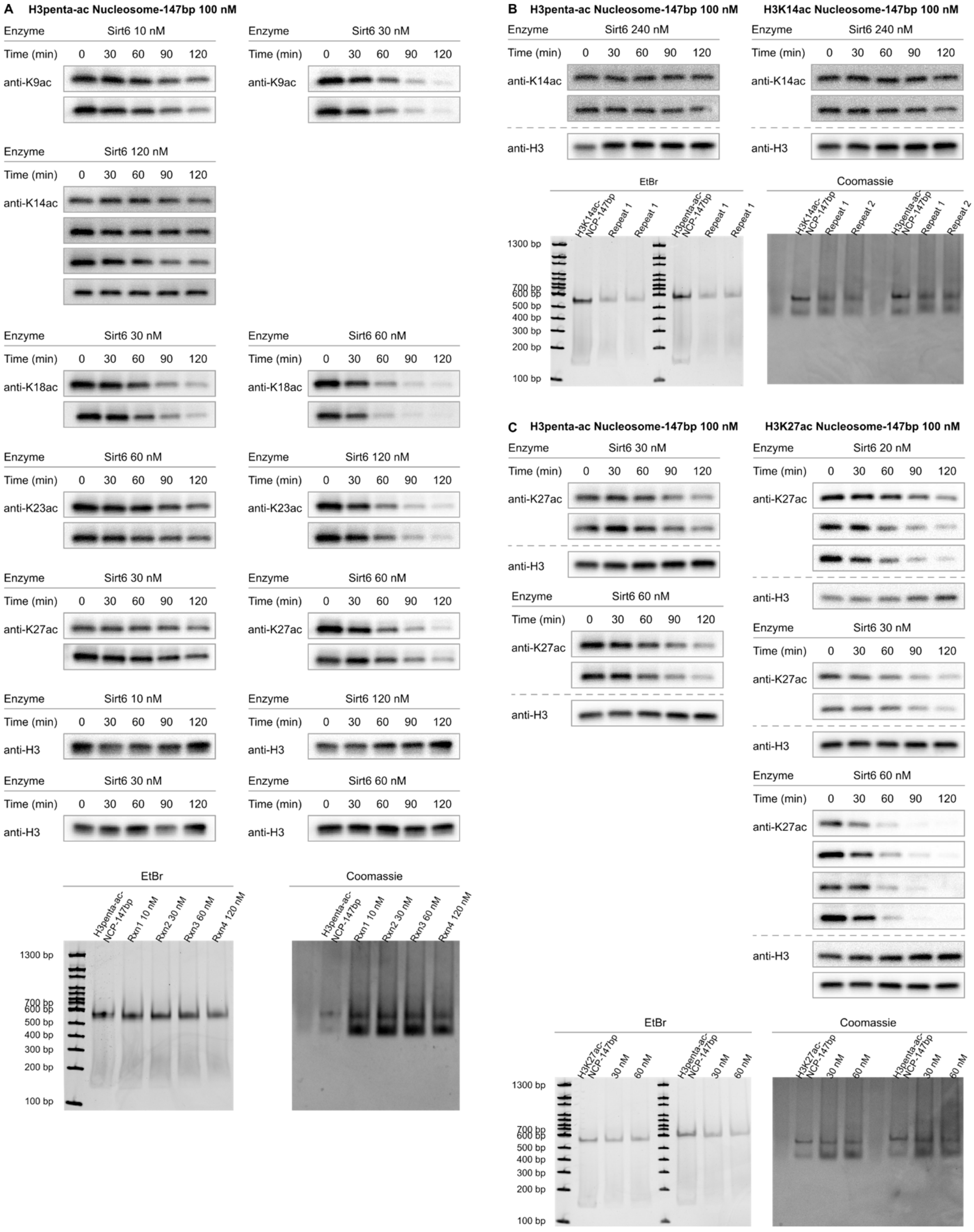

**Figure S30. Western blot measurement of Sirt6 activity toward mono- and penta-acetylated nucleosomes.** (**A**) Western blots (top; anti-H3K9ac, anti-H3K14ac, anti-H3K18ac, anti-H3K23ac, anti-H3K27ac) and native TBE gels (bottom) of Sirt6 deacetylation assays with 147 bp

H3K9ac/K14ac/K18ac/K23ac/K27ac (H3Kpenta-ac) nucleosomes. (**B**) Western blots (top; anti-H3K14ac) and native TBE gels (bottom) of parallel Sirt6 deacetylation assays with 147 bp H3K9ac/K14ac/K18ac/K23ac/K27ac (H3Kpenta-ac) nucleosomes and 147 bp H3K14acetyl (H3K14ac) nucleosomes. (**C**) Western blots (top; anti-H3K27ac) and native TBE gels (bottom) of parallel Sirt6 deacetylation assays with 147 bp H3K9ac/K14ac/K18ac/K23ac/K27ac (H3Kpenta-ac) nucleosomes (left) and 147 bp H3K27acetyl (H3K27ac) nucleosomes.

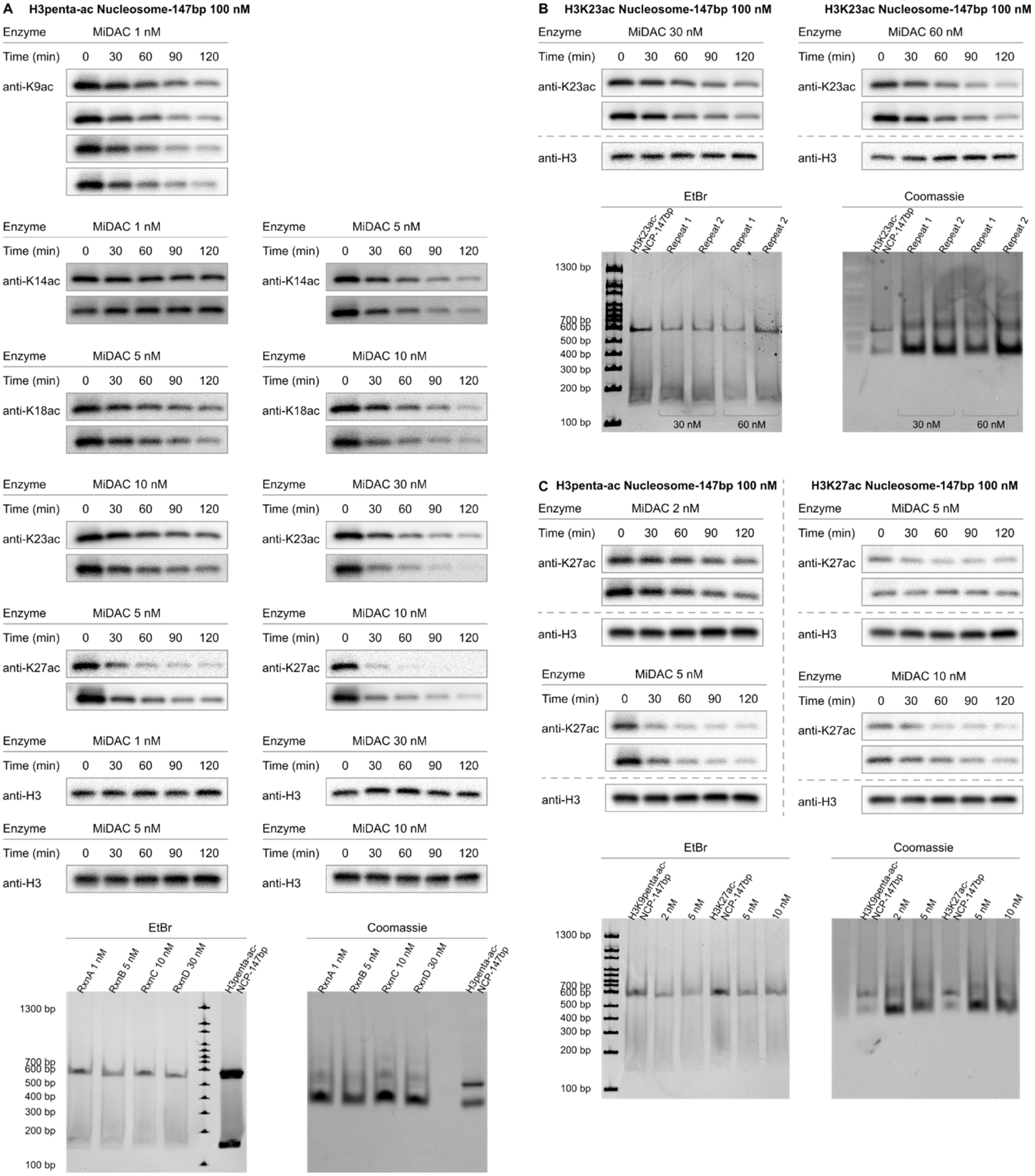

**Figure S31. Western blot measurement of MiDAC activity toward mono- and penta-acetylated nucleosomes.** (**A**) Western blots (top; anti-H3K9ac, anti-H3K14ac, anti-H3K18ac, anti-H3K23ac, anti-H3K27ac) and native TBE gels (bottom) of MiDAC deacetylation assays with 147 bp H3K9ac/K14ac/K18ac/K23ac/K27ac (H3Kpenta-ac) nucleosomes. (**B**) Western blots (top; anti-H3K23ac) and native TBE gels (bottom) of MiDAC deacetylation assays with 147 bp H3K23acetyl (H3K23ac) nucleosomes. (**C**) Western blots (top; anti-H3K23ac) and native TBE gels (bottom) of MiDAC deacetylation assays with 147 bp H3K9ac/K14ac/K18ac/K23ac/K27ac (H3Kpenta-ac) nucleosomes (left) and 147 bp H3K27acetyl (H3K27ac) nucleosomes.

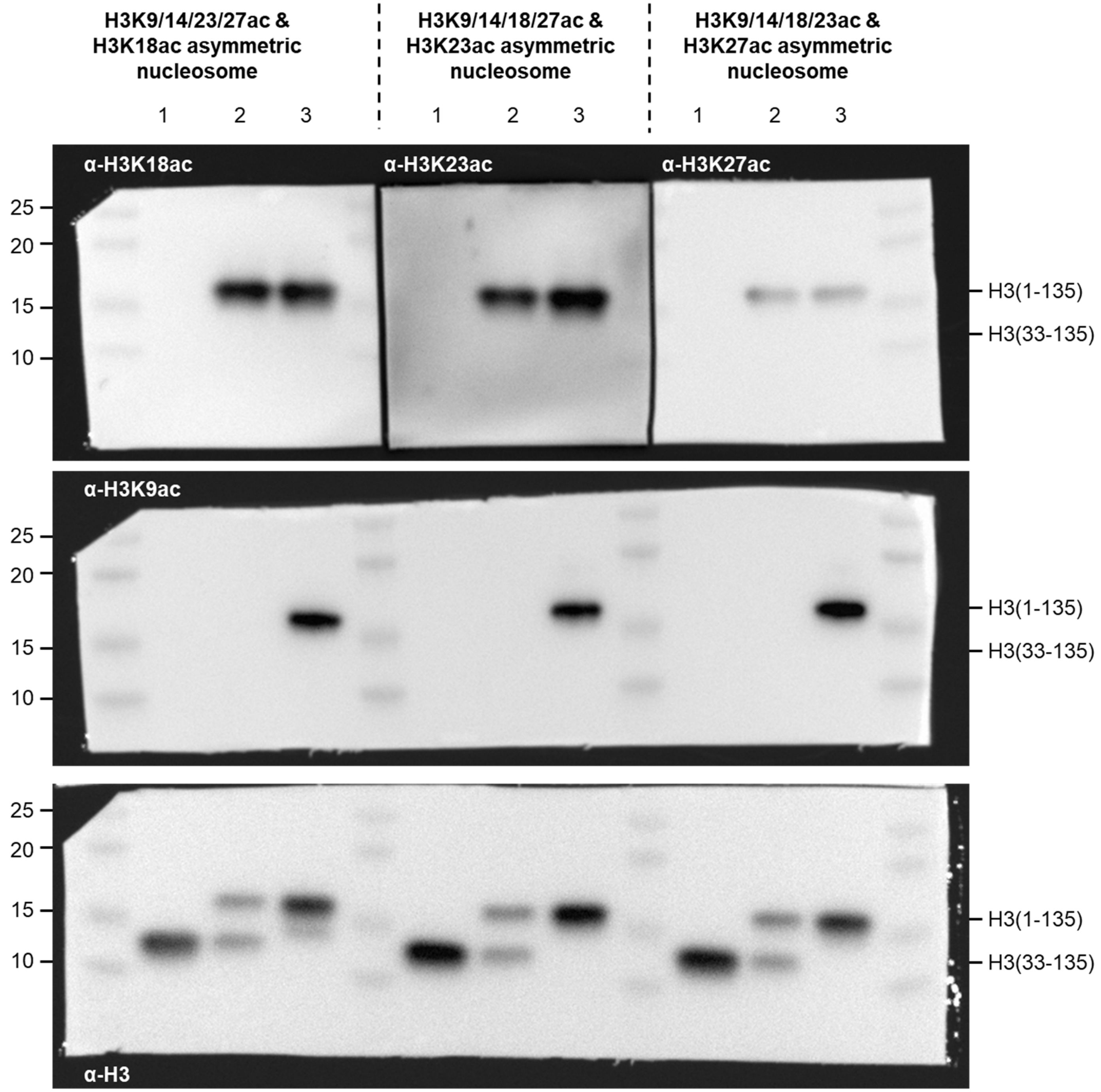

**Figure S32. Western blot characterization of asymmetric mono/tetra-acetylated nucleosomes.** Anti-H3 (bottom), anti-H3K9ac (middle) and either anti-H3K18ac (top left), anti-H3K23ac (top center), or anti-H3K27ac (top right). Lanes: 1 – tailless starting material; 2 – intermediate single H3 tail product; 3 – final asymmetric mono-acetylated / tetra-acetylated nucleosomes.

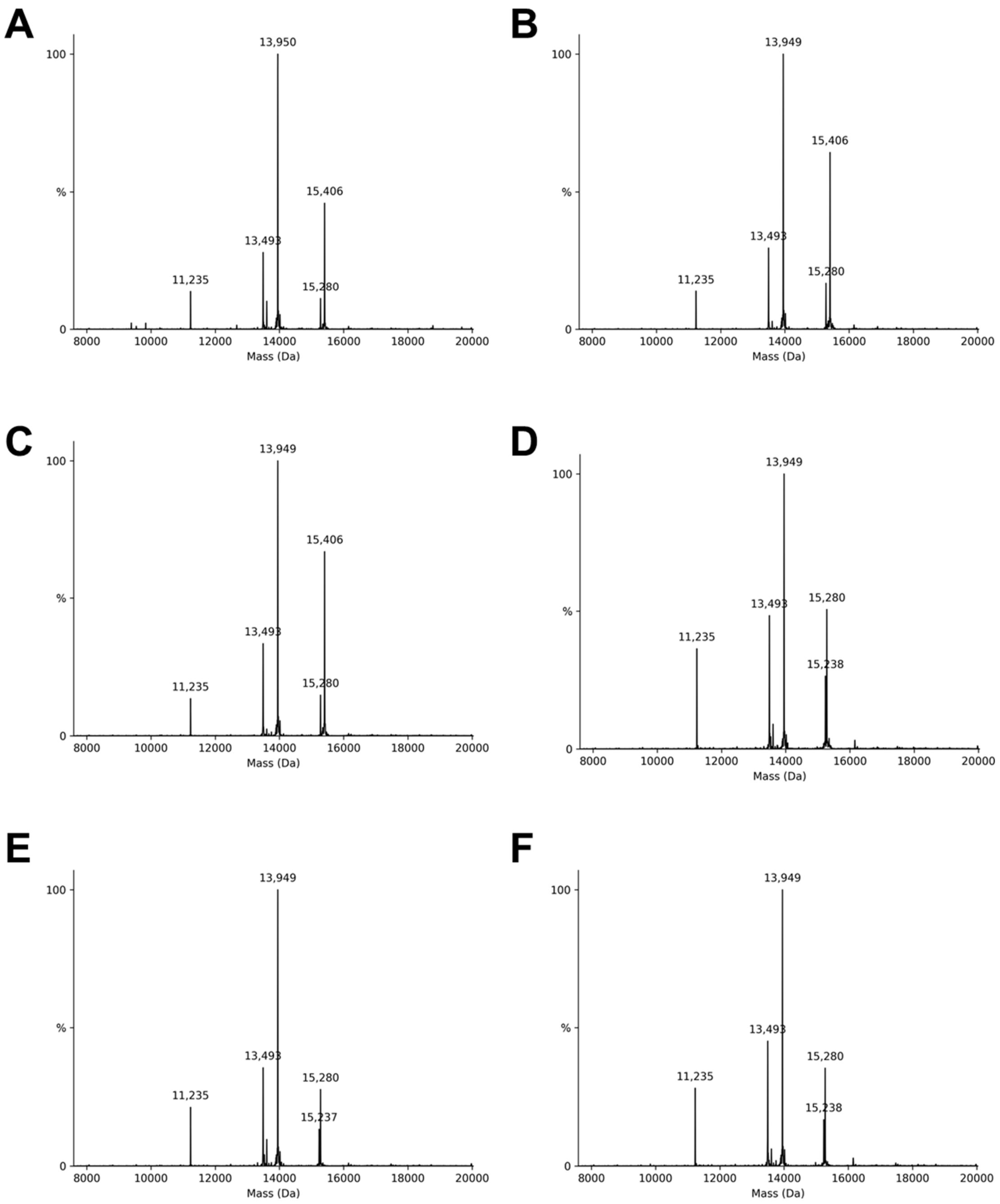

**Figure S33. Mass spectrometric characterization of asymmetric mono/tetra-acetylated nucleosomes and asymmetric mono-acetylated/unmodified nucleosomes.** Deconvoluted mass spectra of (**A**) asymmetric H3K18ac & H3K9/14/23/27ac nucleosome, (**B**) asymmetric H3K23ac & H3K9/14/18/27ac nucleosome, and (**C**) asymmetric H3K27ac & H3K9/14/18/23ac nucleosome: Mono-acetyl H3 calc'd for $C_{672}H_{1133}N_{215}O_{187}S_2$ $[M]^+$: 15280.64 Da, Found: 15280 Da; Tetra-acetyl H3 calc'd for $C_{678}H_{1139}N_{215}O_{190}S_2$ $[M]^+$: 15406.75 Da, Found: 15406 Da. Deconvoluted mass spectra of (**D**) asymmetric H3K18ac & unmodified H3 nucleosome, (**E**) asymmetric H3K27ac & unmodified H3 nucleosome, and (**F**) asymmetric H3K23ac & unmodified H3 nucleosome: Mono-acetyl H3 calc'd for $C_{672}H_{1133}N_{215}O_{187}S_2$ [M]+: 15280.64 Da, Found: 15280 Da; unmodified H3 Calc'd for $C_{670}H_{1131}N_{215}O_{186}S_2$ $[M]^+$: 15238.61 Da, Found: 15238 Da. Raw spectra deconvoluted with UniDec.[5]

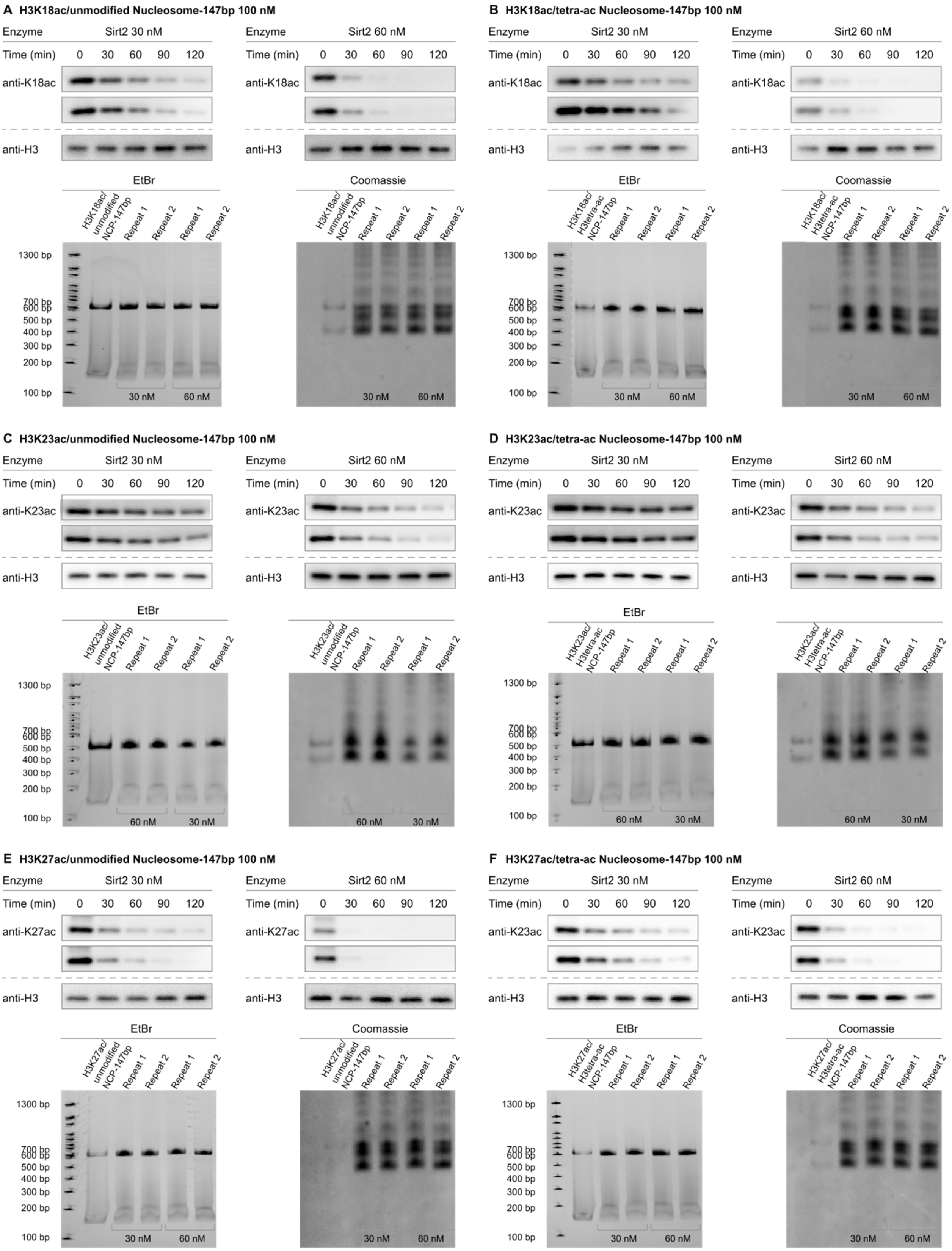
A H3K18ac/unmodified Nucleosome-147bp 100 nM
Enzyme Sirt2 30 nM
Time (min) 0 30 60 90 120
anti-K18ac
anti-H3
Enzyme Sirt2 60 nM
Time (min) 0 30 60 90 120
anti-K18ac
anti-H3
EtBr
H3K18ac/ unmodified NCP-147bp
Repeat 1
Repeat 2
Repeat 1
Repeat 2
1300 bp
700 bp
600 bp
500 bp
400 bp
300 bp
200 bp
100 bp
30 nM
60 nM
Coomassie
B H3K18ac/tetra-ac Nucleosome-147bp 100 nM
Enzyme Sirt2 30 nM
Time (min) 0 30 60 90 120
anti-K18ac
anti-H3
Enzyme Sirt2 60 nM
anti-K18ac
anti-H3
EtBr
H3K18ac/ H3tetra-ac NCP-147bp
Coomassie
C H3K23ac/unmodified Nucleosome-147bp 100 nM
Enzyme Sirt2 30 nM
anti-K23ac
anti-H3
Enzyme Sirt2 60 nM
anti-K23ac
anti-H3
EtBr
H3K23ac/ unmodified NCP-147bp
60 nM
30 nM
D H3K23ac/tetra-ac Nucleosome-147bp 100 nM
Enzyme Sirt2 30 nM
anti-K23ac
anti-H3
Enzyme Sirt2 60 nM
anti-K23ac
anti-H3
EtBr
H3K23ac/ H3tetra-ac NCP-147bp
E H3K27ac/unmodified Nucleosome-147bp 100 nM
Enzyme Sirt2 30 nM
anti-K27ac
anti-H3
Enzyme Sirt2 60 nM
anti-K27ac
anti-H3
EtBr
H3K27ac/ unmodified NCP-147bp
Coomassie
F H3K27ac/tetra-ac Nucleosome-147bp 100 nM
Enzyme Sirt2 30 nM
anti-K23ac
anti-H3
Enzyme Sirt2 60 nM
anti-K23ac
anti-H3
EtBr
H3K27ac/ H3tetra-ac NCP-147bp
Coomassie

**Figure S34. Western blot measurement of Sirt2 activity toward asymmetrically acetylated nucleosomes.** (**A**) Western blots (top; anti-H3K18ac) and native TBE gels (bottom) of Sirt2 deacetylation assays with 147 bp asymmetric unmodified H3 & H3K18ac nucleosomes. (**B**) Western blots (top; anti-H3K18ac) and native TBE gels (bottom) of Sirt2 deacetylation assays with 147 bp asymmetric H3K9ac/K14ac/K23ac/K27ac (H3tetra-ac) & H3K18ac nucleosomes. (**C**) Western blots (top; anti-H3K23ac) and native TBE gels (bottom) of Sirt2 deacetylation assays with 147 bp asymmetric unmodified H3 & H3K23ac nucleosomes. (**D**) Western blots (top; anti-H3K23ac) and native TBE gels (bottom) of Sirt2 deacetylation assays with 147 bp asymmetric H3K9ac/K14ac/K18ac/K27ac (H3tetra-ac) & H3K23ac nucleosomes. (**E**) Western blots (top; anti-H3K27ac) and native TBE gels (bottom) of Sirt2 deacetylation assays with 147 bp asymmetric unmodified H3 & H3K27ac nucleosomes. (**F**) Western blots (top; anti-H3K27ac) and native TBE gels (bottom) of Sirt2 deacetylation assays with 147 bp asymmetric H3K9ac/K14ac/K18ac/K23ac (H3tetra-ac) & H3K27ac nucleosomes.

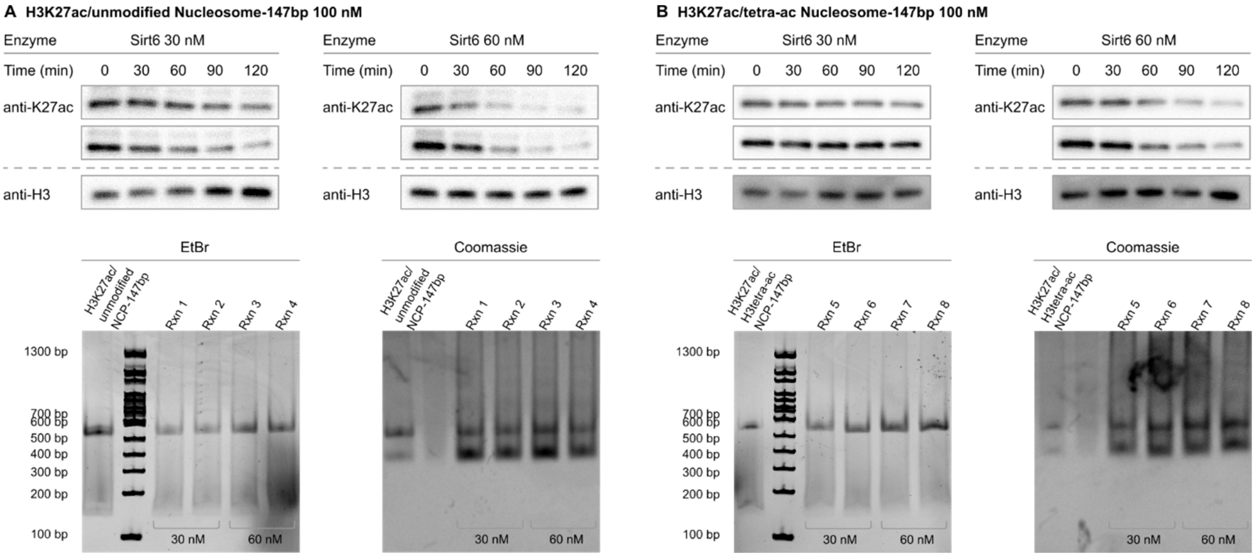

**Figure S35. Western blot measurement of Sirt6 activity toward asymmetrically acetylated nucleosomes.** (**A**) Western blots (top; anti-H3K27ac) and native TBE gels (bottom) of Sirt6 deacetylation assays with 147 bp asymmetric unmodified H3 & H3K27ac nucleosomes. (**B**) Western blots (top; anti-H3K27ac) and native TBE gels (bottom) of Sirt6 deacetylation assays with 147 bp asymmetric H3K9ac/K14ac/K18ac/K23ac (H3tetra-ac) & H3K27ac nucleosomes.

| Sirt2 | | | Sirt2 | | |
|---|---|---|---|---|---|
| **Symmetric Nucleosome V/[E] (min-1)** | **Mono-acetylated** | **Penta-acetylated** | **Asymmetric Nucleosome V/[E] (min-1)** | **mono-acetylated / unmodified** | **mono-acetylated / tetra-acetylated** |
| H3K9ac | 0.038±0.0021* | 0.031±0.0024 | H3K9ac | (-) | (-) |
| H3K14ac | 0.0034±0.0003 | 0.0051±0.0005 | H3K14ac | (-) | (-) |
| H3K18ac | 0.056±0.0028 | 0.11±0.0096 | H3K18ac | 0.058±0.0025 | 0.043±0.0033 |
| H3K23ac | 0.031±0.0011 | 0.056±0.0033 | H3K23ac | 0.034±0.0016 | 0.027±0.0014 |
| H3K27ac | 0.12±0.0071* | 0.051±0.0038 | H3K27ac | 0.12±0.0064 | 0.071±0.0021 |

**Table S9. Calculated V/[E] values for Sirt2 activity toward mono-, penta-, and asymmetrically acetylated nucleosomes.** Values marked with asterisks are re-printed from prior reports.[6]

| Sirt6 | | | Sirt6 | | |
|---|---|---|---|---|---|
| Symmetric Nucleosome V/[E] (min-1) | Mono-acetylated | Penta-acetylated | Asymmetric Nucleosome V/[E] (min-1) | mono-acetylated / unmodified | mono-acetylated / tetra-acetylated |
| H3K9ac | 0.075±0.0044 | 0.072±0.0085 | H3K9ac | (-) | (-) |
| H3K14ac | 0.00080±0.00024 | 0.0013±0.00023 | H3K14ac | (-) | (-) |
| H3K18ac | 0.043±0.0031* | 0.038±0.0033 | H3K18ac | (-) | (-) |
| H3K23ac | 0.011±0.00059* | 0.014±0.0011 | H3K23ac | (-) | (-) |
| H3K27ac | 0.033±0.0029 | 0.019±0.0016 | H3K27ac | 0.036±0.0021 | 0.020±0.0018 |

**Table S10. Calculated V/[E] values for Sirt6 activity toward mono-, penta-, and asymmetrically acetylated nucleosomes.** Values marked with asterisks are re-printed from prior reports.[7]

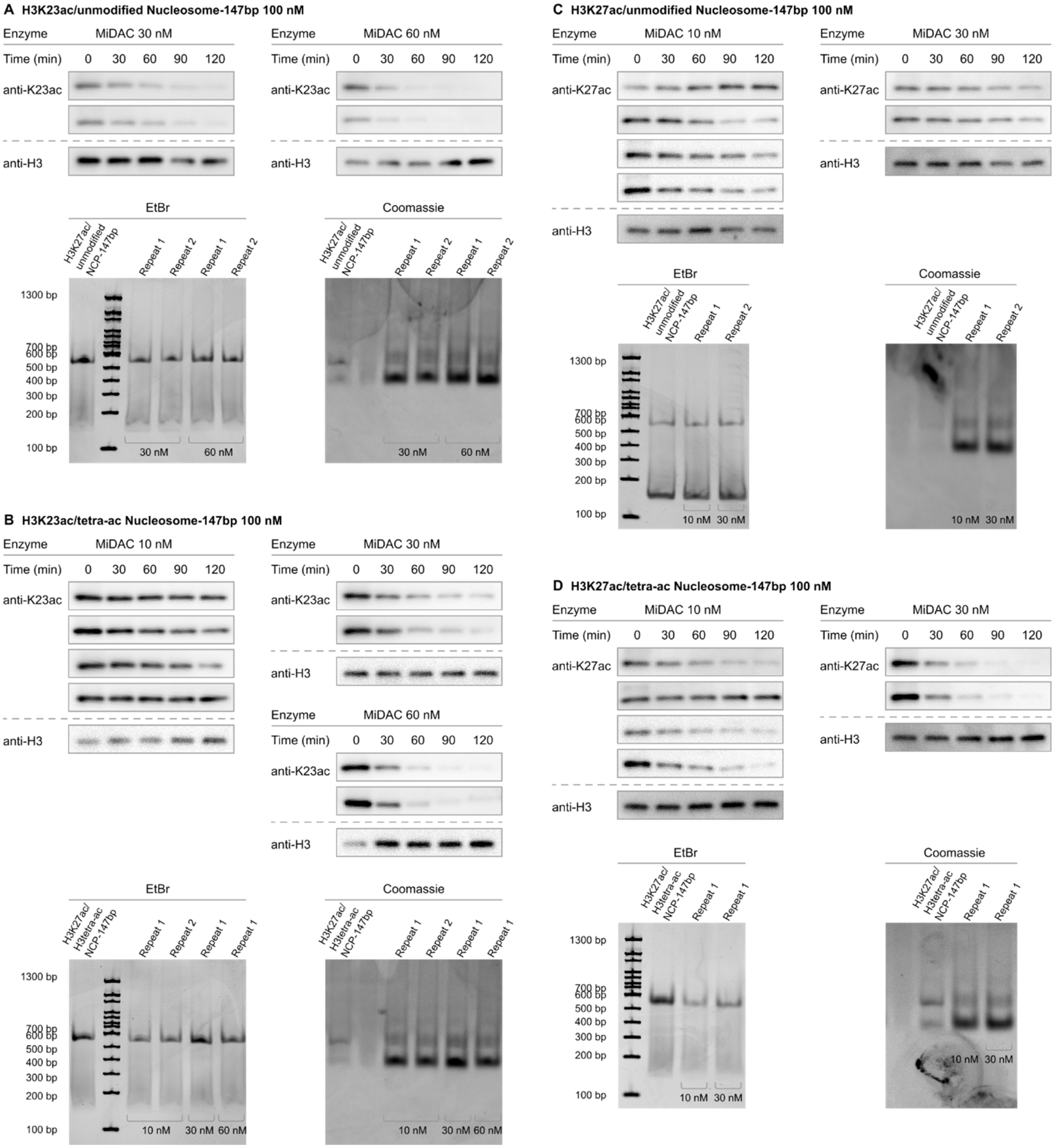

**Figure S36. Western blot measurement of MiDAC activity toward asymmetrically acetylated nucleosomes.** (**A**) Western blots (top; anti-H3K23ac) and native TBE gels (bottom) of MiDAC deacetylation assays with 147 bp asymmetric unmodified H3 & H3K23ac nucleosomes. (**B**) Western blots (top; anti-H3K23ac) and native TBE gels (bottom) of MiDAC deacetylation assays with 147 bp asymmetric H3K9ac/K14ac/K18ac/K27ac (H3tetra-ac) & H3K23ac nucleosomes. (**C**) Western blots (top; anti-H3K27ac) and native TBE gels (bottom) of MiDAC deacetylation assays with 147 bp asymmetric unmodified H3 & H3K27ac nucleosomes. (**D**) Western blots (top; anti-H3K27ac) and native TBE gels (bottom) of MiDAC deacetylation assays with 147 bp asymmetric H3K9ac/K14ac/K18ac/K23ac (H3tetra-ac) & H3K27ac nucleosomes.

| MiDAC | | | MiDAC | | |
|---|---|---|---|---|---|
| Symmetric Nucleosome V/[E] (min-1) | Mono-acetylated | Penta-acetylated | Asymmetric Nucleosome V/[E] (min-1) | mono-acetylated / unmodified | mono-acetylated / tetra-acetylated |
| H3K9ac | 1.4±0.046 | 1.4±0.058 | H3K9ac | (-) | (-) |
| H3K14ac | 0.52 ± 0.034* | 0.39±0.020 | H3K14ac | (-) | (-) |
| H3K18ac | 0.17 ± 0.0051* | 0.18±0.013 | H3K18ac | (-) | (-) |
| H3K23ac | 0.021±0.00063 | 0.11±0.0082 | H3K23ac | 0.065±0.0022 | 0.073±0.0025 |
| H3K27ac | 0.11±0.0094 | 0.40±0.034 | H3K27ac | 0.030±0.00073 | 0.12±0.0017 |

**Table S11. Calculated V/[E] values for MiDAC activity toward mono-, penta-, and asymmetrically acetylated nucleosomes.** Values marked with asterisks are re-printed from prior reports.[8,9]

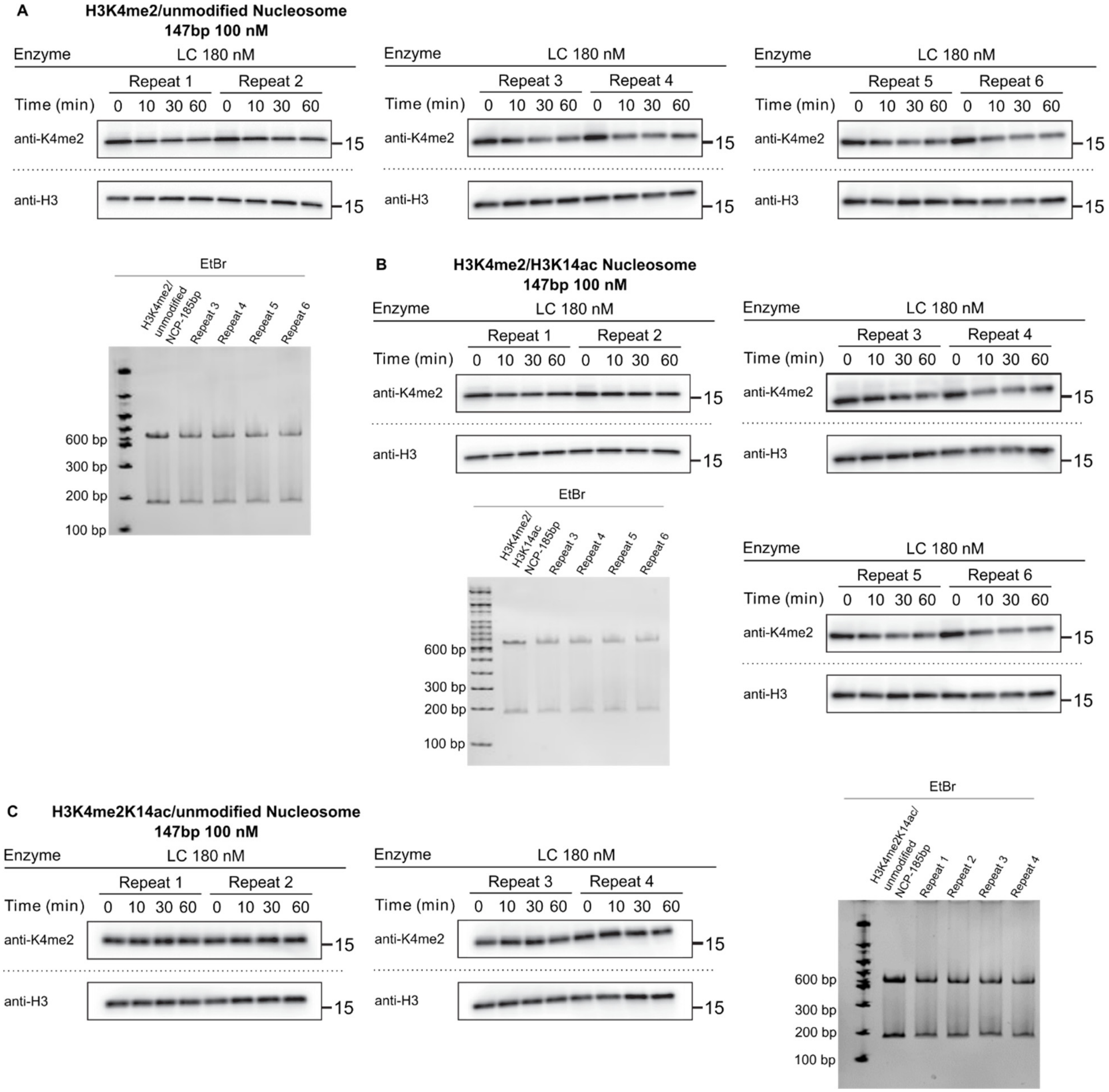

**Figure S37. Western blot measurement of LC activity toward nucleosomes with asymmetric methylation and acetylation.** (**A**) Western blots (top; anti-H3K4me2) and representative TBE native gel (left) of LC demethylation assays with 185 bp asymmetric unmodified H3 & H3K4me2 nucleosomes. (**B**) Western blots (top, right; anti-H3K4me2) and representative TBE native gel (center) of LC demethylation assays with 185 bp asymmetric unmodified H3K14ac & H3K4me2 nucleosomes. (**C**) Western blots (bottom; anti-H3K4me2) and representative TBE native gel (right) of LC demethylation assays with 185 bp asymmetric unmodified H3 & H3K4me2K14ac nucleosomes.

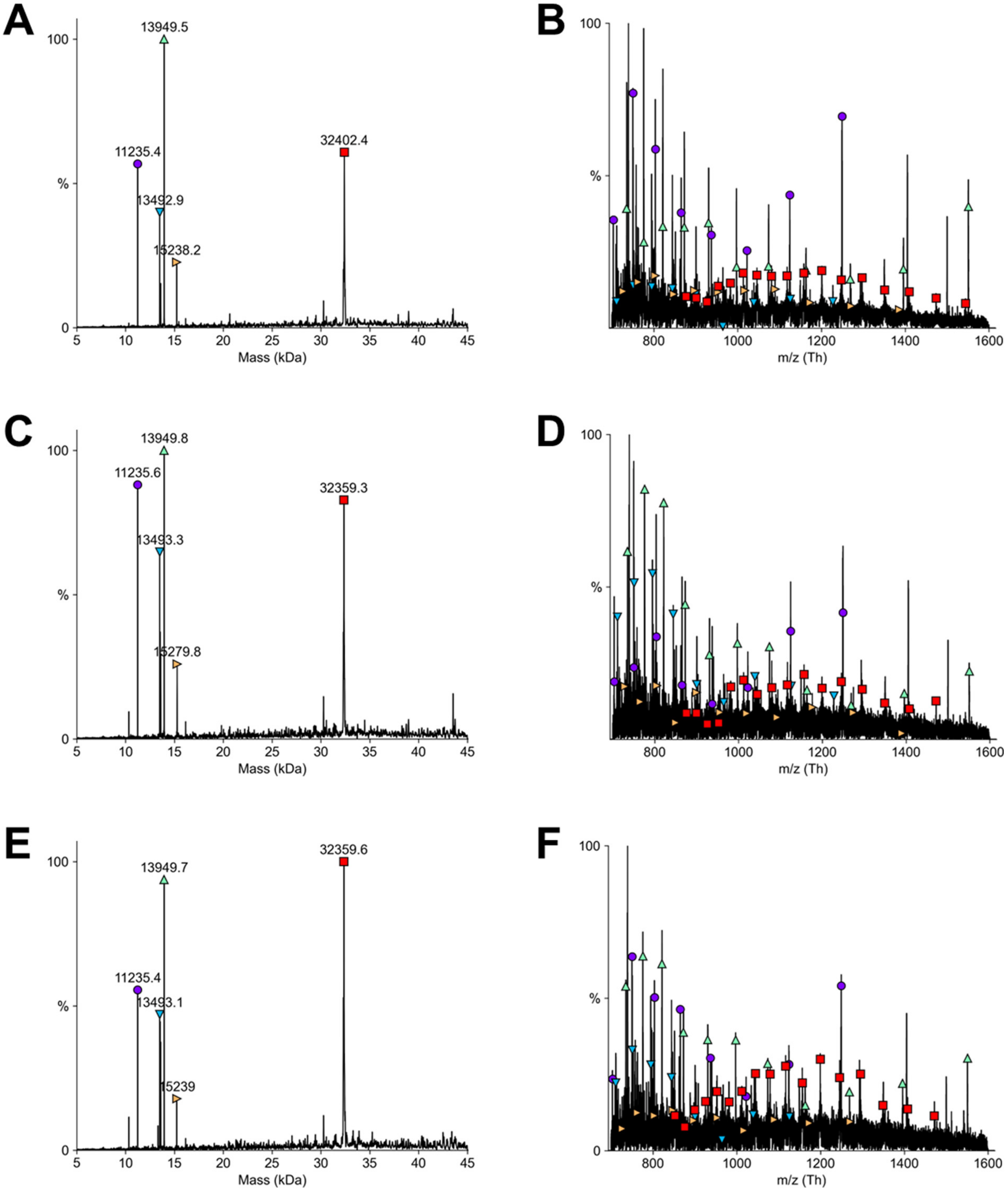

**Figure S38. Mass spectrometric characterization of ubiquitinated H3 peptides and ubiquitinated nucleosome.** Deconvoluted (**A**) and raw (**B**) mass spectra of asymmetric 185 bp H3K9me3/K18Ub(G76A)/K23Ub(G76A) & unmodified H3 nucleosome: H4 (purple circle) calculated mass 11236.15 Da, found: 11235.4 Da; H2B (teal downward pointed triangle) calculated mass 13493.68 Da, found: 13493.1 Da; H2A (green upward pointed triangle) calculated mass 13950.2 Da, found: 13949.7 Da; H3 (orange rightward pointed triangle) calculated mass 15238.61 Da, found: 15239.0 Da; H3K9me3/K18Ub(G76A)/K23Ub(G76A) (red square) calculated mass 32402.18 Da, found: 32402.4 Da. Deconvoluted (**C**) and raw (**D**) mass spectra of asymmetric 185 bp H3K18Ub(G76A)/K23Ub(G76A) &

H3K9me3 nucleosome: H4 (purple circle) calculated mass 11236.15 Da, found: 11235.4 Da; H2B (teal downward pointed triangle) calculated mass 13493.68 Da, found: 13493.3 Da; H2A (green upward pointed triangle) calculated mass 13950.2 Da, found: 13949.8 Da; H3K9me3 (orange rightward pointed triangle) calculated mass 15279.68 Da, found: 15279.8 Da; H3K18Ub(G76A)/K23Ub(G76A) (red square) calculated mass 32360.51 Da, found: 32359.3 Da. Deconvoluted (**E**) and raw (**F**) mass spectra of asymmetric 185 bp H3K18Ub(G76A)/K23Ub(G76A) & unmodified H3 nucleosome: H4 (purple circle) calculated mass 11236.15 Da, found: 11235.4 Da; H2B (teal downward pointed triangle) calculated mass 13493.68 Da, found: 13493.1 Da; H2A (green upward pointed triangle) calculated mass 13950.2 Da, found: 13949.7 Da; H3 (orange rightward pointed triangle) calculated mass 15238.61 Da, found: 15239.0 Da; H3K18Ub(G76A)/K23Ub(G76A) (red square) calculated mass 32360.51 Da, found: 32359.6 Da. Raw spectra deconvoluted with UniDec.[5]

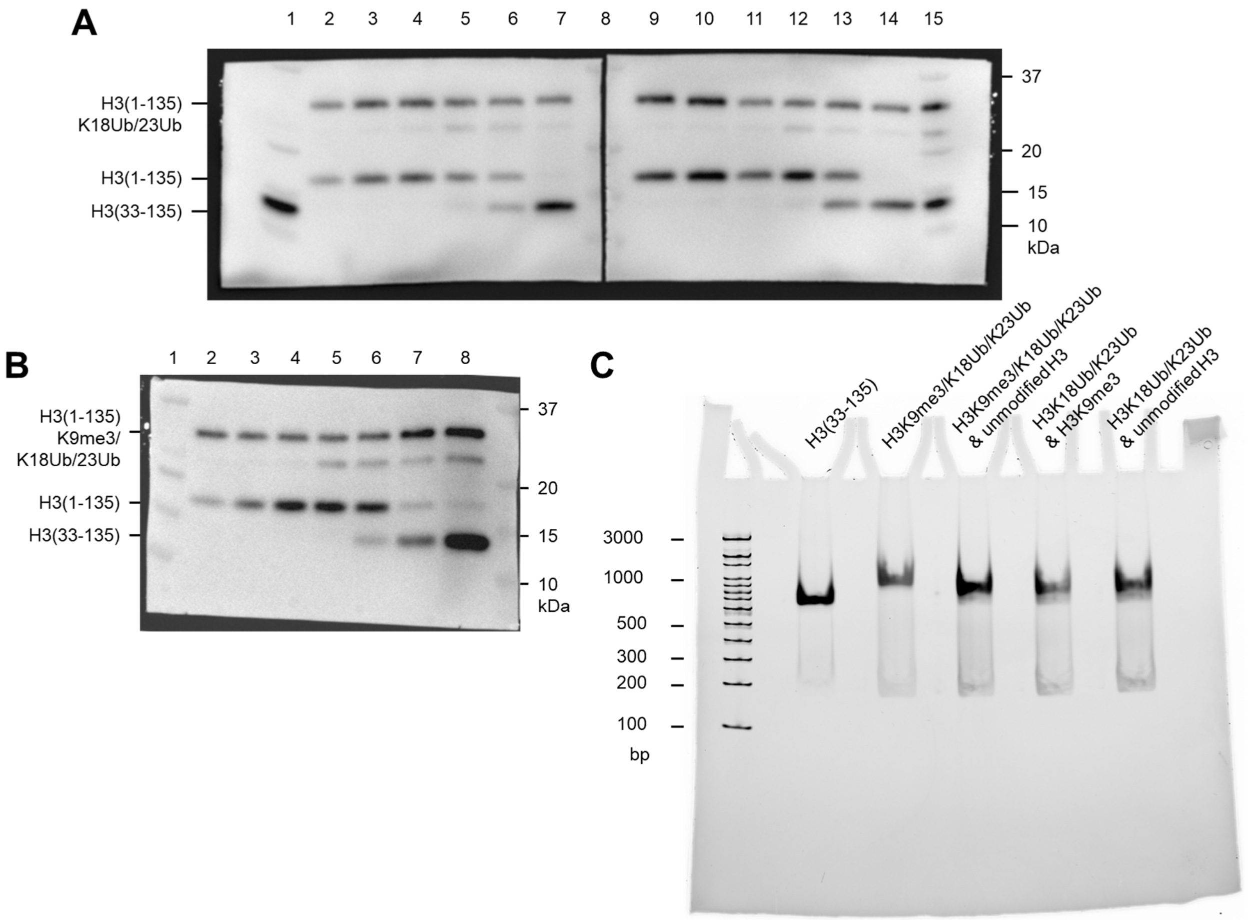

**Figure S39. Characterization of asymmetric unmodified/ubiquitinated nucleosome.** (**A**) Anti-H3 blot of tailless nucleosome starting material, intermediate single tail ligation product, asymmetrically modified products, and intermediate fractions from weak anion exchange purification.
Lanes: 1 – tailless starting material & ladder; 2 –final asymmetric H3K18Ub/K23Ub and unmodified H3 nucleosomes; 3-6 – impure weak anion exchange fractions in elution order ; 7 – intermediate single H3 tail product; 8 – ladder; 9 –final asymmetric H3K18Ub/K23Ub and H3K9me3 nucleosomes; 10-14 – impure weak anion exchange fractions in elution order 14 – intermediate single H3 tail product; 15 – ladder. (**B**) Anti-H3 blot of tailless nucleosome starting material, intermediate single tail ligation product, asymmetrically modified products, and intermediate fractions from weak anion exchange purification.
Lanes: 1 –ladder; 2 –final asymmetric H3K9me3/K18Ub/K23Ub and unmodified H3 nucleosomes; 3-8 – impure weak anion exchange fractions in elution order. (**C**) TBE native gel of 185 bp starting material symmetric ubiquitinated products of the cW11 sortase ligation, and asymmetric ubiquitinated products of the cW11 sortase ligation.

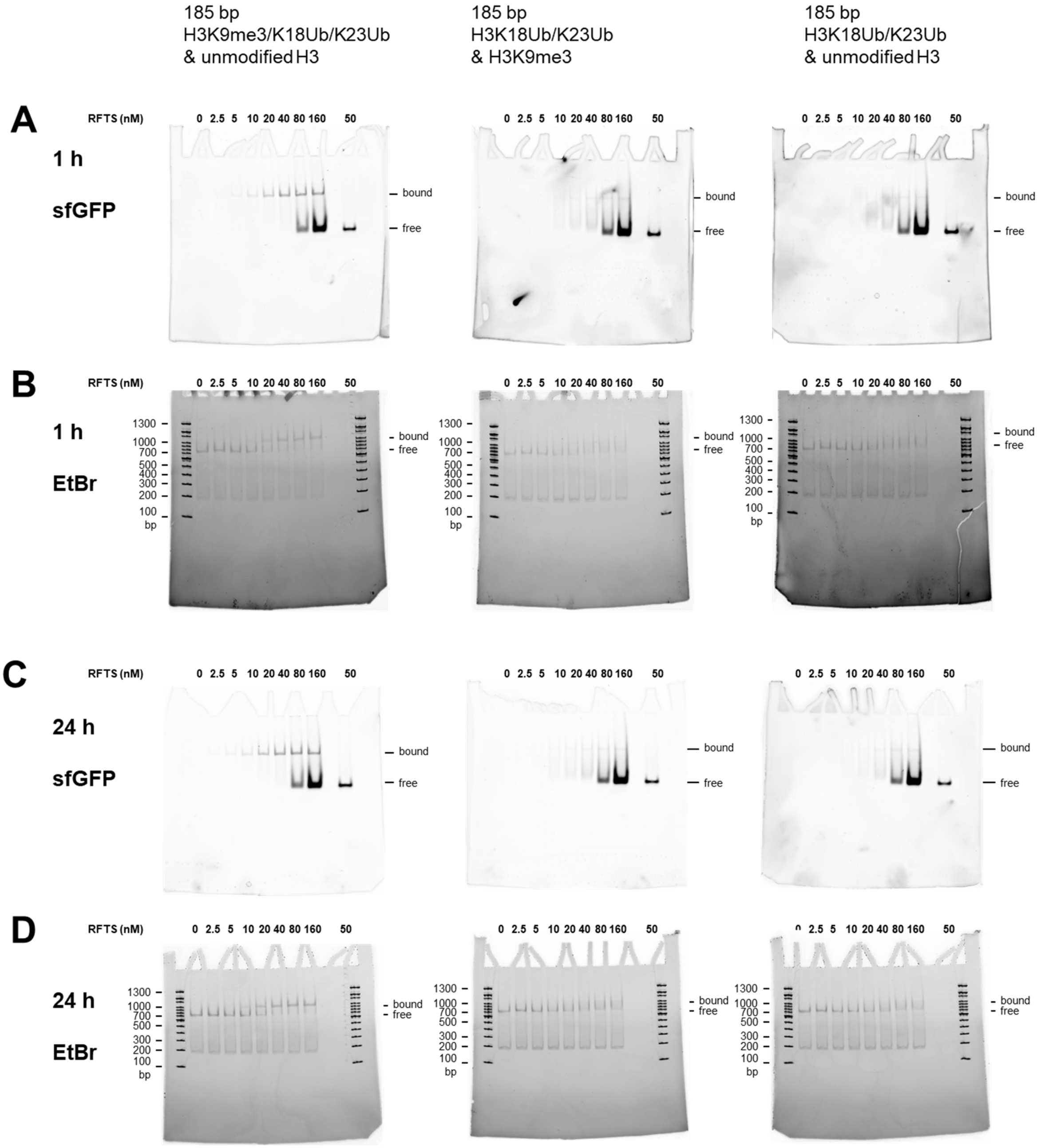

**Figure S40. Electrophoretic mobility shift assay titrating asymmetric nucleosomes with sfGFP-RFTS fusion.** (**A**) Fluorescence visualization of sfGFP-RFTS fusion (titrant) binding to 185 bp nucleosomes with asymmetric H3K9me3/K18Ub/K23Ub and unmodified H3 (left), asymmetric K18Ub/K23Ub and H3K9me3 (middle), or asymmetric K18Ub/K23Ub and unmodified H3 (right) after 1 hr at 4 °C. (**B**) Ethidium bromide (EtBr) visualization of sfGFP-RFTS fusion (titrant) binding to 185 bp nucleosomes with asymmetric H3K9me3/K18Ub/K23Ub and unmodified H3 (left), asymmetric K18Ub/K23Ub and H3K9me3 (middle), or asymmetric K18Ub/K23Ub and unmodified H3 (right) after 1 hr at 4 °C. (**A**) Fluorescence visualization of sfGFP-RFTS fusion (titrant) binding to 185 bp nucleosomes with asymmetric H3K9me3/K18Ub/K23Ub and unmodified H3 (left), asymmetric K18Ub/K23Ub and H3K9me3 (middle), or asymmetric K18Ub/K23Ub and unmodified H3 (right) after 24 hr at 4 °C. (**B**) Ethidium bromide (EtBr) visualization of sfGFP-RFTS fusion (titrant) binding to 185 bp nucleosomes with asymmetric H3K9me3/K18Ub/K23Ub and unmodified H3 (left), asymmetric K18Ub/K23Ub and H3K9me3 (middle), or asymmetric K18Ub/K23Ub and unmodified H3 (right) after 24 hr at 4 °C.

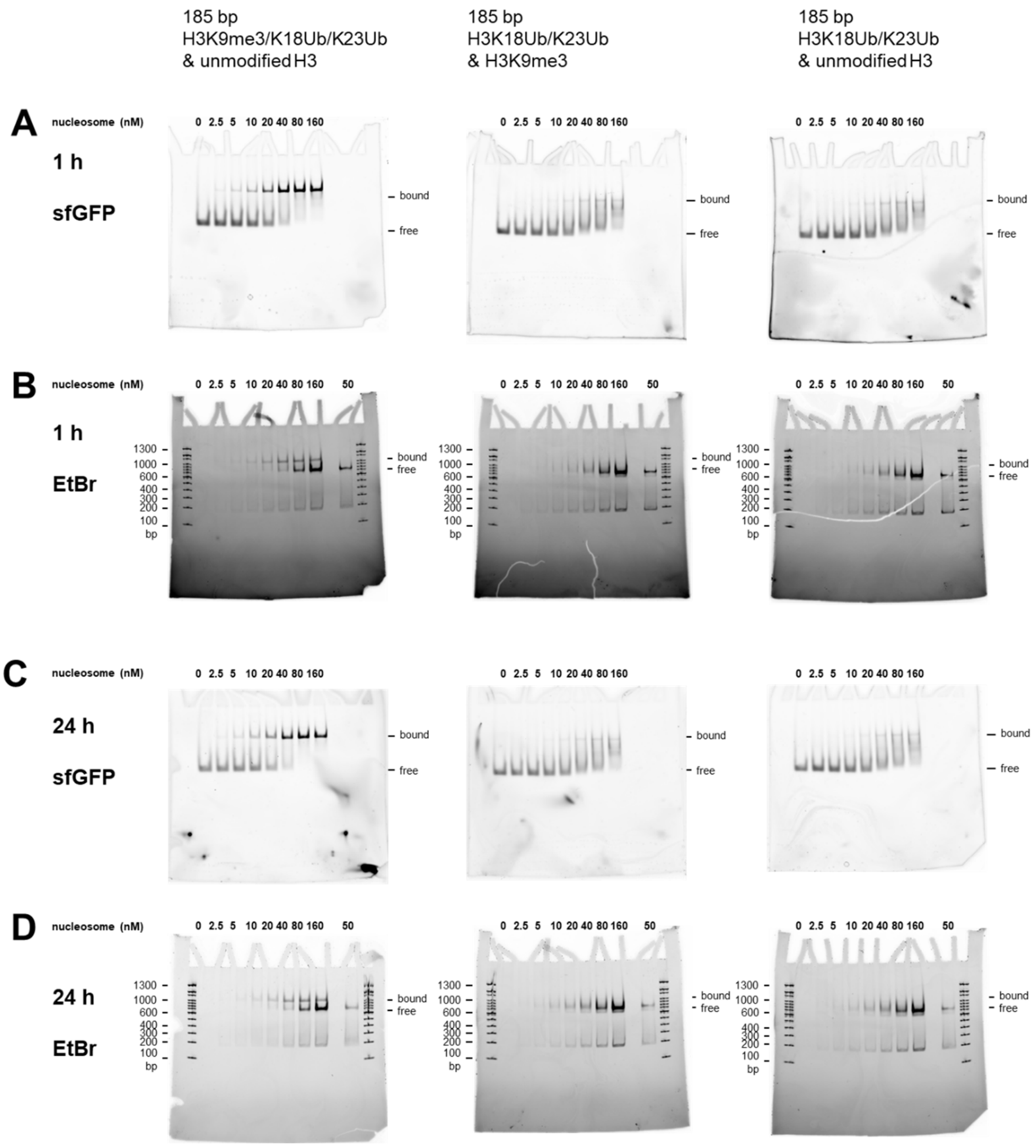

**Figure S41. Electrophoretic mobility shift assay titrating sfGFP-RFTS fusion with asymmetric nucleosomes.** (**A**) Fluorescence visualization of sfGFP-RFTS fusion binding to 185 bp nucleosomes (titrant) with asymmetric H3K9me3/K18Ub/K23Ub and unmodified H3 (left), asymmetric K18Ub/K23Ub and H3K9me3 (middle), or asymmetric K18Ub/K23Ub and unmodified H3 (right) after 1 hr at 4 °C. (**B**) Ethidium bromide (EtBr) visualization of sfGFP-RFTS fusion binding to 185 bp nucleosomes (titrant) with asymmetric H3K9me3/K18Ub/K23Ub and unmodified H3 (left), asymmetric K18Ub/K23Ub and H3K9me3 (middle), or asymmetric K18Ub/K23Ub and unmodified H3 (right) after 1 hr at 4 °C. (**A**) Fluorescence visualization of sfGFP-RFTS fusion binding to 185 bp nucleosomes (titrant) with asymmetric H3K9me3/K18Ub/K23Ub and unmodified H3 (left), asymmetric K18Ub/K23Ub and H3K9me3 (middle), or asymmetric K18Ub/K23Ub and unmodified H3 (right) after 24 hr at 4 °C. (**B**) Ethidium bromide (EtBr) visualization of sfGFP-RFTS fusion binding to 185 bp nucleosomes (titrant) with asymmetric H3K9me3/K18Ub/K23Ub and unmodified H3 (left), asymmetric K18Ub/K23Ub and H3K9me3 (middle), or asymmetric K18Ub/K23Ub and unmodified H3 (right) after 24 hr at 4 °C.

**References:**

1. Piotukh, K. *et al.* Directed Evolution of Sortase A Mutants with Altered Substrate Selectivity Profiles. *J Am Chem Soc* **133**, 17536–17539 (2011).
2. Chen, I., Dorr, B. M. & Liu, D. R. A general strategy for the evolution of bond-forming enzymes using yeast display. *Proc Natl Acad Sci U S A* **108**, 11399–11404 (2011).
3. Zhulenkovs, D., Jaudzems, K., Zajakina, A. & Leonchiks, A. Enzymatic activity of circular sortase A under denaturing conditions: An advanced tool for protein ligation. *Biochem Eng J* **82**, 200–209 (2014).
4. Musil, M. *et al.* FireProt: web server for automated design of thermostable proteins. *Nucleic Acids Res* **45**, W393–W399 (2017).
5. T. Marty, M. *et al.* Bayesian Deconvolution of Mass and Ion Mobility Spectra: From Binary Interactions to Polydisperse Ensembles. *Anal Chem* **87**, 4370–4376 (2015).
6. Abeywardana, M. Y. *et al.* Multifaceted regulation of Sirtuin 2 (Sirt2) Deacetylase Activity. *Journal of Biological Chemistry* **0**, 107722 (2024).
7. Wang, Z. *et al.* Structural Basis of Sirtuin 6-Catalyzed Nucleosome Deacetylation. *J Am Chem Soc* **145**, 6811–6822 (2023).
8. Wang, Z. A. *et al.* Diverse nucleosome site-selectivity among histone deacetylase complexes. *eLife* **9**, e57663 (2020).
9. Wang, Z. A. *et al.* Histone H2B Deacylation Selectivity: Exploring Chromatin's Dark Matter with an Engineered Sortase. *J Am Chem Soc* **144**, 3360–3364 (2022).
